# Multi-omics analysis of type II diabetic wound healing reveals CD44-mediated immune cell crosstalk dysfunction in mice and humans

**DOI:** 10.64898/2026.04.26.720829

**Authors:** Mateusz S. Wietecha, Jingbo Pang, Miya Kang, Avin Hafedi, Samantha Walsdorf, Shalyn Keiser, Mark Maienschein-Cline, Timothy J. Koh

## Abstract

Type II diabetes mellitus (T2DM) is one of the most prevalent diseases in the United States and is associated with diabetic foot ulcers (DFU) and their impaired, often chronic, wound healing. The T2DM mouse model with dysfunctional leptin receptor (db/db) has been used in basic and translational studies of wound healing due to its systemic phenotypes (hyperphagia, hypometabolism, obesity, T2DM) and its notable delayed skin wound healing. However, a characterization of the temporal cellular dynamics of the db/db wound healing model has not been performed, nor has the model been systematically compared to human DFUs. We performed the first comprehensive single-cell, multi-omic analysis of dermal cells in diabetic (db/db) compared to non-diabetic (ND) mice across three time points ranging from the inflammatory to the delayed proliferative and resolution phases of healing. Single-cell transcriptomics were uniquely linked to their corresponding cells’ surface protein expressions of cell-specific receptors, including immune cells (CD45) such as neutrophils (CD11b, Ly6G), monocytes/macrophages (CD11b, F4/80, CD11c, Ly6C) and T lymphocytes (CD3, CD4), and dermal cells such as endothelial cells (CD31) and fibroblasts (CD26, CD140a), and showed high concordance between protein cell markers and their gene expressions in major cell types. Differential multi-omic analyses characterized two neutrophil (*Tnfaip3^+^Sod2^+^*Ly6G^+^, *Csf3r^+^Fos^+^*Ly6G^+^), three monocyte/macrophage (F4/80^high^CD11b^high^, Ly6c^high^CD11b^high^, CD11c^high^CD11b^low^) and three fibroblast (*Pi16^+^Dpp4^+^*CD26^high^, *Lrrc15^+^Tnc^+^*CD140a^high^, *Cilp^+^Mgp^+^*CD26^low^) subtypes showing dysregulated dynamics across the time course of healing in db/db vs ND mice. Notably, NETotic *Tnfaip3^+^Sod2^+^*Ly6G^+^ neutrophils and phagocytic F4/80^high^CD11b^high^ macrophage subtypes were drastically up-regulated in diabetic wounds.

Differential cell-cell communication analyses revealed striking differences in crosstalk dynamics between fibroblast, macrophage and neutrophil subtypes in the early phase of healing, and ligand-receptor interactome analyses identified CD44 as the hub of dysregulated immune cell interactions in diabetic wounds, implicating cell adhesion, migration and inflammatory pathways, especially those mediated by ICAM1. Inhibition of CD44 using blocking antibodies in primary macrophages from db/db mice and via intradermal injections in db/db mice significantly normalized the early wound immune dysfunction, in part by inhibiting ICAM1 and reversing the excessive neutrophil influx into diabetic wounds. A new integrated dataset of single-cell human chronic wound studies revealed similar CD44-mediated immune cell dysfunctions in diabetic vs non-diabetic foot ulcers, pointing to CD44 as a promising therapeutic target for T2DM-associated chronic wounds.

## INTRODUCTION

Type II diabetes mellitus (T2DM) is one of the most prevalent diseases in the United States, and it is associated with chronic, non-healing diabetic foot ulcers (DFU), which themselves account for an estimated 5-year mortality rate of 30%, which rises to over 70% after a major amputation ^1,2^. Indeed, each year DFUs affect approximately 18.6 million people worldwide and 1.6 million people in the United States, resulting in a major healthcare burden. During physiological skin wound healing, the precise temporal coordination of cells across inflammatory, proliferative and resolution phases ensures timely wound closure and the re-establishment of structural and metabolic tissue homeostasis ^3,4^. However, in T2DM-affected healing, this healing phase progression is fundamentally derailed, leading to a state of persistent inflammation and impaired repair ^5^. Recent advances in single-cell transcriptomics have begun to map the heterogeneity of the DFU microenvironment, revealing distinct states of fibroblasts, macrophages, neutrophils and other cell types that may drive either repair or chronicity ^6–11^. Despite these insights, the exact molecular hubs that mediate the aberrant crosstalk between cell types in diabetic wounds remain poorly defined.

While the leptin-receptor-deficient (db/db) mouse has served as a major model for studying diabetes-driven delayed wound healing due to its T2DM-like systemic phenotypes ^12–14^, our understanding of its temporal cellular and molecular dynamics remains incomplete. Previous studies have profiled specific cellular aspects of this model, including altered neutrophil and monocyte/macrophage infiltrations ^15–18^. However, a multi-cellular characterization of the db/db mouse model’s healing dynamics and its systematic comparison to human DFUs has been missing, significantly hindering translational research progress.

In this study, we performed the first comprehensive single-cell, multi-omic analysis of myeloid and dermal cells in diabetic (db/db) and non-diabetic (ND) mice across the inflammatory, proliferative, and resolution phases of healing. By uniquely linking surface protein expression to transcriptomic profiles, we identified specific dysregulated cell subtypes, including neutrophils enriched for NETosis signatures and macrophages enriched for oxidative stress and phagocytic signatures that dominated the early diabetic wound bed. Differential cell-cell communication analyses identified CD44 as a central hub of dysregulated immune cell and dermal interactions, and when CD44 was blocked *in vitro* and *in vivo*, the excessive neutrophil infiltration and macrophage dysfunction seen in diabetic wounds were reversed. Comparative analysis with a novel integrated human DFU dataset showed similar immune and dermal cell crosstalk disruptions and confirmed that the CD44 interactome is dysregulated across species. Our findings establish CD44 as a primary driver of the early-phase diabetic wound microenvironment and a promising therapeutic target for chronic wounds.

## METHODS

### Animals and skin wound model

Male C57Bl/6 (Non-diabetic (ND), control) and BKS.Cg-*Dock7^m^* +/+ *Lepr^db^*/J (*db/db*, diabetic) mice were purchased from the Jackson Laboratory (Bar Harbor, ME) and housed in an environmentally controlled animal facility for at least two weeks before experiments. Adult mice (age 10-12 weeks) were subjected to excisional wounding of the shaved dorsal skin with an 8 mm biopsy punch (two wounds per mouse) ^17^. Wounds were covered with Tegaderm (3M,1626W) until tissue harvest.

Prior to tissue collection, mice were anesthetized and perfused with ice-cold DPBS (Corning, Corning, NY, USA) via the left ventricle. Skin wounds were harvested on the designated experimental days post-injury using a 12 mm biopsy punch, including both the wound and surrounding tissue. Tissues were dissociated into single cell suspensions through combined mechanical and enzymatic digestion. Briefly, wound samples were placed in a Petri dish containing 1 mL of an enzyme cocktail composed of collagenase I, collagenase XI, and hyaluronidase (5 mg/mL each, Sigma, St. Louis, MO) with DNase I (25 µg/mL, Sigma). Using forceps, samples were flattened, and the epidermis was carefully scraped from the wound with a Lance scalpel blade (Cincinnati Surgical, Cincinnati, OH). The dermis was then minced and digested in the enzyme cocktail for 45 min at 37 °C. Finally, cells were filtered through a 40 µm cell strainer and resuspended in cold 1× DPBS.

All animal procedures were approved by the Animal care and Use Committee of the University of Illinois at Chicago.

### Total cell population of skin wound single cell RNA and protein sequencing

Skin cells were pooled from four wounds (two wounds from each of two mice per genotype) and collected on days 3, 6, and 10 post-injury, as described above. After washing, red blood cells were lysed and removed from the single cell suspension using 1x Red Blood Cell Lysis Solution (Miltenyi Biotec, Gaithersburg, MD, USA), followed by debris removal with Debris Removal Solution (Miltenyi Biotec) according to the manufacturer’s instructions. Cells were then resuspended in cold 1x DPBS and passed through a 30 µm Pre-Separation Filter (Miltenyi Biotec). After centrifugation, dead cells were removed using the Dead Cell Removal Kit (Miltenyi Biotec) following the manufacturer’s instructions. Briefly, cells were incubated with an optimized volume of Dead Cell Removal MicroBeads at room temperature for 15 min. Subsequently, 1x Binding Buffer was added to a final volume of 500 µL, and cells were passed through an MS MACS column system (Miltenyi Biotec) for a negative magnetic selection, with a total of four separations. Final live cells were washed, counted, and resuspended in cell staining buffer (BioLegend), and processed for surface protein labeling.

Next, cells were incubated with TruStain FcX™ PLUS (anti-mouse CD16/32) antibody (BioLegend) on ice for 10 min to block Fc receptors, followed by staining with the TotalSeq™-A antibody pool (BioLegend) on ice for 30 min according to the manufacturer’s instructions. A complete list of antibodies is provided in **Table 1**. Cells were then washed three times with cell staining buffer, filtered through a Flowmi™ Cell Strainer (40μm, SP Bel-Art, Wayne, NJ, USA), and resuspended in 1x DPBS supplemented with 10% fetal bovine serum. After cell counting and viability assessment (>85% live cells), samples were immediately processed using the Chromium platform (10x Genomics, San Francisco, CA, USA). Gene and surface protein libraries were constructed according to the Chromium GEM-X Single Cell 3 Reagent Kits v4 User Guide (CG000732, 10x Genomics) and the TotalSeq™-A antibodies with 10x Single Cell 3 Reagent Kit v3 protocol (BioLegend), and sequenced on a NovaSeq 6000 Sequencing System (Illumina, San Diego, CA, USA).

**Table 1:** Antibodies used for protein sequencing.

| Name | Clone | Manufacturer |
| --- | --- | --- |
| TotalSeq™-A0014 anti-mouse/human CD11b Antibody | M1/70 | BioLegend |
| TotalSeq™-A0015 anti-mouse Ly-6G Antibody | 1A8 | BioLegend |
| TotalSeq™-A0013 anti-mouse Ly-6C Antibody | HK1.4 | BioLegend |
| TotalSeq™-A0114 anti-mouse F4/80 Antibody | BM8 | BioLegend |
| TotalSeq™-A0106 anti-mouse CD11c Antibody | N418 | BioLegend |
| TotalSeq™-A0118 anti-mouse NK-1.1 Antibody | PK136 | BioLegend |
| TotalSeq™-A0184 anti-mouse CD335 (NKp46) Antibody | 29A1.4 | BioLegend |
| TotalSeq™-A0182 anti-mouse CD3 Antibody | 17A2 | BioLegend |
| TotalSeq™-A0001 anti-mouse CD4 Antibody | RM4-5 | BioLegend |
| TotalSeq™-A0002 anti-mouse CD8a Antibody | 53-6.7 | BioLegend |
| TotalSeq™-A0096 anti-mouse CD45 Antibody | 30-F11 | BioLegend |
| TotalSeq™-A0904 anti-mouse CD31 Antibody | 390 | BioLegend |
| TotalSeq™-A0449 anti-mouse CD326 (Ep-CAM) Antibody | G8.8 | BioLegend |
| TotalSeq™-A0883 anti-mouse CD26 (DPP-4) Antibody | H194-112 | BioLegend |
| TotalSeq™-A0573 anti-mouse CD140a Antibody | APA5 | BioLegend |
| TotalSeq™-A0937 anti-mouse CD202b (Tie-2, CD202) Antibody | TEK4 | BioLegend |
| TotalSeq™-A0012 anti-mouse CD117 (c-kit) Antibody | 2B8 | BioLegend |
| TotalSeq™-A0105 anti-mouse CD115 (CSF-1R) Antibody | AFS98 | BioLegend |

### Bioinformatics analysis setup and package versions

The single-cell RNA sequencing analysis workflow used herein was described in a detailed published protocol ^19^. A complete list of software used for the bioinformatics analyses is provided in **Table 2**.

**Table 2:**
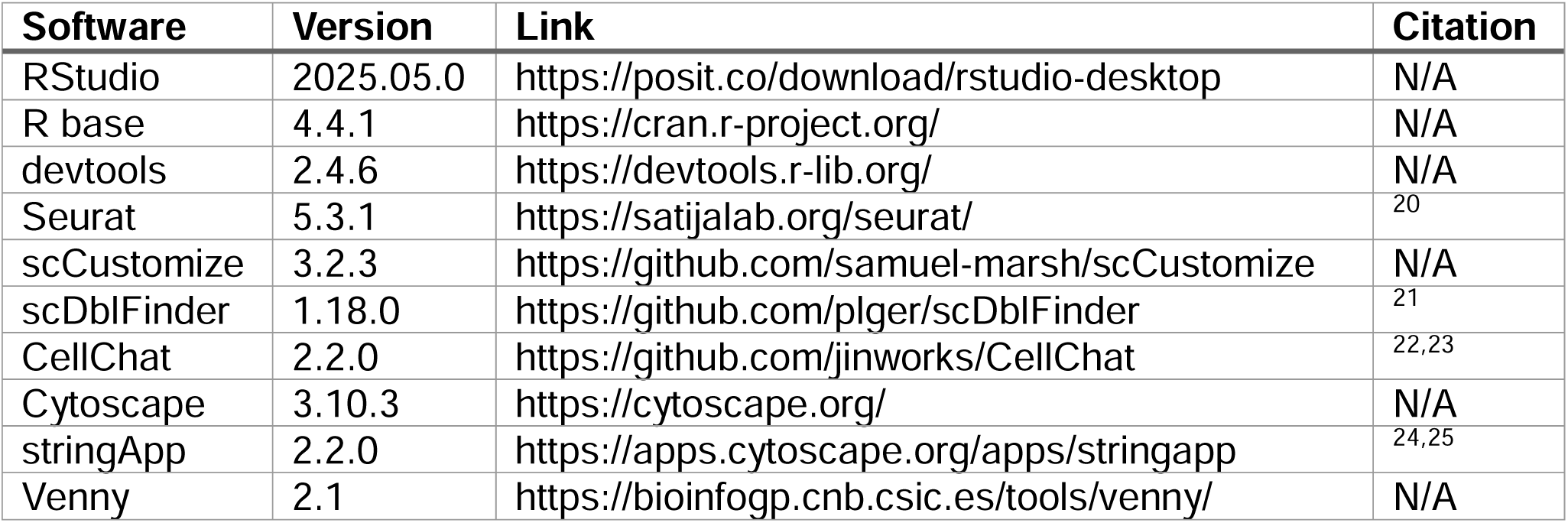
Software used for bioinformatics analyses.

| Software | Version | Link | Citation |
| --- | --- | --- | --- |
| RStudio | 2025.05.0 | https://posit.co/download/rstudio-desktop | N/A |
| R base | 4.4.1 | https://cran.r-project.org/ | N/A |
| devtools | 2.4.6 | https://devtools.r-lib.org/ | N/A |
| Seurat | 5.3.1 | https://satijalab.org/seurat/ | <sup>20</sup> |
| scCustomize | 3.2.3 | https://github.com/samuel-marsh/scCustomize | N/A |
| scDblFinder | 1.18.0 | https://github.com/plger/scDblFinder | <sup>21</sup> |
| CellChat | 2.2.0 | https://github.com/jinworks/CellChat | <sup>22,23</sup> |
| Cytoscape | 3.10.3 | https://cytoscape.org/ | N/A |
| stringApp | 2.2.0 | https://apps.cytoscape.org/apps/stringapp | <sup>24,25</sup> |
| Venny | 2.1 | https://bioinfogp.cnb.csic.es/tools/venny/ | N/A |

### Dataset processing, quality control, merging and protein decomplexing

Raw scRNAseq data from the six samples (ND_3, ND_6, ND_10, db_3, db_6, db_10) was individually de-multiplexed and aligned to reference mouse genome GRCm38 (mm10) using CellRanger software (10x Genomics) and default settings. Counts matrices were exported for analysis using the R package Seurat.

Quality control for each sample was performed as described previously ^19^. The distributions of the UMI counts, unique features and percent mitochondrial genes per cell were visualized via *FeatureScatter* **(Fig S1A)**. High quality cells were selected via the *subset* function by omitting cells with: >10% mitochondrial gene expression, <1000 UMI counts, and <500 unique features. Samples were then combined into a single integrated dataset using the *merge* function.

Integrated data was normalized via *NormalizeData*, scaled via *ScaleData*, and the top 5000 most variably expressed genes (via *FindVariableFeatures*) were used for unsupervised principal component (PC) analysis via *RunPCA* for the top 200 PCs. PCA p-values were computed for PCs using *JackStraw*, the max deviation statistic from Kolmogorov-Smirnov test was computed from the JackStraw statistics and visualized using *ElbowPlot* **(Fig S1B)**. The top PCs that had max deviation > 0.1 were selected, resulting in 131 PCs.

Clustering was performed via *FindNeighbors* using ndims = 131, followed by *FindClusters* using Louvain clustering with resolution = 0.1. Dimensionality reduction via UMAP was performed via *RunUMAP* with ndims = 131, resulting in the final UMAP embedding with 12 Seurat-determined clusters used for downstream analysis **(Fig S1C)**. Individual samples within the merged dataset were visualized via *DimPlot*, which showed no batch effects between samples, and therefore no additional batch correction nor algorithmic integration was necessary **(Fig S1D)**.

Protein data were de-complexed using *HTODemux* as described previously ^19,26^. The first and second top scoring protein scores per cell were identified using the maxID and secondID annotations and visualized via *DimPlot* **(Fig S1E).***Cell typing according to protein and RNA markers*

The correlations of expressions of antibody-tagged proteins and their corresponding genes were visualized in the overall dataset by *DotPlot* and grouping of cells according to their protein maxID (left) and secondID (right) annotations **(Fig S2A)**. This revealed the following antibodies were the most reliable in the dataset, in that they correlated well across both modalities in cells grouped according to their first and second top scoring protein scores: CD45, CD11b, Ly6G. F4/80, Ly6C, CD11c, CD3, CD4, CD117, CD140a, CD26, CD31, CD202b. In contrast, the following antibodies had non-specific expressions across clusters that did not correlate well with their corresponding genes: CD115, CD326, CD335, CD8a, NK1.1. Therefore, we used the former list’s protein and RNA expressions for cell typing, beginning with a visualization of their expressions in cells grouped by Seurat-identified clusters **(Fig S2B)**.

Differentially-expressed protein (DEP) analysis was performed via *FindAllMarkers* (assay = Protein), and the top up- and down-regulated proteins per Seurat cluster were visualized in a heatmap **(Fig S2C, Tables S1-S2)**. We used a logical workflow **(Fig S2D)** to identify immune (CD45+) vs non-immune cells (CD45-), myeloid (CD11b+) vs non-myeloid cells (Cd11b-), followed by neutrophils (Ly6G+), monocytes (CD117+) and macrophages (F4/80+, Ly6C+), T/NK cells (CD3+, CD4+), fibroblasts (CD140a+, CD26+) and endothelial cells (CD31+, CD202b+). Indeed, DEP analysis confirmed the positive and negative expressions of these top proteins for the major cell identities identified **(Fig S2E, Tables S3-S4)**.

For cell clusters that were protein-negative (and to validate the protein-determined cell identities), we performed differentially-expressed gene (DEG) analysis via *FindAllMarkers*

(assay = RNA) **(Table S5)**. The top 20 DEGs for each cluster that were expressed in at least 75% of the cluster’s cells were determined **(Fig S3, Table S6)**. Functional enrichment analysis using EnrichR (as described in ^19^) confirmed our initial cell annotations and revealed the likely identities for the remaining clusters **(Fig S2D)**. Because of the very low cell numbers for clusters 10 and 11, these cells were removed for downstream analysis. A final DEG analysis for the major cell annotations was performed and the top 30 DEGs for each major cell were visualized **(Fig S4, Tables S7-S8)**.

The protein/RNA expressions in the annotated major cells were visualized via *DotPlot* **(Fig S2F)** and via paired UMAP feature plots (via *FeaturePlot*) and scatter plots (via *FeatureScatter*) **(Fig S5)**, further confirming the observed high protein-RNA concordance of the top cell markers in the integrated dataset. Finally, we used violin plots to show the distributions of UMIs and features in cells grouped according to the major cell annotations **(Fig S2G)**.

### Sub-typing and analysis of mouse fibroblasts, macrophages and neutrophils

Fibroblast-, Mo/MΦ- and neutrophil-specific datasets were obtained via the *subset* function (as described in ^19^). The following functions were then performed on each of the subsetted Fibroblast-, Mo/MΦ- and neutrophil-specific datasets: *RunPCA*, *ElbowPlot* **(Fig S7A)**, *FindNeighbors* (dims = 6,9,5, respectively), *FindClusters* (resolution = 0.1), *RunUMAP* (dims = 6,9,5, respectively). DEP and DEG analyses were performed via *FindAllMarkers*, as described earlier for the full dataset **(Table S9 for fibroblast, Table S12 for Mo/M**Φ**, Table S15 for neutrophil)**. Functional enrichment of cell subtype specific DEGs was performed using EnrichR, and the top unique pathways were selected for visualization based on number and proportion of non-overlapping enriched genes.

To test the influence of T2DM on cell subtype transcriptomes, DEG analyses for each cell type sub-cluster were performed using the genotype annotation (db/db vs ND) via *FindAllMarkers* **(Table S10 for fibroblast, Table S13 for Mo/M**Φ**, Table S16 for neutrophil)**. Venny was used to compare and visualize the shared up-regulated genes between sub-clusters **(Table S11 for fibroblast, Table S14 for Mo/M**Φ**, Table S17 for neutrophil)**. Shared genes were then inputted into StringDB for protein-protein interaction (PPI) network analysis, unconnected genes were removed, and the networks were visually annotated according to top unique enriched pathways.

Module scoring of cell subtype signatures was performed via *AddModuleScore* as described previously ^19^, using relevant mouse fibroblast, macrophage and neutrophil gene sets derived from the literature **(Table S18)**.

### Re-analysis and integration of human chronic wound datasets

Individual datasets ^6–8,10,11^ were downloaded as patient-derived samples from the Gene Expression Omnibus (GEO) using *Read10X_GEO*, as described in ^19^. Each dataset was re-analyzed individually for quality control, as described previously and including cell doublet detection via *scDblFinder*, and cell annotation was performed, using the parameters listed in **Table 3**. The output of these re-analyses is shown in **Fig S10A-F**, including UMAP plots of tissue types and cell types via *DimPlot* and a validation of cell type annotation using cell marker genes via *DotPlot*.

**Table 3:** Quality control parameters used for re-analysis of human datasets.

| Dataset GEO accession number: | GSE 223964 | GSE 231643 | GSE 241132 | GSE 248247 | GSE 265972 | GSE 268834 |
| --- | --- | --- | --- | --- | --- | --- |
| Samples | 9 | 8 | 12 | 2 | 9 | 8 |
| nFeature > | 100 | 100 | 50 | 50 | 50 | 50 |
| % MT < | 15 | 15 | 20 | 25 | 20 | 20 |
| % RPS < | 30 | 20 | 20 | 20 | 20 | 20 |
| % RPL < | 40 | 30 | 30 | 30 | 30 | 25 |
| DoubletScore < | 0.5 | 0.5 | 0.5 | 0.5 | 0.5 | 0.5 |
| Final cell count | 25748 | 26439 | 54749 | 403 | 60605 | 71903 |
| Cell types identified | Keratinocyte<br>Fibroblast<br>Macrophage<br>Endothelial<br>Smooth mm<br>T cell<br>B cell<br>Mast cell<br>Melanocyte | Fibroblast<br>Macrophage<br>Neutrophil<br>Endothelial<br>Smooth mm<br>T cell<br>B cell<br>Mast cell<br>Melanocyte | Keratinocyte<br>Fibroblast<br>Macrophage<br>Endothelial<br>Smooth mm<br>T cell<br>B cell<br>Mast cell<br>Melanocyte | Keratinocyte<br>Fibroblast<br>Macrophage<br>Neutrophil<br>Endothelial<br>Smooth mm | Keratinocyte<br>Fibroblast<br>Macrophage<br>Neutrophil<br>Endothelial<br>Smooth mm<br>T cell<br>B cell<br>Mast cell<br>Melanocyte | Keratinocyte<br>Fibroblast<br>Macrophage<br>Neutrophil<br>Endothelial<br>Smooth mm<br>T cell<br>B cell<br>Mast cell<br>Plasma cell |
| Tissue types | Non-diabetic foot ulcers;<br>Diabetic foot ulcers | Diabetic foot ulcers | Non-diabetic acute wounds;<br>Non-diabetic skin | Non-diabetic foot ulcers;<br>Diabetic foot ulcers | Non-diabetic foot ulcers;<br>Non-diabetic skin | Non-diabetic foot ulcers;<br>Diabetic foot ulcers |

Quality-controlled datasets were combined using the *merge* function, followed by *NormalizeData*, *FindVariableFeatures*, *CellCycleScoring*, *ScaleData* (with cell cycle scores as variables to regress) and *RunPCA*. *JackStraw* and *ElbowPlot* were used to choose the first 15 PCs for downstream functions. For initial analyses, all tissue types (acute wounds, chronic wounds and healthy skin) were included to maximize the robustness of data integration and cell type identification. For analyses of diabetic vs non-diabetic chronic wounds, cells from acute wounds were removed.

To test for batch effects, the following were first run using reduction = pca: cell clustering was performed via *FindNeighbors* using ndims = 15, followed by *FindClusters* using Louvain clustering with resolution = 0.3. Dimensionality reduction via UMAP was performed via *RunUMAP* with ndims = 15. The un-integrated merged dataset was visualized via *DimPlot* according to original dataset annotation **(Fig S11A)** and sample ID **(Fig S11B)**, revealing considerable clustering according to the study and sample ID. Therefore, the merged datasets were integrated via *IntegrateLayers* via method = RPCAIntegration, followed by *JoinLayers*. The following were then run using reduction = RPCA: cell clustering was performed via *FindNeighbors* using ndims = 15, followed by *FindClusters* using Louvain clustering with resolution = 0.3. Dimensionality reduction via UMAP was performed via *RunUMAP* with ndims = 15. The integrated dataset was visualized via *DimPlot* according to tissue type annotation **(Fig S11C)** and sample ID **(Fig S11D)**, revealing satisfactory integration of cells. Indeed, cell type distribution analysis according to dataset and sample showed broad representation of all cell types across datasets and samples **(Fig S11E)**.

### Sub-typing of human fibroblasts, macrophages and neutrophils

Fibroblast-, macrophage- and neutrophil-specific datasets were obtained via the *subset* function (as described in ^19^). The following functions were then performed on each of the subsetted fibroblast-, macrophage- and neutrophil-specific datasets: *RunPCA*, *ElbowPlot* **(Fig S12A)**, *FindNeighbors* (dims = 6,10,8, respectively), *FindClusters* (resolution = 0.15,0.2,0.1, respectively), *RunUMAP* (dims = 6,10,8, respectively). Whereas fibroblasts and macrophages were analyzed in all datasets, neutrophils were not found in the acute wound dataset and therefore neutrophil subtypes were analyzed only in the chronic wound datasets. DEG analyses were performed via *FindAllMarkers*, and cell cub-types were annotated according to their top expressed genes. *DimPlot* was used to visualize the cell subtype datasets by grouping according to tissue type, Seurat-determined clusters and final subtype annotation **(Fig S12B-D)**. Fibroblast, macrophage and neutrophil subtype datasets were combined via the *merge* function, and cell subtype distribution analysis showed satisfactory representation across datasets and samples **(Fig S11F)**. The combined dataset was used to analyze DEGs in cell subtypes according to the diabetic vs non-diabetic chronic wound condition via *FindAllMarkers*.

### Cell-cell interaction analyses

CellChat was used to analyze predicted cell-cell interactions of mouse and human fibroblast, macrophage and neutrophil subtype datasets. Standard settings were used for the workflow, as described previously ^19^. For the mouse 3 DPW and cell subtype datasets, the combined Seurat objects were subsetted according to the genotype annotation (db/db vs ND) and analyzed separately before performing differential CellChat analyses. For the human cell subtype dataset, the combined Seurat object was subsetted according to the chronic wound type annotation (Diabetic vs Non-diabetic) and analyzed separately before performing differential CellChat analysis. For individual dataset analyses, signaling scatter plots were visualized via *netAnalysis_signalingRole_scatter* and circle plots were visualized via *netVisual_circle*. For differential analyses, diabetic and non-diabetic CellChat objects were first merged via *mergeCellChat*. Numbers and strength of interactions were visualized via *compareInteractions*. Differential circle plots were visualized via *netVisual_diffInteraction*, and differential heatmap plots were visualized via *netVisual_heatmap*. Differential pathway scatter plots were visualized via *netAnalysis_signalingChanges_scatter*. To narrow down the list of interactions to those that were significantly over-expressed in diabetic vs non-diabetic cells, the following workflow was utilized. First, *identifyOverExpressedGenes* was run with the parameters: only.pos = FALSE, thresh.pc = 0.1, thresh.fc = 0.1, thresh.p = 0.05. Then, DEGs were saved via *netMappingDEG* with thresh = 0.05. Up-regulated genes were identified via *subsetCommunication* with ligand.logFC = 0.20 and receptor.logFC = 0.20, and down-regulated genes were identified with ligand.logFC = -0.20 and receptor.logFC = -0.20. Gene names were extracted from ligand-receptor pairs via *extractGeneSubsetFromPair* and were arranged with interaction and pathway names via *arrange*. Finally, the data were saved as tables for downstream analysis **(Table S19-S20)**. Venny was used to compare and visualize the shared up-regulated ligands and receptors between the mouse and human datasets **(Table S21)**.

### Re-analysis of CD44 knock-out macrophages

The dataset was downloaded from GEO (accession number GSE138445) ^27^. The samples analyzed were of macrophages isolated from alveolar tissue of 6–10-week-old CD44-/- (CD44-KO) and CD44+/+ (WT) female mice (N=3). DEGs were identified, and significantly down-regulated genes (log_2_FC < -0.5, FDR < 0.10) were used for downstream analysis.

### Protein-protein interaction analyses

Singificantly up-regulated genes in db/db vs ND cells, enriched ligand-receptor genes in D vs ND cells, and significantly down-regulated genes in CD44-KO vs WT cells were uploaded to the stringApp on Cytoscape. Protein-protein interaction (PPI) networks were formed using the physical interaction filter and confidence level >0.10-0.80, depending on the analysis.

Unconnected genes were removed, functional enrichment analysis was performed via GO Biological Process, and the top unique pathways were visualized using node border colors.

### Flow cytometry

Single cell suspension from skin wounds was obtained as described above. After washing, all cells were incubated with Zombie NIR™ Fixable Viability Kit (Biolegend, San Diego, CA, USA) for cell viability analysis then followed by Fc blocking by TruStain FcX™ (anti-mouse CD16/32, Biolegend) antibody. Next, skin cells were labeled with antibody cocktails for cell surface markers to define various target populations following manufacturer’s instruction. All samples were analyzed on Cytek ® Aurora cytometer (Cytek Biosciences, Fremont, CA, USA). The complete list of antibodies used is presented in **Table 4**. Data were analyzed using FlowJo (FlowJo LLC, Ashland, OR, USA).

**Table 4:** Antibodies used for flow cytometry.

| Name | Clone | Conjugate | Manufacturer |
| --- | --- | --- | --- |
| Zombie NIR™ |  |  | BioLegend |
| TruStain FcX™ (CD16/32) | 93 |  | BioLegend |
| Ly6G | 1A8 | Alexa Fluor® 700 | BioLegend |
| CD45 | S18009F | FITC | BioLegend |
| CD11b | M1/70 | Alexa Fluor® 594 | BioLegend |
| F4/80 | BM8 | Brilliant Violet 510™ | BioLegend |
| Ly6C | AL-21 | PerCP-Cy™5.5 | BD Biosciences |
| Ly6C | AL-21 | BD Horizon™ BV421 | BD Biosciences |
| CD11c | N418 | Brilliant Violet 605™ | BioLegend |
| CD140a | APA5 | APC | BioLegend |
| CD140a | APA5 | Brilliant Violet 421™ | BioLegend |
| CD44 | IM7 | Brilliant Violet 711™ | BioLegend |
| CD44 | IM7 | Brilliant Violet 421™ | BioLegend |
| CD54 (ICAM-1) | YN1/1.7.4 | PE/Cyanine7 | BioLegend |

### Cell culture experiments

Ten-week-old male non-diabetic (ND; C57BL/6J, Strain # 000664) and diabetic (db/db; BKS.Cg-*Dock7^m^* +/+ *Lepr^db^*/J , Strain # 000642) mice were obtained from The Jackson Laboratory. Bone marrow–derived monocytes (BMDMs) were isolated from femurs and tibias of mice (N=4 mice per group) by flushing the marrow with sterile DPBS containing 5% fetal bovine serum (FBS; Gibco) using a syringe with a 23 gauge needle attached. The bone marrow suspension was passed through a 70 µm cell strainer and centrifuged at 300 g for 5 minutes. Cells were maintained in DMEM (Gibco) supplemented with 10% FBS and 20 ng/mL macrophage colony-stimulating factor (M-CSF; #315-02, Gibco Peprotech) and differentiated for 7 days, with fresh medium supplementation performed on day 4.

For polarization experiments, mature BMDMs (M0) were seeded into 24-well culture plates at a density of 200,000 cells per well (n=3 per group). Cells were stimulated with 100 ng/mL lipopolysaccharide (LPS; #L4391, Sigma) and 25 ng/mL interferon-γ (IFN-γ; #315-05, Gibco Peprotech) for M1 activation, or with 20 ng/mL interleukin-4 (IL-4; #214-14, Gibco Peprotech) for M2a activation, and 25 ng/mL interleukin-10 (IL-10; #210-10, Gibco Peprotech) and 50nM Dexamethasone (DEX, #D4902, Sigma) for M2c activation prior to downstream analyses. For CD44 blocking experiments, BMDMs were treated with rat anti-mouse CD44 antibody in serum-free DMEM medium (5 μg/mL; clone IM7, #BDB741057, BD Biosciences) for 30 minutes prior to the addition of polarization stimuli.

After 48 hours, total RNA was isolated from cells using the Quick-RNA Miniprep Kit (#R1055, Zymo Research). RNA concentration was determined using a NanoDrop Ultra spectrophotometer (Thermo Fisher Scientific), and equal amounts of RNA were used for cDNA synthesis with the RevertAid RT Reverse Transcription Kit (#K1691, Thermo Fisher Scientific). Quantitative real time PCR (RT-qPCR) was performed using Fast SYBR™ Green Master Mix (#4385612, Applied Biosystems), and reactions were run on a QuantStudio™ 3 Real-Time PCR System (Applied Biosystems). Relative mRNA expression levels were calculated using the ΔΔCt method, with M0 cells from one ND mouse used as the reference control. Four biological replicates were analyzed. Primer sequences are provided in **Table 5**.

**Table 5:**
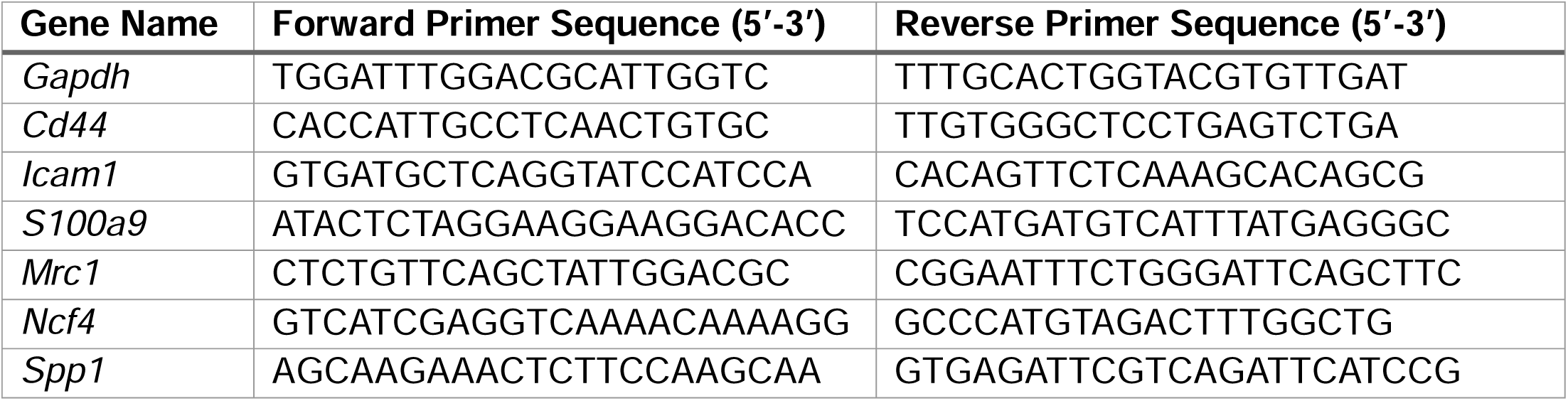
Primer sequences used for RT-qPCR.

### Local treatment with CD44 blocking antibody

To assess the function of CD44 in wounds, 20μg purified rat anti-mouse CD44 antibody (clone IM7, BD Biosciences, San Jose, CA, USA) was injected intradermally around the periphery of each wound in adult db mice (N=4 mice per group) at four evenly distributed sites starting immediately after wounding day 0 and day 2 post-injury. Control mice received injections of an equal amount of purified rat IgG2b, k isotype control (A95-1, BD Biosciences). All wounds were harvested 24 hours after the last injection (day 3 post-injury), and cells were dissociated from wounds and analyzed by flow cytometry as described above.

To assess the effects of CD44 blocking on wound closure, digital images were taken of the wound surface. Wound size was measured on the days when injections were performed and the day of wound collection using Fiji Image J and expressed as a percentage of the original wound area ((Day X area/Day 0 area) ×100).

### Statistical analysis

Statistical analyses were performed using GraphPad Prism (version 11) or R (version 4.4.1) in RStudio (version 2025.05.0). For comparison of more than two groups, two-way ANOVA and Tukey’s multiple comparisons post-hoc tests were used. For Pearson correlation analyses, two-sided P values were computed for each pair of variables tested. A P value of less than 0.05 was considered significant.

### Code availability

All code to recreate the analyses and figures in this manuscript will be posted on the Wietecha Lab Github upon publication.

### Data availability

The authors declare that all data supporting the findings of this study are available within the article and its supplementary materials, or from the corresponding authors upon request. The protein and RNA single-cell sequencing datasets of mouse db/db and ND wounds will be uploaded to Gene Expression Omnibus, and access will be released upon manuscript publication. All other transcriptomics datasets analyzed in this manuscript are listed in **Fig 7A** and **Table 3**.

## RESULTS

### Temporal dynamics of major dermal cells in diabetic mouse wounds

To study cell dynamics of major dermal cell populations during T2DM impaired wound healing, we harvested full-thickness, excisional wounds from diabetic (db/db) and non-diabetic (ND) mice at three time points post-injury (capturing the inflammatory, proliferative and resolution phases of healing), digested the dermal and granulation tissue layers, antibody stained for cell-specific surface markers, and performed single-cell multi-omics analysis via the 10X Genomics pipeline **(Fig 1A)**. Following robust quality control and clustering protocols ^19^ that resulted in the final dataset consisting of 49,963 high-quality cells **(see Methods, Fig S1A-D)**, we performed protein de-complexing to annotate each cell with its top scoring antibody ^26^ **(Fig S1E, Fig S2A-B)**. This annotation projected on a UMAP plot showed that CD11b/Ly6G positive cells (neutrophil markers), F4-80/Ly6C/CD11c positive cells (monocyte/macrophage markers), CD3/CD4 positive cells (T lymphocyte markers), and CD140a/CD26 positive cells (fibroblast markers) all formed individual clusters separate from one another and from CD31 positive cells (endothelial marker) **(Fig 1B)**. Visualizing the cell annotations according to the days post-wounding (DPW) at which the cells were harvested **(Fig 1C)** and the mouse genotype **(Fig 1D)** showed strong DPW- and genotype-associated cell distributions across most cell clusters, especially those positive for neutrophil, monocyte/macrophage and fibroblast markers. We then determined the differentially expressed proteins (DEPs) **(Fig S2C-S2E, Tables S1-S4)** and differentially expressed genes (DEGs) **(Fig S3-S4, Tables S5-S8)** for each of the cell clusters, and we annotated them as major cell types **(Fig S2D)**, in order from most to least abundant: fibroblasts, monocytes/macrophages (Mo/MΦ), neutrophils, endothelial, smooth muscle, T/NK, muscle progenitor and skeletal muscle cells **(Fig 1E)**. Comparisons of the protein and RNA expressions of the major cell markers within the annotated cell types revealed very high concordance between the two orthogonal assays, confirming our confidence in the cell type annotations **(Fig 1F, Fig S2F, Fig S5)**.

**Figure 1.**
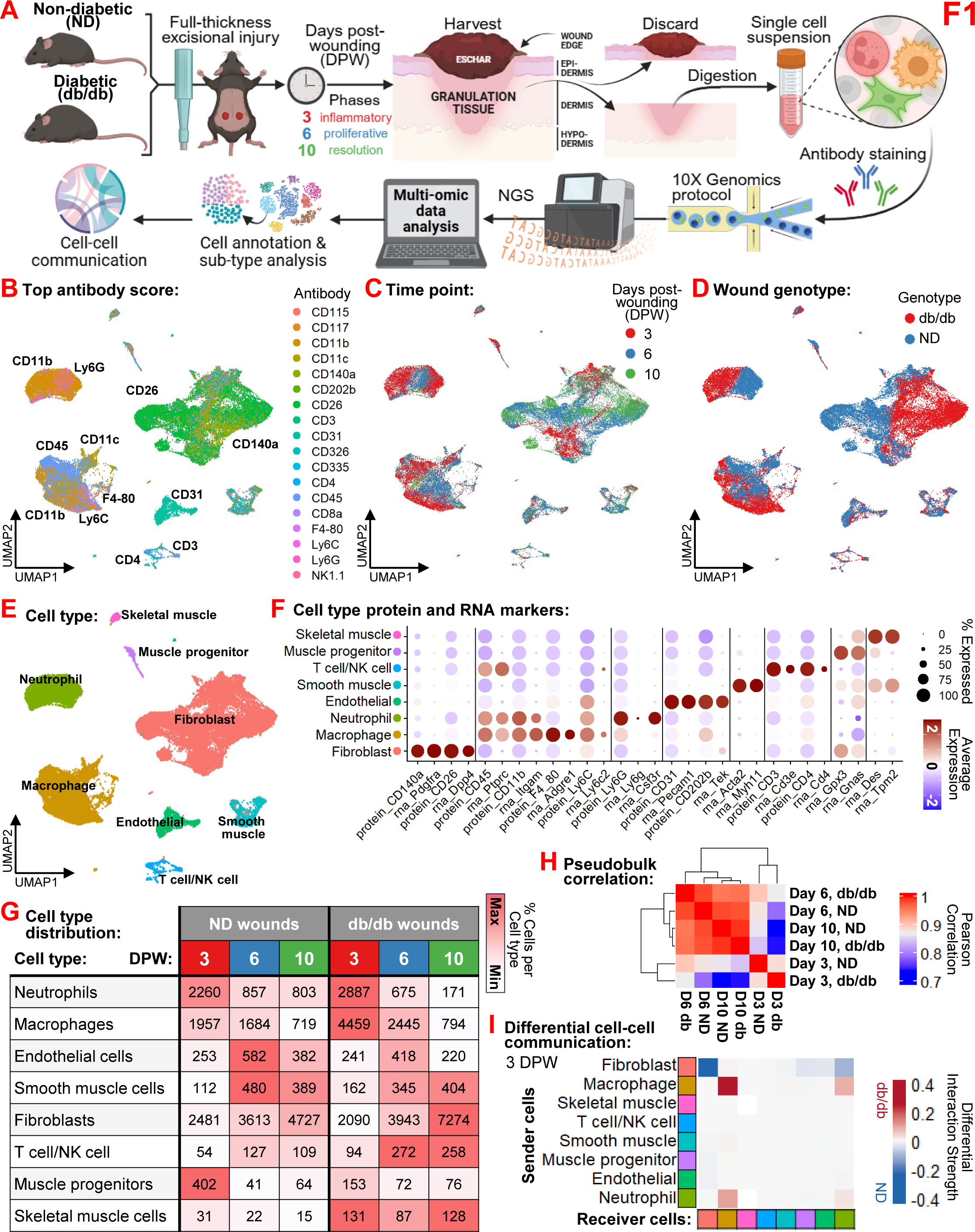
Multi-omic single-cell landscape of diabetic dermal wound healing dynamics. (A) Experimental workflow for the single-cell multi-omic analysis of dermal wound healing. Full-thickness excisional injuries were performed on non-diabetic (ND) and diabetic (db/db) mice. Wounds were harvested at 3 (inflammatory), 6 (proliferative), and 10 (resolution) days post-wounding (DPW) and processed for single-cell multi-omics (simultaneous surface protein and RNA profiling). (B–D) UMAP visualization of 49,963 integrated cells from all conditions, colored by (B) top-scoring antibody-derived surface proteins, (C) harvest time point (DPW 3, 6, and 10), and (D) wound genotype (ND vs db/db). (E) UMAP of cells according to major cell type annotations, including: fibroblasts, macrophages, neutrophils, endothelial cells, smooth muscle cells, T/NK cells, muscle progenitor cells, skeletal muscle cells. (F) Dot plot showing the concordance between average surface protein expression and corresponding RNA marker expression across all annotated cell types. (G) Distribution table of absolute cell counts per major cell type across genotypes and time points. (H) Pearson correlation heatmap of pseudobulk transcriptomes across time points and genotypes. (I) Differential cell-cell communication interaction strength at 3 DPW between db/db and ND wounds, highlighting altered sender-receiver dynamics in the early-phase diabetic wound microenvironment.

We then performed cell type distribution analysis according to both genotype (db/db vs ND) and DPW (3, 6, 10) and visualized them as a heatmap **(Fig 1G)**. This revealed high cell yields (>2000 cells per category) for three major cell types across genotypes and DPW, namely for neutrophils, Mo/MΦ and fibroblasts. These cell types showed notable differences for db/db vs ND wounds, particularly at 3 DPW, when db/db wounds had 27.7% more neutrophils, 127.4% more Mo/MΦ, and 16.8% less fibroblasts than ND wounds in absolute cell numbers.

Pseudobulk quantification of the transcriptomes of all cells across the genotypes and DPW followed by Pearson correlation analysis confirmed that major differences between db/db and ND wounds occurred at 3 DPW **(Fig 1H)**. To analyze cell-cell communication (CCC) differences between db/db and ND wounds at 3 DPW, we performed differential CCC analyses **(Fig S6)**.

We visualized the differential interaction strengths between major cell types in db/db vs ND wounds via a heatmap, which revealed that, in db/db wounds, there were major increases in outgoing interactions from neutrophils and Mo/MΦ to other neutrophils and Mo/MΦ, paired with major decreases in outgoing interactions from fibroblasts to other cell types **(Fig 1I)**.

These major disturbances in cell type abundances and cell-cell communications across fibroblasts, Mo/MΦ, and neutrophils during the early stages of wound healing in diabetic wounds encouraged us to study each of these cell types in more detail.

### Three fibroblast subtypes are dysregulated in diabetic mouse wounds

We first sub-clustered fibroblasts from the overall dataset and visualized through UMAP plots their DPW **(Fig 2A)** and genotype **(Fig 2B)** distributions. Sub-clustering resulted in four fibroblast subtypes, termed fα, fβ, fγ and fδ **(Fig 2C)**. We performed fibroblast subtype distribution analysis according to both genotype and DPW, revealing major differences in their temporal dynamics between db/db and ND wounds **(Fig 2D)**. Notably, db/db wounds had 68.4% less cluster fβ fibroblasts than ND wounds at 3 DPW and nearly double the numbers of cluster fγ fibroblasts at 6 and 10 DPW in absolute cell numbers.

**Figure 2.**
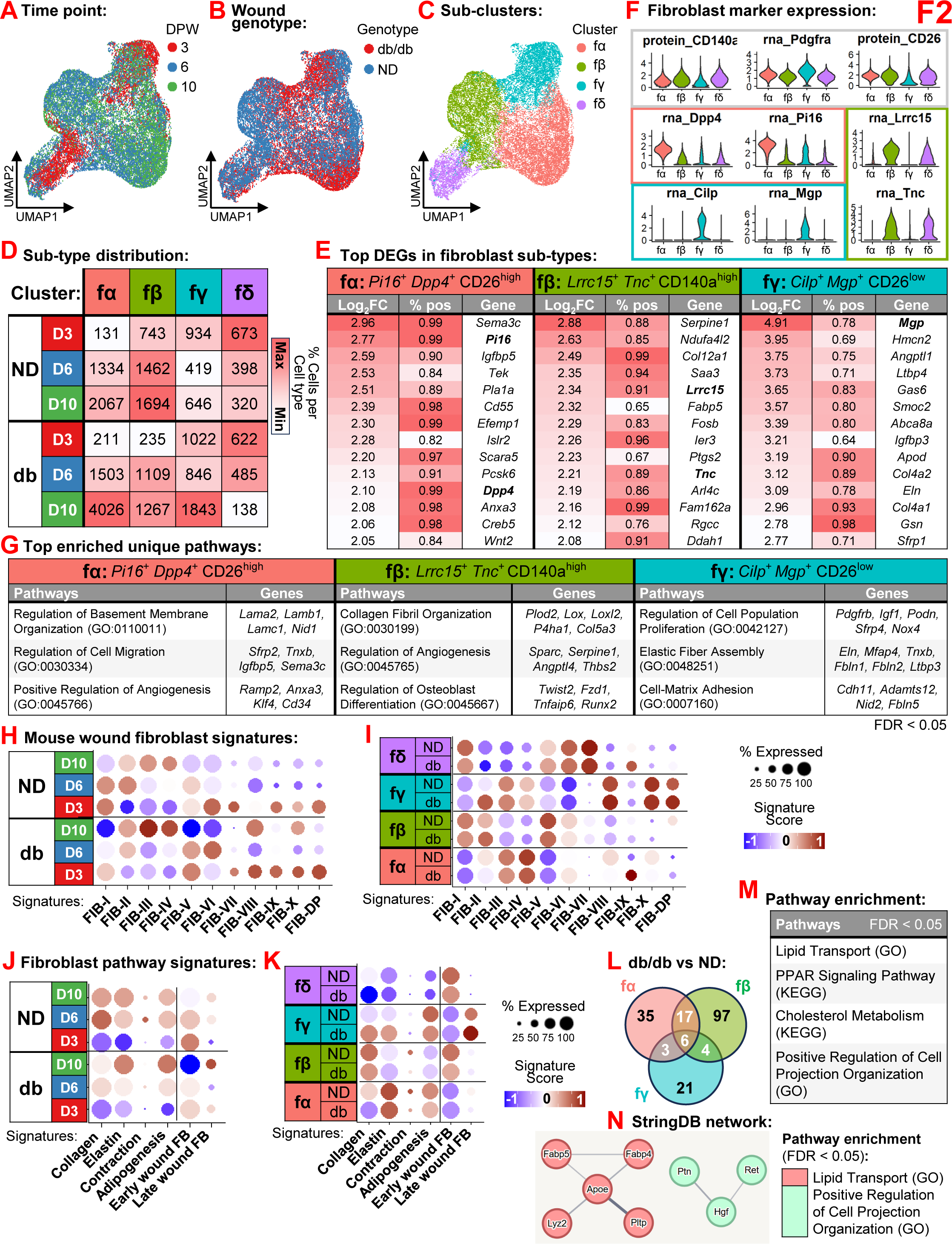
Temporal dysregulation of fibroblast subtypes in diabetic wounds. (A–C) UMAP projection of sub-clustered fibroblasts colored by (A) harvest time point (3, 6, 10 days post-wounding (DPW)), (B) wound genotype (non-diabetic (ND) vs db/db), and (C) four identified subtypes: fα, fβ, fγ and fδ. (D) Cell count distribution of fibroblast subtypes across DPW in db/db and ND wounds. (E) Top differentially expressed genes (DEGs) expressed in at least 50% of the cells for the identified fibroblast subtypes. Bolded genes are also shown in violin plots in (F). (F) Violin plots showing subtype-specific protein (CD140a, CD26) and RNA (*Pdgfra, Dpp4, Cilp, Pi16, Lrrc15, Mgp, Tnc*) expression. (G) Functional enrichment of unique gene ontology (GO) pathways in fα, fβ and fγ subtypes, including examples of the top enriched genes. (H–I) Dot plots comparing fibroblasts to published mouse wound fibroblast signatures ^28^ between time points and genotypes (H) and between fibroblast subtypes (I). (J–K) Dot plots comparing fibroblasts to curated Reactome pathway signatures and early vs late wound fibroblast signatures ^3^ between time points and genotypes (J) and between fibroblast subtypes (K). (L) Comparison of DEGs in fibroblast subtypes in db/db and ND wounds. (M) Pathway enrichment analysis of shared fibroblast subtype genes up-regulated in db/db wounds showing top unique pathways from curated GO and Kyoto Encyclopedia of Genes and Genomes (KEGG) databases. (N) StringDB protein-protein interaction network of shared up-regulated genes associated with enriched GO pathways in db/db vs ND fibroblasts.

We performed DEP and DEG analyses to find fibroblast subtype specific markers **(Table S9)**, and we visualized via a heatmap the top DEGs that are also expressed by greater than 50% of the cells **(Fig 2E)**. This analysis showed that cluster fδ was identified by cell cycle genes, therefore it was annotated as a Proliferative Fibroblast **(Fig S7B)**. We used violin plots to show the expressions of major fibroblast markers identified for the four subtypes **(Fig 2F)**, including the protein expressions of canonical fibroblast markers CD140a and CD26 (top), the RNA expressions of cluster fα markers *Dpp4* and *Pi16* (middle), the RNA expressions of cluster fβ markers *Lrrc15* and *Tnc* (right), and the RNA expressions of cluster fγ markers *Cilp* and *Mgp* (bottom). They also showed that cluster fδ resembled fβ in its marker expression. Due to these marker protein and gene analyses, we labeled the fibroblast subtypes as Pi16^+^Dpp4^+^CD26^high^, Lrrc15^+^Tnc^+^CD140a^high^ and Cilp^+^Mgp^+^CD26^low^ and for clusters fα, fβ and fγ, respectively.

Pathway enrichment analyses of the top fibroblast subtype marker genes showed that the fα: Pi16^+^Dpp4^+^CD26^high^ fibroblasts were enriched for basement membrane and cell migration pathways, fβ: Lrrc15^+^Tnc^+^CD140a^high^ fibroblasts were enriched for collagen formation and angiogenesis pathways, and the fγ: Cilp^+^Mgp^+^CD26^low^ fibroblasts were highly enriched for elastin formation and regulation of cell proliferation pathways **(Fig 2G)**.

To compare the fibroblast transcriptomes from db/db and ND wounds to published wound fibroblast subtype signatures, we performed module scoring analyses with gene sets derived from a recent murine wound atlas that characterized ten fibroblast subtypes, termed FIB-I through -X and DP for dermal papilla ^28^. The module scores of these signatures were first analyzed across all fibroblasts separated by their DPW and genotype of origin, and this showed expected patterns of time-resolved enrichment of the major fibroblast subtypes previously identified in large skin wounds, with FIB-I,V expressed at 3 DPW and FIB-II expressed at 6 and 10 DPW in both genotypes **(Fig 2H)**. The FIB-IV,IX signatures were high in the fα: Pi16^+^Dpp4^+^CD26^high^ subtype, the FIB-I,II,V fibroblast signatures were high in the fβ: Lrrc15^+^Tnc^+^CD140a^high^ fibroblast subtype, the FIB-VIII,X,DP signatures were high in the fγ: Cilp^+^Mgp^+^CD26^low^ subtype, the FIB-III signature was shared among fα and fγ, and the proliferative fδ subtype was enriched in FIB-I,VI,VII signatures **(Fig 2I)**. Interestingly, the diabetic condition highly skewed expression for the FIB-IX signature in the fα and fδ subtypes.

We also compared the fibroblast transcriptomes to gene sets derived from relevant wound fibroblast phenotype Reactome pathways (*e.g.* collagen formation, elastin formation, contraction and adipogenesis) and our previous work that characterized early versus late wound fibroblasts using bulk sequencing ^3^. The module scores of these signatures were first analyzed across all fibroblasts separated by their DPW and genotype of origin, and this showed an enrichment for the early wound fibroblast signature in the ND fibroblasts at 3 and 6 DPW and for the late wound fibroblast signature at 10 DPW, while the collagen and contraction signatures were enriched in the ND fibroblasts at 6 DPW **(Fig 2J)**. In diabetic fibroblasts, adipogenesis and late wound fibroblast signatures were more highly enriched at 10 DPW while the other signatures were less expressed throughout healing. The early wound fibroblast signature was especially high in fβ: Lrrc15^+^Tnc^+^CD140a^high^ and proliferative fδ fibroblast subtypes, the collagen signature was high in the fβ: Lrrc15^+^Tnc^+^CD140a^high^ subtype, the elastin signature was high in the fα: Pi16^+^Dpp4^+^CD26^high^ subtype, and the late wound fibroblast signature was high in the fγ: Cilp^+^Mgp^+^CD26^low^ subtype **(Fig 2K)**. Interestingly, the diabetic condition skewed the fγ: Cilp^+^Mgp^+^CD26^low^ subtype toward collagen and elastin pathways, while the ND condition skewed fα: Pi16^+^Dpp4^+^CD26^high^ and fβ: Lrrc15^+^Tnc^+^CD140a^high^ subtypes toward contraction pathways.

To analyze how T2DM may polarize fibroblast subtypes, we compared the DEGs from db/db vs ND analyses for all subtypes **(Table S10)** using a Venn diagram, and this showed that 30 genes were commonly up-regulated by T2DM in at least two fibroblast subtypes **(Fig 2L, Table S11)**. These genes enriched for a few unique pathways, including lipid transport, PPAR signaling and cell projection regulation **(Fig 2M)**. We then used StringDB to create protein-protein interaction networks of these T2DM-regulated genes, revealing two highly connected networks involved in lipid transport (including *Pltp, Apoe,* and *Fabp4/5*) and cell projection organization (including *Ptn* and *Ret*) **(Fig 2N)**.

In sum, these results suggest that T2DM dysregulates wound fibroblast subtype dynamics in healing wounds, decreasing signatures characterized by early wound fibroblasts as well as contraction and collagen formation phenotypes in early-to-mid phase wounds while increasing adipogenesis and lipid transport features in late phase wounds.

### Three monocyte/macrophage subtypes are dysregulated in diabetic mouse wounds

We next subset monocytes/macrophages (Mo/MΦ) from the overall dataset and through UMAP plots visualized their DPW **(Fig 3A)** and genotype **(Fig 3B)** distributions. Sub-clustering resulted in four Mo/MΦ subtypes, termed mα, mβ, mγ and mδ **(Fig 3C)**. We previously identified via FACS and single-cell sequencing four Mo/MΦ subtypes in the wounds of db/db mice, which were distinguished by their differential expression for surface markers CD11b, F4/80, Ly6C and CD11c ^17,29^. Therefore, we generated flow cytometry-like scatter plots for these markers using the protein assay of the single-cell multi-omics dataset **(Fig 3D)**. These plots showed that clusters mα and mβ had high CD11b, cluster mα had high F4/80, cluster mβ had high Ly6C, and cluster mγ had high CD11c expressions. Violin plots of these markers at the protein and RNA levels confirmed these observations, including that cluster mδ resembled mα in its surface marker expression **(Fig 3E)**. We then performed Mo/MΦ subtype distribution analysis according to both genotype and DPW, and by visualizing them as a heatmap we showed major temporal differences in Mo/MΦ subtype dynamics between db/db and ND wounds **(Fig 3F)**. Indeed, while clusters mβ and mγ had similar cell abundances across the three time points in both genotypes, db/db wounds had 8.6 times more cluster mα cells at 3 DPW, 2.1 times more at 6 DPW and 1.8 times more at 10 DPW in absolute cell numbers.

**Figure 3.**
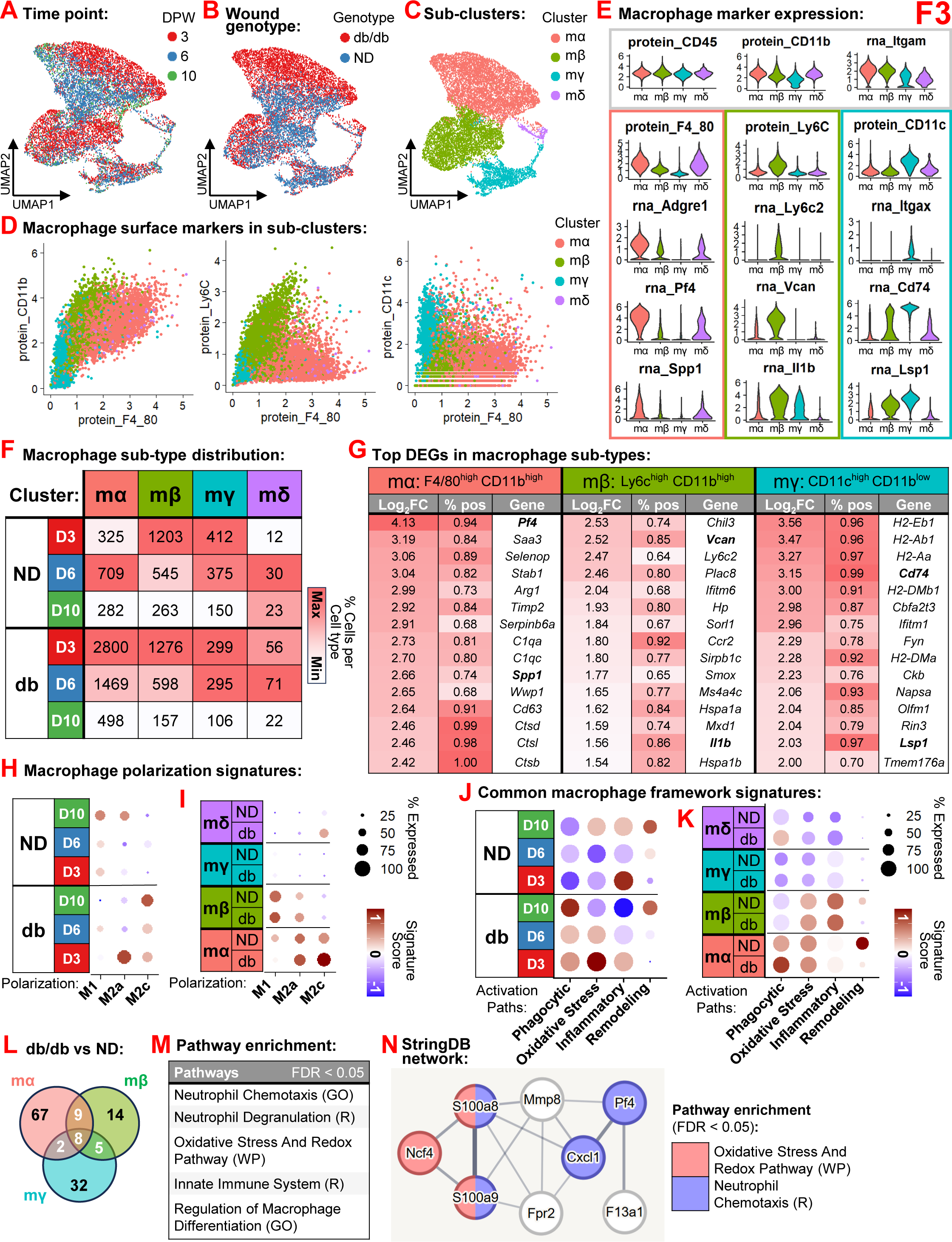
Altered macrophage polarization and subtype signatures in diabetic wounds. (A–C) UMAP of integrated monocyte/macrophage (Mo/MΦ) sub-clusters colored by (A) harvest time point (3, 6, 10 days post-wounding (DPW)), (B) wound genotype (non-diabetic (ND) vs db/db), and (C) four identified subtypes: mα, mβ, mγ and mδ. (D) Surface marker FACS-like scatter plots showing differential expressions of CD11b, F4/80, Ly6C and CD11c across Mo/MΦ subtypes. (E) Violin plots showing expressions of protein (CD45, CD11b, F4/80, Ly6C, CD11c) and RNA (*Itgam, Adgre1, Ly6c2, Itgax, Pf4, Vcan, Cd74, Spp1, Il1b*) markers in Mo/MΦ subtypes. (F) Cell count distribution of Mo/MΦ subtypes across DPW in db/db and ND wounds. (G) Top differentially expressed genes (DEGs) expressed in at least 50% of the cells for the identified Mo/MΦ subtypes. Bolded genes are also shown in violin plots in (E). (H–I) Dot plots comparing Mo/MΦ to literature-derived macrophage polarization signatures (M1, M2a, M2c) between time points and genotypes (H) and between Mo/MΦ subtypes (I). (J–K) Dot plots comparing Mo/MΦ to published common activation framework path (Phagocytic, Oxidative Stress, Inflammatory, Remodeling) signatures ^31^ between time points and genotypes (J) and between Mo/MΦ subtypes (K). (L) Comparison of DEGs in Mo/MΦ subtypes in db/db and ND wounds. (M) Pathway enrichment analysis of shared Mo/MΦ subtype genes up-regulated in db/db wounds showing top unique pathways from curated gene ontology (GO), Reactome (R) and WikiPathways (WP) databases. (N) StringDB protein-protein interaction network of shared up-regulated genes associated with enriched GO pathways in db/db vs ND Mo/MΦ.

We performed DEP and DEG analyses to find Mo/MΦ subtype specific markers **(Table S12)**, and we visualized via a heatmap the top DEGs that are also expressed by greater than 50% of the cells **(Fig 3G)**. This analysis showed that cluster mδ was identified by cell cycle genes, therefore it was annotated as a Proliferative Macrophage **(Fig S7C)**. Violin plots confirmed the differential expressions of major Mo/MΦ markers identified for the subtypes **(Fig 3E)**, including of cluster mα markers *Adgre1*, *Pf4* and *Spp1* (left), cluster mβ markers *Ly6c2*, *Vcan* and *Il1b* (middle), and cluster mγ markers *Itgax*, *Cd74* and *Lsp1* (right). Due to these marker protein and gene analyses, we labeled the Mo/MΦ subtypes as F4/80^high^CD11b^high^, Ly6c^high^CD11b^high^ and CD11c^high^CD11b^low^ for clusters mα, mβ and mγ, respectively.

To compare the Mo/MΦ transcriptomes from db/db and ND wounds to classical macrophage polarization signatures, we performed module scoring analyses with gene sets derived from the literature describing the gene expression signatures of macrophages polarized *in vitro* into the M1 (pro-inflammatory), M2a (anti-inflammatory) or M2c (phagocytic) phenotypes ^30^. The module scores of these signatures were first analyzed across all Mo/MΦ separated by their DPW and genotype of origin, and this showed an overall enrichment of M1 signature in ND Mo/MΦ across all time-points and of M2a at 10 DPW, whereas diabetic Mo/MΦ were less enriched in the M1 signature throughout healing and highly enriched in M2a at 3 DPW and remarkably high in M2c throughout healing but especially at 10 DPW **(Fig 3H)**. In terms of Mo/MΦ subtypes, the mα: F4/80^high^CD11b^high^ subtype was enriched for both M2a and M2c signatures, and the mβ: Ly6c^high^CD11b^high^ subtype was enriched for M1 **(Fig 3I).** Interestingly, the diabetic condition skewed the mα: F4/80^high^CD11b^high^ and proliferative mδ Mo/MΦ subtypes toward the M2c signature.

To compare the Mo/MΦ transcriptomes from db/db and ND wounds to published Mo/MΦ activation signatures, we performed module scoring analyses with gene sets derived from a major study that devised a comprehensive framework for signatures of monocyte-derived macrophages during their *in vivo* maturation and activation toward phagocytic, oxidative stress, inflammatory or remodeling phenotypes ^31^. The module scores of these signatures were first analyzed across all Mo/MΦ separated by their DPW and genotype of origin, and this showed a remarkable enrichment for the phagocytic and oxidative stress signatures in the diabetic Mo/MΦ across healing, especially at 3 and 6 DPW, whereas the inflammatory and remodeling signatures were enriched in the ND Mo/MΦ at 3 and 6 DPW, respectively **(Fig 3J)**. The phagocytic signature was especially high in the F4/80^high^CD11b^high^ Mo/MΦ subtype, and this was further increased in the diabetic condition, whereas the remodeling signature was strongly increased in the ND condition of this Mo/MΦ subtype **(Fig 3K)**.

To analyze how T2DM may polarize Mo/MΦ subtypes, we compared the DEGs from db/db vs ND analyses for all subtypes **(Table S13)** using a Venn diagram, and this showed that 24 genes were commonly up-regulated by T2DM in at least two Mo/MΦ subtypes **(Fig 3L, Table S14)**.

These genes enriched for several unique pathways, including neutrophil degranulation, neutrophil chemotaxis and oxidative stress **(Fig 3M)**. We then used StringDB to create a protein-protein interaction network of these T2DM-regulated genes, revealing that several are highly connected and involved in oxidative stress and neutrophil chemotaxis pathways, including *S100a8* and *S100a9* **(Fig 3N)**.

In sum, these results suggest that T2DM polarizes wound Mo/MΦ toward altered inflammatory signatures characterized by phagocytosis and oxidative stress, and that a specific F4/80^high^CD11b^high^ Mo/MΦ subtype is the primary driver of this early immune dysfunction in diabetic wounds.

### Two neutrophil subtypes are dysregulated in diabetic mouse wounds

We subset neutrophils from the overall dataset and visualized through UMAP plots their DPW **(Fig 4A)** and genotype **(Fig 4B)** distributions. Sub-clustering resulted in two neutrophil subtypes, termed nα and nβ **(Fig 4C)**, which split along the db/db-ND distribution **(Fig 4B)**. We then performed neutrophil subtype distribution analysis according to both genotype and DPW, and by visualizing them as a heatmap we confirmed this remarkable genotype split between the two subtypes **(Fig 4D)**. Indeed, 82.6% of cluster nβ neutrophils were represented in ND wounds and 54.7% of them were present at 3 DPW, whereas 93.7% of cluster nα neutrophils were represented in db/db wounds, with 85.3% of them were present at 3 DPW in absolute cell numbers.

**Figure 4.**
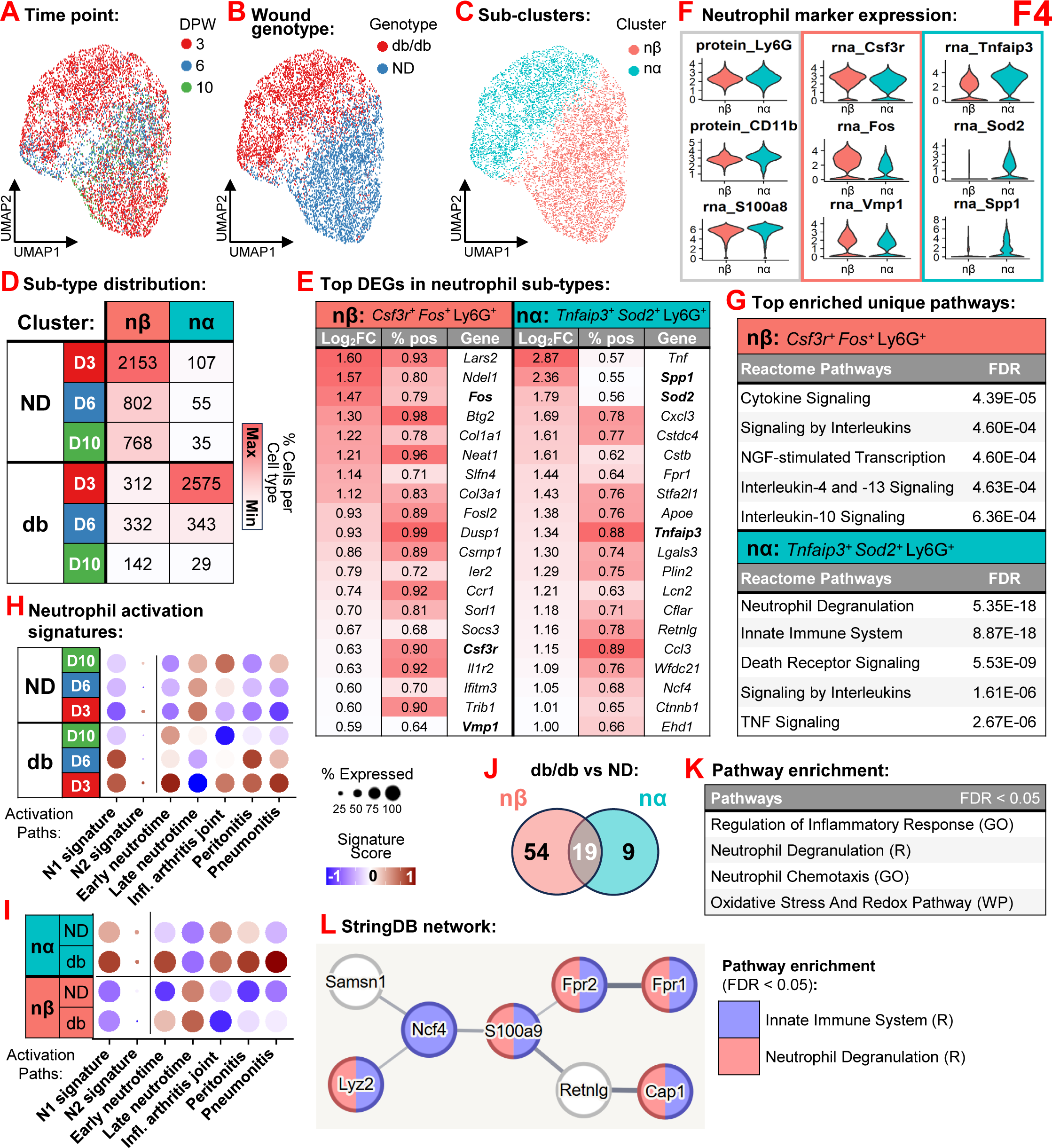
Neutrophil heterogeneity and subtype signatures in diabetic wounds. (A–C) UMAP visualization of neutrophil sub-clusters colored by (A) harvest time point (3, 6, 10 days post-wounding (DPW)), (B) wound genotype (non-diabetic (ND) vs db/db), and (C) two identified subtypes: nα and nβ. (D) Cell count distribution of neutrophil subtypes across DPW in db/db and ND wounds. (E) Top differentially expressed genes (DEGs) expressed in at least 50% of the cells for the identified neutrophil subtypes. Bolded genes are also shown in violin plots in (F). (F) Violin plots showing expressions of protein (Ly6G, CD11b) and RNA (*Csf3r, Tnfaip3, Fos, Sod2, S100a8, Vmp1, Spp1*) markers in neutrophil subtypes. (G) Functional enrichment of unique Reactome pathways in neutrophil subtypes. (H–I) Dot plots comparing neutrophils to published N1/N2 signatures ^32^ and "Neutrotime" signatures (Early/Late Neutrotime and local/systemic chronic inflammation) ^33^ between time points and genotypes (H) and between neutrophil subtypes (I). (J) Comparison of DEGs in neutrophil subtypes in db/db and ND wounds. (K) Pathway enrichment analysis of shared neutrphil subtype genes up-regulated in db/db wounds showing top unique pathways from curated gene ontology (GO), Reactome (R) and WikiPathways (WP) databases. (L) StringDB protein-protein interaction network of shared up-regulated genes associated with enriched GO pathways in db/db vs ND neutrophils.

We performed DEG analyses to find neutrophil subtype specific markers **(Table S15)**, and we visualized via a heatmap the top DEGs that are also expressed by greater than 50% of the cells **(Fig 4E)**. We used violin plots to show the expressions of major neutrophil markers identified for the two subtypes **(Fig 4F)**, including the protein expressions of canonical neutrophil markers Ly6G and CD11b (left), the RNA expressions of cluster nβ markers *Csf3r* and *Fos* (middle), and the RNA expressions of cluster nα markers *Tnfaip3*, *Sod2* and *Spp1* (right). Due to these marker gene analyses, we labeled the neutrophil subtypes as Tnfaip3^+^Sod2^+^Ly6G^+^ and Csf3r^+^Fos^+^Ly6G^+^ for clusters nα and nβ, respectively.

Pathway enrichment analyses of the top neutrophil subtype marker genes showed that the nβ: Csf3r^+^Fos^+^Ly6G^+^ neutrophils were enriched for cytokine and Interleukin signaling pathways, whereas the nα: Tnfaip3^+^Sod2^+^Ly6G^+^ neutrophils were highly enriched for neutrophil degranulation, death receptor and TNF signaling pathways **(Fig 4G)**.

To compare the neutrophil transcriptomes from db/db and ND wounds to published neutrophil activation signatures, we performed module scoring analyses with gene sets derived from two studies: one that alternatively polarized neutrophils in vitro toward the pro-inflammatory, N1 phenotype (via LPS and IFN-γ) or the anti-inflammatory, N2 phenotype (via IL-4) ^32^; and another study that comprehensively captured signatures of neutrophils during their *in vivo* maturation (termed Neutrotime) and activation within chronic inflammatory conditions such as arthritis joints, peritonitis and pneumonitis ^33^. The module scores of these signatures were first analyzed across all neutrophils separated by their DPW and genotype of origin, and this showed a remarkable enrichment for N1, early Neutrotime and all three *in vivo* chronic inflammatory conditions in the diabetic neutrophils, especially at 3 and 6 DPW, whereas the late Neutrotime signature was enriched in the ND neutrophils **(Fig 4H)**. The N1 and chronic inflammatory signatures were high in the nα: Tnfaip3^+^Sod2^+^Ly6G^+^ neutrophil subtype, and this was further increased in the db/db vs ND condition **(Fig 4I)**.

To further analyze how T2DM may polarize neutrophil subtypes, we compared the DEGs from db/db vs ND analyses on the two subtypes **(Table S16)** using a Venn diagram, and this showed that 19 genes were commonly up-regulated by T2DM in the two neutrophil subtypes **(Fig 4J, Table S17)**. These genes enriched for several notable pathways, including neutrophil degranulation, neutrophil chemotaxis, and oxidative stress **(Fig 4K)**. We then used StringDB to create a protein-protein interaction network of these T2DM-regulated genes, revealing that they are highly connected and centered around a hub of genes involved in neutrophil degranulation, including *Fpr1/2, S100a9* and *Lyz2* **(Fig 4L)**.

In sum, these results suggest that T2DM polarizes infiltrating wound neutrophils toward a chronic inflammatory signature characterized by neutrophil degranulation and oxidative stress, and that a specific Tnfaip3^+^Sod2^+^Ly6G^+^ neutrophil subtype is the primary driver of this early immune dysfunction in diabetic wounds.

### Cell-cell communication disruptions of T2DM mouse wound cell subtypes

We integrated our multi-omic single-cell datasets of two neutrophil, four macrophage and four fibroblast subtypes, and a combined dot plot confirmed the shared and differential expressions of all subtype protein and RNA markers **(Fig 5A)**. We then split the data into db/db and ND datasets at 3 DPW and performed individual comprehensive CCC analyses via CellChat.

**Figure 5.**
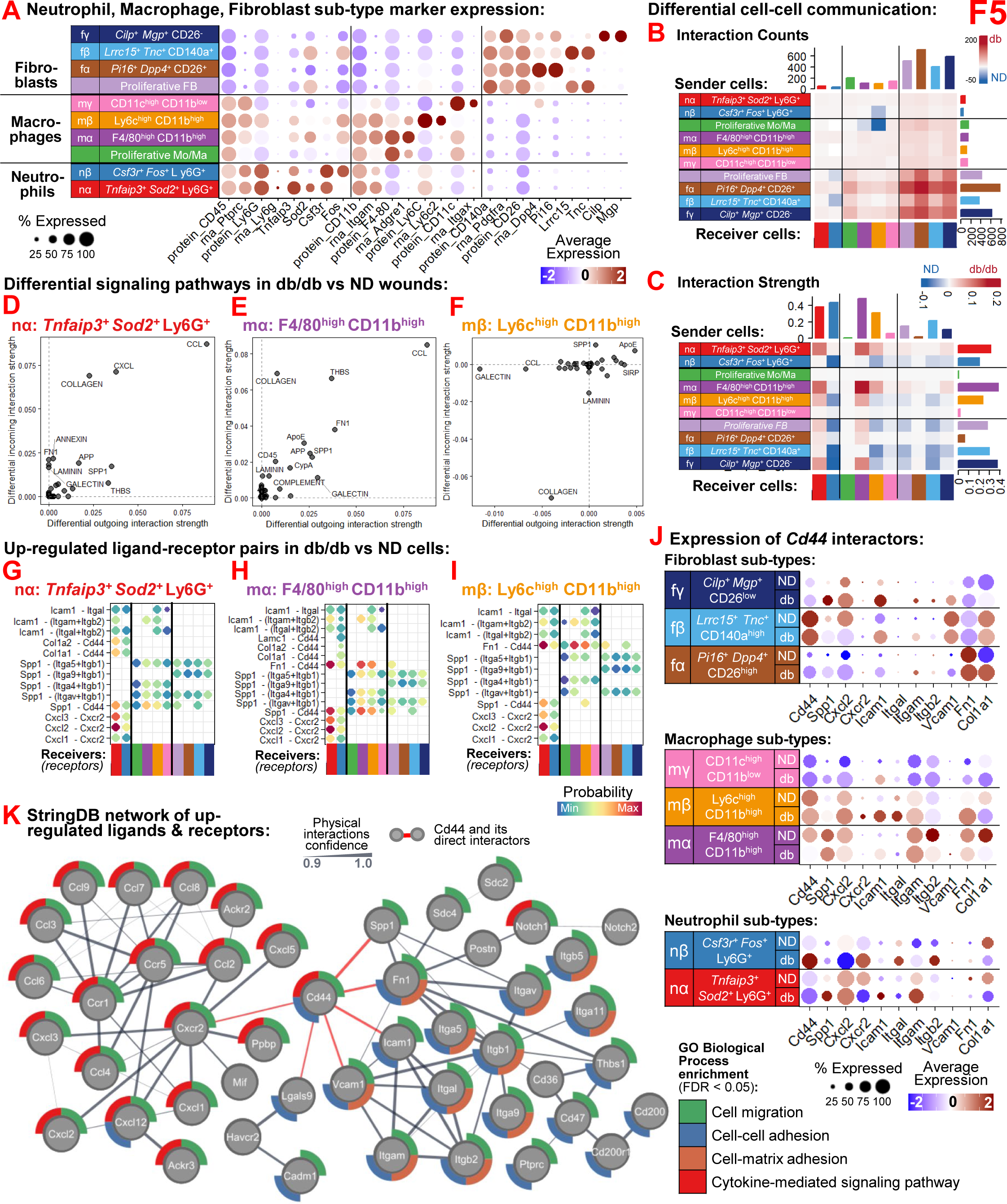
The CD44-mediated immune-stromal interactome is dysregulated in diabetic wounds. (A) Dot plot showing differential protein and RNA marker expressions for all identified neutrophil, Mo/MΦ and fibroblast subtypes in an integrated dataset. (B–C) Differential cell-cell communication analysis showing total interaction counts (B) and interaction strength (C) across immune and dermal cell subtypes in db/db vs Non-diabetic (ND) wounds. (D–F) Scatter plots of major pathways with differential incoming and outgoing interaction strengths for nα: *Tnfaip3^+^Sod2^+^*Ly6G^+^ neutrophils (D), mα: F4/80^high^CD11b^high^ Mo/MΦ (E) and mβ: Ly6c^high^CD11b^high^ Mo/MΦ (F). (G–I) Dot plots of up-regulated ligand-receptor pairs in db/db vs ND wounds for nα: *Tnfaip3^+^Sod2^+^*Ly6G^+^ neutrophils (G), mα: F4/80^high^CD11b^high^ Mo/MΦ (H), and mβ: Ly6c^high^CD11b^high^ Mo/MΦ (I). (J) Dot plot showing the specific expression of *Cd44* interactors across genotypes and neutrophil (bottom), Mo/MΦ (middle) and fibroblast (top) subtypes. (K) StringDB physical interaction network of top up-regulated ligands and receptors in db/db vs ND immune and dermal cell subtypes, highlighting top gene ontology (GO) enriched pathways as node border colors and identifying *Cd44* as a central communication hub (red edges).

Scatter plots graphing incoming versus outgoing interaction strengths across dermal subtypes showed that in ND wounds the nβ: Csf3r^+^Fos^+^Ly6G^+^ neutrophils, fβ: Lrrc15^+^Tnc^+^CD140a^high^ fibroblasts, and Proliferative FB (fibroblasts) were especially active; in db/db wounds the nα: Tnfaip3^+^Sod2^+^Ly6G^+^ neutrophils and mα: F4/80^high^CD11b^high^ Mo/MΦ were active; and in both genotypes mβ: Ly6c^high^CD11b^high^ Mo/MΦ and fγ: Cilp^+^Mgp^+^CD26^low^ fibroblasts were active **(Fig S8A-B)**.

We then performed differential CCC analyses **(Fig S8C)**, and we visualized the differential interaction counts **(Fig 5B)** and strengths **(Fig 5C)** between cell subtypes in db/db versus ND wounds via heatmaps, which revealed that in diabetic wounds there were major increases in outgoing interactions from nα: Tnfaip3^+^Sod2^+^Ly6G^+^ neutrophils and mα: F4/80^high^CD11b^high^ Mo/MΦ to nearly all other subtypes, whereas mβ: Ly6c^high^CD11b^high^ Mo/MΦ had increased interaction strengths against only these two other subtypes **(Fig S8D)**. We then mapped the differential incoming and outgoing signaling pathways in db/db vs ND wounds in nα: Tnfaip3^+^Sod2^+^Ly6G^+^ neutrophils **(Fig 5D)**, mα: F4/80^high^CD11b^high^ Mo/MΦ **(Fig 5E)**, and mβ: Ly6c^high^CD11b^high^ Mo/MΦ **(Fig 5F)**. In nα: Tnfaip3^+^Sod2^+^Ly6G^+^ neutrophils the CCL, CXCL, COLLAGEN, and SPP1 pathways were higher in db/db vs ND wounds; in mα: F4/80^high^CD11b^high^ Mo/MΦ the CCL, THBS, COLLAGEN, FN1, ApoE, APP, and SPP1 pathways were higher; and in mβ: Ly6c^high^CD11b^high^ Mo/MΦ only the ApoE and SPP1 pathways were higher. Differential signaling in the other immune **(Fig S8E)** and fibroblast **(Fig S8F)** subtypes was also mapped.

We performed DEG analyses to filter the CellChat-determined probable ligand-receptor interactions to those that had both the ligands (in sender cells) and receptors (in receiver cells) also significantly up-regulated **(see Methods, Table S19)**. We mapped these up-regulated ligand-receptor pairs for nα: Tnfaip3^+^Sod2^+^Ly6G^+^ neutrophils **(Fig 5G)**, mα: F4/80^high^CD11b^high^ Mo/MΦ **(Fig 5H)**, and mβ: Ly6c^high^CD11b^high^ Mo/MΦ **(Fig 5I)**. In all three subtypes, shared ligand-receptor pairs were identified, including several involving *Icam1*, *Spp1*, and *Cxcl1-3* as ligands originating from these cell subtypes and *Cd44*, *Cxcr2* and integrin complexes as receptors expressed in other receiver cell subtypes. Intriguingly, the *Spp1-Cd44* and *Spp1*-integrin interactions were up-regulated across nearly all cell subtypes, while the *Icam1* and *Cxcl1-3* interactions were mostly limited to the immune cells. Dot plots confirmed the differential expressions of the *Cd44*-associated pathway genes in immune and dermal subtypes in the diabetic condition **(Fig 5J)**.

We then used StringDB to create a physical protein-protein interaction network of all up-regulated ligand-receptor pairs in the diabetic wound cell subtypes **(Fig 5K)**, revealing that many are highly connected and centered around a hub of *Cd44* and its direct interactors - including *Spp1, Fn1, Vcam1, Icam1* and *Cxcr2* - and that this hub connects together gene modules that are enriched for cell-cell and cell-matrix adhesion (right) to those enriched for cytokine-mediated signaling pathways (left), with nearly all genes commonly enriched for cell migration.

Overall, these results suggest that T2DM causes major disruption of dermal and immune cell subtype communication in healing wounds, with Tnfaip3^+^Sod2^+^Ly6G^+^ neutrophils, F4/80^high^CD11b^high^ Mo/MΦ and Ly6c^high^CD11b^high^ Mo/MΦ CCCs being particularly dysregulated in diabetic wounds via ligand-receptor interactions centered around CD44 that implicate defects in innate immune cell migration during the early stages of diabetic wound healing.

### Validation of CD44-mediated immune cell dysregulation in diabetic cells and wounds

We harvested 3 DPW wounds from db/db and ND mice and performed flow cytometry analysis for major immune and dermal cells **(Fig S9A)**. This showed that db/db wounds had nearly a two-fold increase in neutrophils and Ly6C^+^ Mo/MΦ and almost half the number of F4/80^+^ Mo/MΦ compared to ND controls **(Fig 6A)**. Importantly, neutrophils, Ly6C^+^ Mo/MΦ, and fibroblasts all had significantly higher CD44^+^ cells in diabetic wounds **(Fig 6B, Fig S9B).**

**Figure 6.**
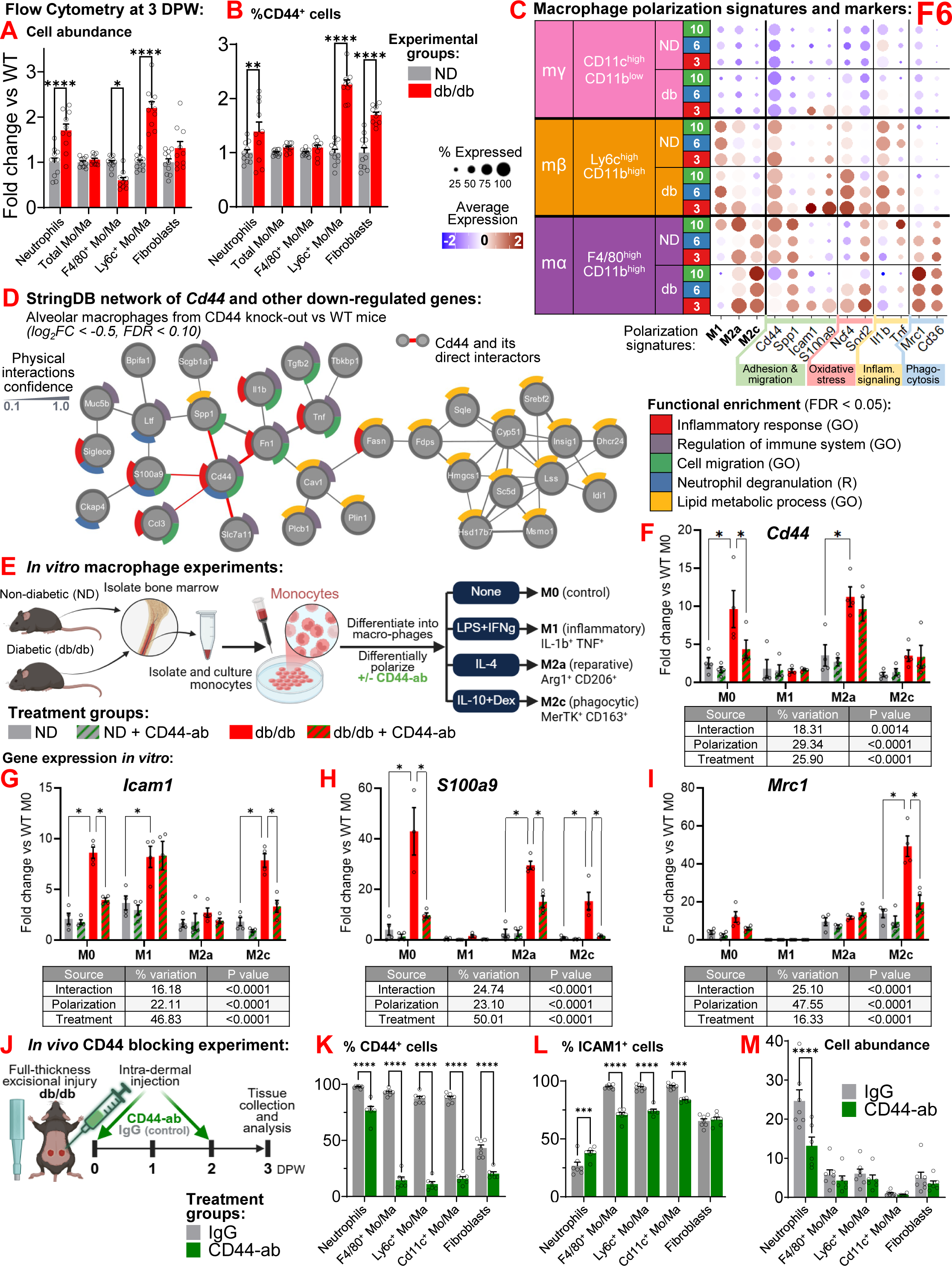
Validation of CD44 as a driver of diabetic immune cell dysfunction. (A–B) Flow cytometry analysis of db/db and non-diabetic (ND) wounds at 3 days post-wounding (DPW), showing relative cell abundance (A) and relative percentage of CD44^+^ cells (B) in db/db wounds compared to ND wounds. N=12 wounds per genotype; **** P<0.0001 and **P<0.01 by two-way ANOVA and Tukey’s post-hoc tests. (C) Dot plot of macrophage polarization signatures and associated gene markers in Mo/MΦ subtypes across all time points and genotypes. (D) StringDB protein-protein interaction network *Cd44* and other down-regulated genes in alveolar macrophages from CD44 knock-out (KO) mice, highlighting top gene ontology (GO) enriched pathways as node border colors and identifying *Cd44* as a central communication hub (red edges). (E–I) *In vitro* CD44 inhibition experiments using primary macrophages: (E) experimental schematic showing monocyte isolation from db/db and ND mice and their differential polarization into M0, M1, M2a and M2c phenotypes ± CD44 blocking antibody (CD44-ab) treatment; (F–I) fold change of *Cd44* (F), *Icam1* (G), *S100a9* (H), and *Mrc1* (I) in M0, M1, M2a, and M2c states ± CD44-blocking antibody, relative to ND M0. N=3-4 biological replicates; *P<0.05 by two-way ANOVA and Tukey’s post-hoc tests; results of two-way ANOVA shown under each graph. (J–M) *In vivo* CD44 blocking experiments in db/db mice: (J) experimental schematic showing full-thickness wounding and subsequent intradermal injection of CD44-ab or IgG at 0 and 2 DPW; flow cytometry analysis of db/db wounds at 3 DPW, showing percentage of CD44^+^ cells (K), percentage of ICAM1^+^ cells (L), and total cell abundance (M) following IgG or CD44-ab treatment. N=5-6 wounds per treatment; ****P<0.0001, ***P<0.001 by two-way ANOVA and Tukey’s post-hoc tests. All graphs show values as mean ± SEM.

**Figure 7.**
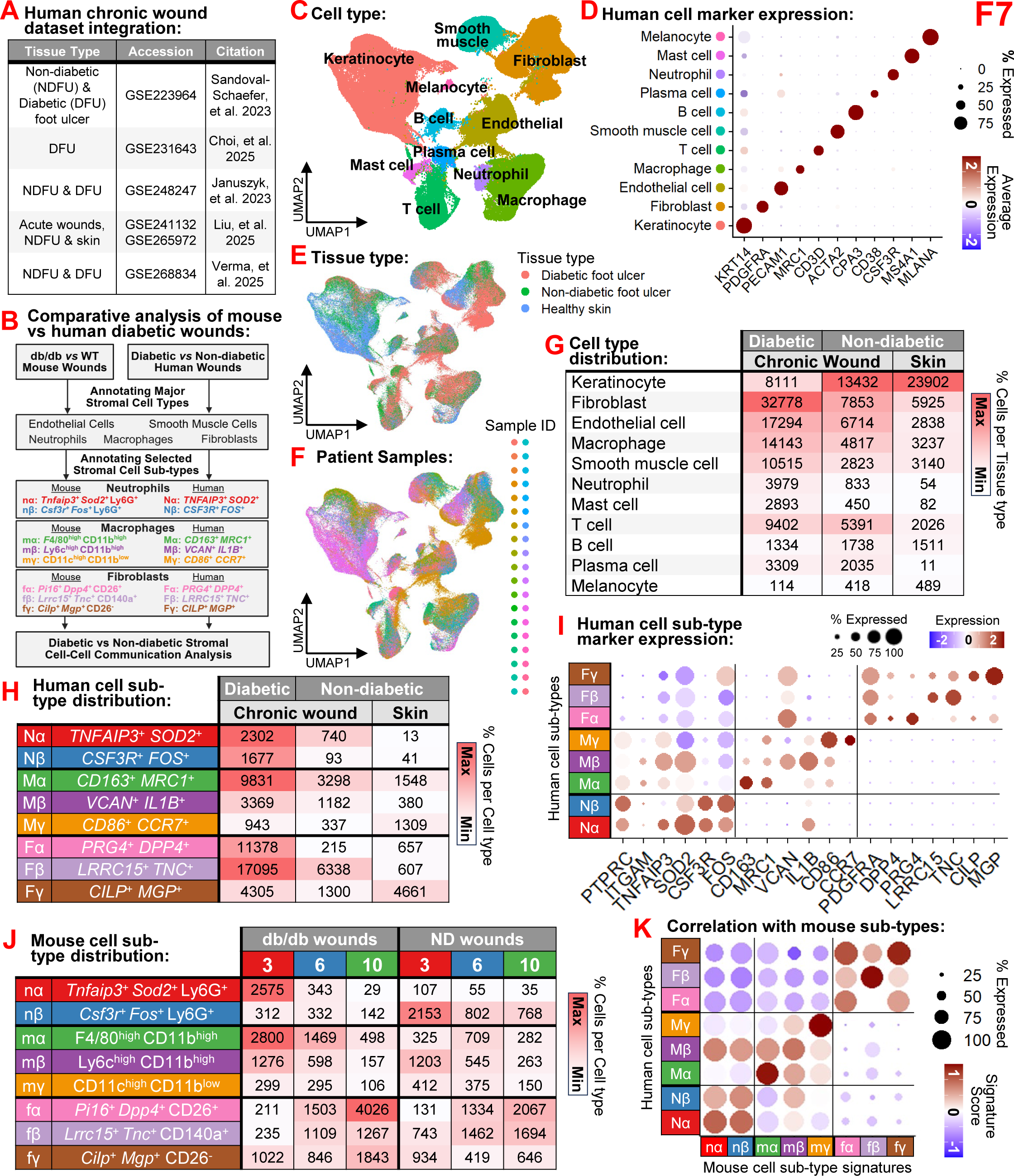
Integration and cross-species analysis of cells from human diabetic and non-diabetic foot ulcers. (A–B) Integration pipeline of human diabetic (DFU) and non-diabetic (ND-FU) foot ulcer datasets ^6–8,10,11^ (A) and workflow for comparative mouse-human immune and dermal cell subtype analysis (B). (C) UMAP visualization of 193,591 integrated human cells colored by major cell type annotation. (D) Dot plot showing the expressions of major cell type markers across annotated cell types. (E-F) UMAP visualizations of integrated human cells colored by tissue type (DFU, ND-FU, Healthy Skin) (E) and by patient sample ID (F). (G) Table of major cell type distributions in human diabetic vs non-diabetic conditions. (H) Table of human neutrophil, macrophage and fibroblast subtype distributions in human diabetic vs non-diabetic conditions. (I) Dot plot showing the expressions of differential human cell subtype markers. (J) Table of mouse neutrophil, macrophage and fibroblast subtype distributions across time points and diabetic vs non-diabetic conditions. (K) Dot plot comparing human cell subtype transcriptomes to humanized signatures of mouse cell subtypes.

Since flow cytometry confirmed the up-regulation of CD44^+^ immune cells in diabetic wounds but revealed a discrepancy in F4/80^+^ cells compared to our single-cell dataset **(Fig 3F)**, we focused on further deciphering the Mo/MΦ subtype data. We broke down the expression of macrophage polarization signatures (complementing data in **Fig 3H**) and associated gene markers of major macrophage phenotypes between the Mo/MΦ subtypes in db/db vs ND wounds across the three healing timepoints **(Fig 6C).** This confirmed the higher expression of *Cd44* in M1-like mβ: Ly6c^high^CD11b^high^ Mo/MΦ, which also expressed more inflammatory signaling (*Il1b, Tnf*) and oxidative stress (*Ncf4, Sod2*) genes, with the latter being higher in the diabetic condition. In contrast, the M2-like mα: F4/80^high^CD11b^high^ Mo/MΦ showed higher expression for *Spp1* and phagocytosis genes (*Mrc1, Cd36*), especially in the diabetic condition. Across all Mo/MΦ subtypes, adhesion and migration genes *Icam1* and *S100a9* were highly up-regulated in diabetic 3 DPW, pointing to major immune infiltration dysfunction during the early phase of diabetic wound healing.

To test how blocking of CD44 could influence macrophage gene expression, we re-analyzed a bulk RNA sequencing dataset of alveolar macrophages isolated from CD44 knock-out and ND mice ^27^. We used StringDB to create a protein-protein physical interaction network of all down-regulated genes, which revealed a well-connected hub of *Cd44* and its direct interactors, including *S100a9*, *Spp1* and *Fn1*, which also connected *Il1b* and *Tnf*, integrating modules of genes enriching for inflammation, neutrophil degranulation and cell migration **(Fig 6D)**.

Intriguingly, there was another module of down-regulated genes that highly enriched for lipid metabolism, possibly implicating *Cd44* not only in Mo/MΦ migration but also in diabetes-related metabolic dysfunction.

All bioinformatics analyses to this point led us to hypothesize that CD44 is causative in the immune cell dysfunction of T2DM-impaired wound healing, and that blocking CD44 may be an effective strategy to normalize immune cell behavior and lead to improved healing.

To directly test this hypothesis, we performed a series of *in vitro* experiments, wherein bone-marrow derived monocytes harvested from db/db and ND mice were first differentiated into macrophages and then differentially polarized into M0 (control), M1 (inflammatory), M2a (reparative) and M2c (phagocytic) phenotypes with or without pre-treatment with a CD44 blocking antibody (CD44-ab)^34^ **(Fig 6E)**. Gene expression analyses confirmed that M0 and M2a macrophages from diabetic mice had increased expression of *Cd44*, but that CD44-ab reduced its expression only in M0 macrophages **(Fig 6F)**. *Icam1* was significantly up-regulated in diabetic M0, M1 and M2c macrophages, and CD44-ab reduced its expression only in M0 and M2c macrophages **(Fig 6G)**. *S100a9* was significantly up-regulated in diabetic M0, M2a and M2c macrophages, and CD44-ab reduced its expression in all affected macrophages **(Fig 6H)**. *Mrc1* was significantly up-regulated and CD44-ab reduced its expression in diabetic M2c macrophages **(Fig 6I)**. *Ncf4* was significantly up-regulated in diabetic M0, M2a and M2c macrophages, and CD44-ab reduced its expression only in M0 macrophages **(Fig S9C)**, and *Spp1* showed no differences in expression amongst experimental groups **(Fig S9D)**. These experiments not only confirmed the phenotypic dysfunction of T2DM-affected Mo/MΦ but also showed how blocking CD44 could reverse the gene expression associated with these Mo/MΦ defects, including adhesion/migration (*Icam1*, *S100a9*), oxidative stress (*Ncf4*) and phagocytosis (*Mrc1*) in phenotype-specific conditions.

To test whether blocking CD44 could also be efficacious *in vivo*, we designed a mechanistic experiment to inhibit CD44 in healing wounds via intradermal injection of a blocking anti-CD44 antibody (CD44-ab)^34^ in db/db mice **(Fig 6J)**. Wounds were harvested at 3 DPW and subjected to flow cytometry **(Fig S9A)**, which confirmed a large reduction of CD44^+^ neutrophils, F4/80^+^ Mo/MΦ, Ly6C^+^ Mo/MΦ, CD11c^+^ Mo/MΦ, and fibroblasts in CD44-ab treated wounds compared to IgG controls **(Fig 6K, Fig S9E)**. ICAM1^+^ cells also decreased significantly amongst all Mo/MΦ subtypes after CD44-ab treatment, while ICAM1^+^ neutrophils increased **(Fig 6L, Fig S9E)**.

Importantly, total neutrophil numbers were halved in diabetic wounds treated with CD44-ab **(Fig 6M)**, effectively reducing their cell abundance back to ND levels in early wounds **(Fig 6A)**.

Though not statistically significant, CD44-ab treated wounds also trended towards faster closure at 2 and 3 DPW **(Fig S9F)**.

These mechanistic experiments showed that T2DM results in considerable dysregulation of innate immune cell infiltration and activation during the early stages of wound healing, that these phenotypes are mediated to a large extent by CD44 and ICAM1 leading to cell adhesion, migration and activation defects in neutrophils and Mo/MΦ, and that blocking CD44 may be an effective strategy to normalize this dysregulation of innate immune cells during diabetic wound healing.

### Re-analysis and integration of human diabetic chronic wound datasets

To test whether our findings from the db/db mouse model could be translated to human chronic wounds, we integrated six published single-cell sequencing datasets containing wound samples from diabetic (D) and non-diabetic (ND) foot ulcers (FU), acute wounds and healthy skin ^6–8,10,11^ **(Fig 7A)**. We devised a bioinformatics analysis pipeline to annotate human dermal wound cells and subtypes that parallels the one we performed for the db/db mouse samples, to enable direct inter-species comparisons **(Fig 7B)**. Following robust quality control, dataset integration, clustering and major cell annotation protocols **(see Methods, Fig S10-S11)**, we generated an integrated human single-cell dataset of 193,591 cells containing all expected major cell types **(Fig 7C)** expressing canonical cell type markers **(Fig 7D)**, across three major tissue types including D and ND chronic wounds and unwounded skin **(Fig 7E)**, obtained from 38 individual patients **(Fig 7F)**. We performed major cell type distribution analysis according to both disease (D vs ND) and tissue type (wound vs skin) annotations **(Fig 7G)**. Notably, DFUs had less keratinocytes but more neutrophils, macrophages and fibroblasts than NDFUs and ND skin.

We next subset the neutrophil, macrophage and fibroblast human cells, re-clustered them into subtypes and performed DEG analyses to determine their subtype gene markers **(see Methods, Fig S12)**. We characterized two neutrophil subtypes (Nα: identified by *TNFAIP3*^+^ *SOD2*^+^; Nβ: *CSF3R*^+^ *FOS*^+^), three macrophage subtypes (Mα: *CD163*^+^ *MRC1*^+^; Mβ: *VCAN*^+^ *IL1B*^+^; Mγ: *CD86*^+^ *CCR7*^+^) and three fibroblast subtypes (Fα: *PRG4*^+^ *DPP4*^+^; Fβ: *LRRC15*^+^ *TNC*^+^; Fγ: *CILP*^+^ *MGP*^+^), all represented in both D and ND FUs **(Fig 7H)** and across multiple patients **(Fig S11F)**. We integrated the single-cell datasets of the human cell subtypes, and a combined dot plot confirmed the shared and differential expressions of all subtype gene markers **(Fig 7I)**.

We mapped the integrated mouse dermal and immune cell subtypes across all the DPW in db/db and ND wounds **(Fig 7J)**, generated humanized gene lists of their cell, and compared the human subtype transcriptomes with mouse subtype signatures via module scoring **(Fig 7K)**.

This showed that most mouse neutrophil, fibroblast and macrophage subtypes correlated appreciably well with respective human dermal wound subtypes, especially nα-Nα, mα-Mα, mγ-Mγ, fβ-Fβ and fγ-Fγ, that Mβ showed some similarity to mouse neutrophils, and that there was some cross-correlation between the α and γ fibroblast subtypes.

### Cell-cell communication disruptions of human diabetic chronic wound cell subtypes

To test whether the CD44 interactors that were dysregulated in mouse diabetic wounds were also changed in human diabetic wounds, we visualized via dot plot the differential expressions of their human orthologs in human cell subtypes **(Fig 8A)**. This showed remarkable congruency with the mouse subtypes **(Fig 5J)**, including increased *CD44* and *ICAM1* expression in D vs ND neutrophil and macrophage subtypes, increased *SPP1* expression in macrophage subtypes, and increased *VCAM1* in fibroblast subtypes.

**Figure 8.**
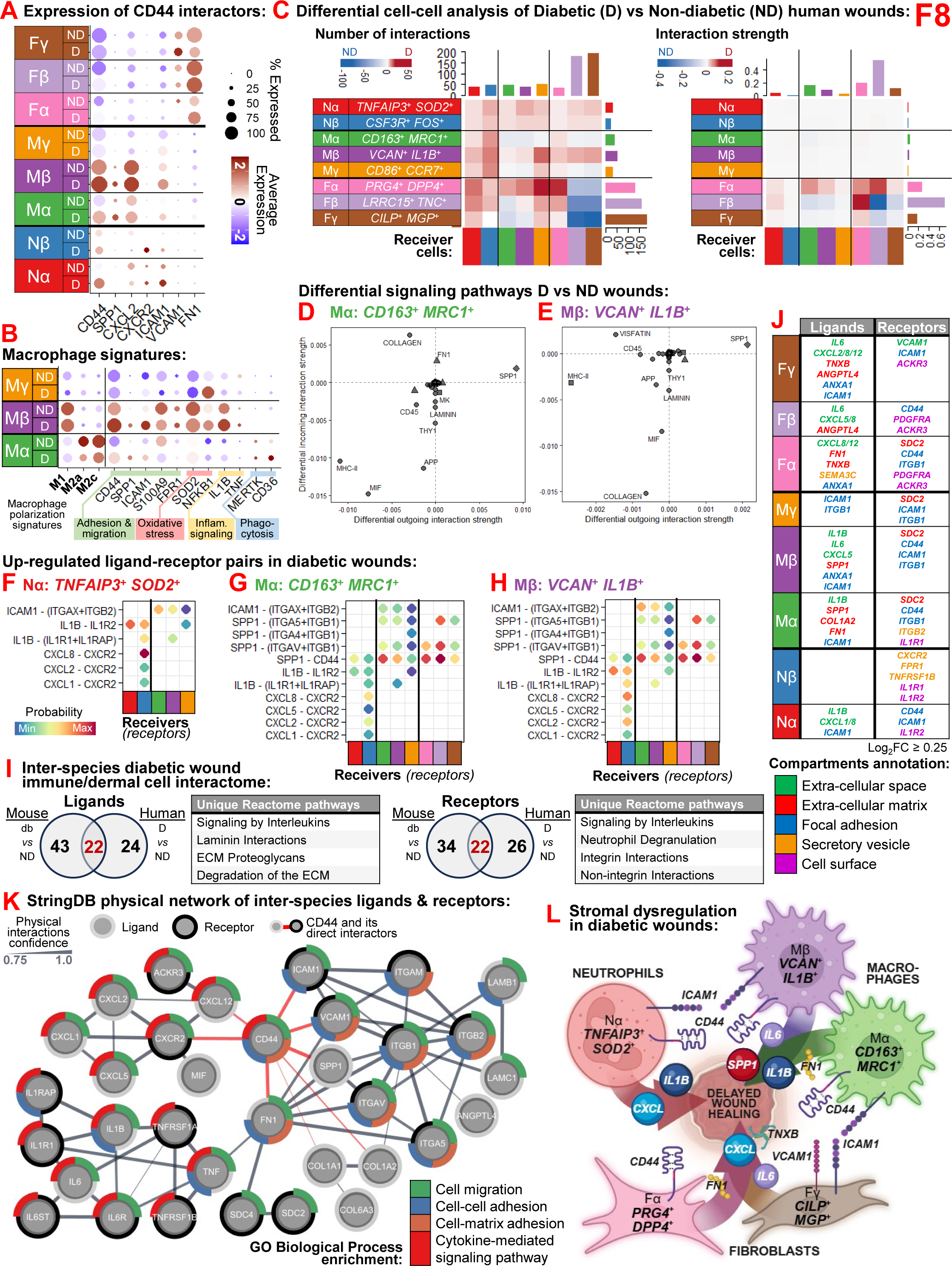
Conserved immune cell crosstalk disruptions and the CD44 hub in human chronic wounds. (A-B) Dot plots showing expressions of major CD44 interactors (A) and macrophage polarization signatures/markers (B) in human cell subtypes. (C) Differential cell-cell interaction analysis, showing relative interaction counts (left) and relative interaction strengths (right) across human cell subtypes in diabetic (D) vs non-diabetic (ND) human wounds. (D-E) Scatter plots of major pathways with differential incoming and outgoing interaction strengths for Mα: *CD163*^+^ *MRC1*^+^ (D) and Mβ: *VCAN*^+^ *IL1B*^+^ (E) macrophage subtypes. (F-H) Dot plots of up-regulated ligand-receptor pairs in D vs ND wounds for Nα: *TNFAIP3*^+^ *SOD2*^+^ neutrophils (F), Mα: *CD163*^+^ *MRC1*^+^ Mo/MΦ (G), and Mβ: *VCAN*^+^ *IL1B*^+^ Mo/MΦ (H). (I) Inter-species comparisons of up-regulated ligands (left) and receptors (right) in diabetic vs non-diabetic wounds of mice and humans. Venn diagrams identify 22 ligands and 22 receptors shared between mouse and human diabetic wounds, and tables show the top unique Reactome pathways enriched in the shared ligands/receptors. (J) Table listing the inter-species diabetic wound interactome according to the human cell subtype in which it is expressed and colored according to the Compartments annotation. (K) StringDB integrated physical network of inter-species ligands (black outline) and receptors (gray outline) centered on the CD44 hub (red edges), highlighting top gene ontology (GO) enriched pathways as node border colors. (L) Summary diagram of immune and dermal cell dysregulation in diabetic wounds, illustrating the key role of CD44 and ICAM1 in the disruption of wound cell crosstalk.

To compare the human macrophage transcriptomes to macrophage polarization signatures, we performed module scoring analyses with gene sets derived from the literature describing the gene expression signatures of macrophages polarized *in vitro* into the M1 (pro-inflammatory), M2a (anti-inflammatory) or M2c (phagocytic) phenotypes ^30^. The module scores of these signatures and associated gene markers of major macrophage phenotypes were analyzed in human macrophage subtypes from D and ND chronic wounds **(Fig 8B).** This showed that Mβ: *VCAN*^+^ *IL1B*^+^ macrophages enriched for the M1 polarization signature, whereas Mα: *CD163*^+^ *MRC1*^+^ macrophages enriched for M2 polarization signatures. Mβ: *VCAN*^+^ *IL1B*^+^ macrophages had higher expression of inflammatory signaling (*IL1B, TNF*) and oxidative stress (*SOD2, NFKB1*) genes, with both being higher in the diabetic condition. In contrast, the Mα: *CD163*^+^ *MRC1*^+^ macrophages showed higher expression for phagocytosis genes (*MERTK, CD36*), especially in the diabetic condition. Across all macrophage subtypes, adhesion and migration genes (*CD44*, *ICAM1*, *FPR1)* were up-regulated in the diabetic condition, pointing to major immune infiltration dysfunction during human diabetic wound healing.

We split the human subtype data into D and ND FU datasets and performed individual CCC analyses via CellChat **(Fig S12A-B)**. We then performed a differential CCC analysis **(Fig S12C)**, visualizing the differential interaction counts **(Fig 8C)** and strengths **(Fig 8D)** between cell subtypes in D versus ND wounds, which revealed that in D wounds there were major increases in outgoing and incoming interactions amongst most cell subtypes, except for fibroblast subtypes Fβ: *LRRC15*^+^ *TNC*^+^ and Fγ: *CILP*^+^ *MGP*^+^ **(Fig S12D)**. We mapped the differential incoming and outgoing signaling pathways in D versus ND wounds in all cell subtypes **(Fig S12E)**, revealing increases in the SPP1 pathways in Mα: *CD163*^+^ *MRC1*^+^ **(Fig 8D)** and Mβ: *VCAN*^+^ *IL1B*^+^ macrophages **(Fig 8E)**, consistent with the analysis of our mouse data **(Fig 5D-F)**.

We performed DEG analyses to filter the CellChat-determined probable ligand-receptor interactions to those that had both the ligands (in sender cells) and receptors (in receiver cells) also significantly up-regulated **(Table S20)**. We mapped these up-regulated ligand-receptor pairs for Nα: *TNFAIP3*^+^ *SOD2*^+^ neutrophils **(Fig 8F)**, Mα: *CD163*^+^ *MRC1*^+^ macrophages **(Fig 8G)** and Mβ: *VCAN*^+^ *IL1B*^+^ macrophages **(Fig 8H)**. In all subtypes, shared ligand-receptor pairs were identified, including several involving *ICAM1*, *SPP1*, *IL1B* and *CXCL* as ligands originating from these cell subtypes and *CD44*, *IL1R*, *CXCR2* and integrin complexes as receptors expressed in other receiver cell subtypes. As in the mouse subtypes **(Fig 5G-I)**, the *SPP1-CD44* and *SPP1*-integrin interactions were up-regulated across nearly all cell subtypes, while the *ICAM1* and *CXCL* interactions were mostly limited to the immune cells.

We used Venn diagrams to identify shared up-regulated ligands and receptors between the mouse and human subtype CCC analyses, leading to 22 common ligands and 22 common receptors **(Fig 8I, Table S21)**. The shared ligands enriched for unique Reactome pathways including Laminin Interactions, ECM Proteoglycans and ECM Degradation (left), the shared receptors enriched for Neutrophil Degranulation, Integrin and Non-integrin Interactions (right), and both were highly enriched for Signaling by Interleukins. These shared ligands and receptors were mapped according to which cell subtypes over-expressed them in human diabetic wounds, and they were annotated according to the curated Compartments database via EnrichR, which showed that in diabetic wound cell subtypes CD44 and ICAM1 are focal adhesion proteins that interact with integrins and extracellular matrix proteins SPP1 and FN1, with several cell subtypes also over-expressing inflammatory cytokines such as IL1B, IL6 and several CXCLs **(Fig 8J)**.

Finally, we built an inter-species diabetic wound interactome. We used StringDB to create a protein-protein physical interaction network of all inter-species shared up-regulated ligand-receptor pairs **(Fig 8K)**. This revealed that many genes are highly connected and centered around a hub of *CD44* and its direct interactors - including *SPP1, FN1 and ICAM1* - and that this hub connects together gene modules that are enriched for cell-cell and cell-matrix adhesion (right) and cytokine-mediated signaling pathways (left), all commonly enriched for cell migration.

Overall, these results suggest that T2DM causes major dermal and immune cell subtype disruptions in human chronic wounds that are mirrored in the db/db mouse model **(Fig 8L)**, with Nα:*TNFAIP3*^+^ *SOD2*^+^ neutrophils (analogous to the mouse nα:*Tnfaip3^+^ Sod2^+^*Ly6G^+^ neutrophils), Mα:*CD163*^+^ *MRC1*^+^ macrophages (analogous to the mouse mα: F4/80^high^ CD11b^high^ Mo/MΦ) and Mβ: *VCAN*^+^ *IL1B*^+^ macrophages (analogous to the mouse mβ: Ly6c^high^ CD11b^high^ Mo/MΦ) CCCs being particularly dysregulated in diabetic compared to non-diabetic wounds via ligand-receptor interactions centered around *CD44* and *ICAM1*.

## DISCUSSION

The failure of many diabetic wounds to transition from inflammation to proliferation and successful resolution represents a significant clinical challenge ^1,2^. Our multi-omic analysis provides a high-resolution atlas of this failure, revealing that the T2DM-affected wound microenvironment is characterized by profound temporal dysregulations of dermal and immune cell abundances and cell-cell communication networks. Central to this dysfunction is the CD44-mediated interactome, which acts as a hub for aberrant adhesion, migration and inflammatory pathways across fibroblasts, macrophages and neutrophils. By integrating mouse and human data, we demonstrate that these cellular states are conserved across species, providing a unified framework for understanding diabetic wound chronicity at the early stages of repair.

An important finding of our study is the identification of the nα: *Tnfaip3^+^ Sod2^+^* Ly6G^+^ neutrophil and mα: F4/80^high^ CD11b^high^ macrophage subtypes as the primary drivers of early immune dysfunction in mouse and human diabetic wounds. The nα: *Tnfaip3^+^ Sod2^+^* Ly6G^+^ subtype exhibits signatures associated with neutrophil degranulation, apoptosis and TNF signaling, aligning with studies that showed that diabetes primes neutrophils for NETosis ^35–37^. These findings suggest that T2DM traps neutrophils in a chronic inflammatory state that prevents their timely resolution.

Similarly, the diabetic mα: F4/80^high^ CD11b^high^ macrophage population is skewed toward a dysfunctional phagocytic and oxidative stress phenotype. This aligns with the recently-described "common framework" of macrophage activation, where metabolic stress polarizes cells toward dysfunctional rather than restorative phagocytic states ^31^. Notably, while previous paradigms suggested that M1-like (pro-inflammatory) macrophages are the primary bad actors in diabetic wounds ^38,39^, recent studies of human diabetic foot ulcers have shown that healing wounds actually exhibit increased M1-like polarization and non-healing wounds are dominated by dysfunctional M2-like signatures ^2,5,9^. Our *in vivo* data makes clear that wound macrophages are highly heterogeneous and their classifications in complex microenvironments need more nuance than the classical M1/M2 dichotomy, which is useful for *in vitro* studies under controlled conditions but falls short of describing the many possible functional states of macrophages ^30,31^.

In the dermal compartment, the loss of fβ: *Lrrc15*^+^ *Tnc*^+^ CD140a^high^ fibroblasts, which we previously identified as an early phase-specific fibroblast signature that enriched for collagen production and contraction ^3,40^, may explain the delays in connective tissue formation and mechanical closure in diabetic wounds. This subtype partly matches the healing-associated fibroblasts described in human healing wounds, which overexpress matrix extracellular matrix remodeling factors ^9^. Our analysis further links these to the stable scar states recently mapped in murine wound atlases ^28^. Conversely, the expansion of fγ: *Cilp*^+^ *Mgp*^+^ CD26^low^ fibroblasts in late-stage diabetic wounds, characterized by adipogenesis and lipid transport signatures, suggests a pathological metabolic reprogramming of the wound granulation tissue that may prevent proper reconstruction and resolution.

Our results identify CD44 and its downstream interactor ICAM1 as key mediators of the immune cell dysfunction in diabetic wounds. While CD44 is known to play roles in leukocyte recruitment and metabolic homeostasis ^27,34,41–44^, we show here that its up-regulation in diabetic wounds directly correlates with excessive neutrophil influx and defective macrophage activation.

Mechanistically, blocking CD44 not only normalized the neutrophil-to-macrophage ratio in early diabetic wounds but also inhibited the expression of *Icam1* and *S100a9* in primary macrophages, thereby attenuating their adhesion and oxidative stress defects. This therapeutic potential is further bolstered by our human meta-analysis, which confirmed the up-regulation of the CD44-ICAM1 axis in human neutrophil, macrophage and fibroblast subtypes in diabetic compared to non-diabetic chronic wounds.

While the db/db mouse model recapitulates many systemic metabolic effects of T2DM, the wound healing outcomes are less severe compared to the human condition: db/db skin healing is delayed, not chronically impaired ^12–14^. However, our db/db wound healing atlas shows major parallels between the cellular and molecular microenvironments of the mouse early inflammatory wound and the human diabetic chronic wound, pointing to its translational potential for modeling this stalled inflammatory state. Future research should build upon this baseline by incorporating variables such as oxidative stress, biofilm and mechanical strain to better understand their contributions to increased chronicity in the db/db model ^45–48^. Future studies will also systematically compare our db/db wound atlas to other delayed healing models, such as the high-fat diet and streptozocin-induced diabetes models, to create a unified understanding of their pathophysiology and appropriate uses for translational research ^48–50^.

In conclusion, our study provides a systems-level view of the cellular and molecular parallels between early-phase mouse and human diabetic wounds. We show that T2DM causes a major shift in the dermal and immune cell interactome, moving away from reparative fibroblast-mediated signals toward a dysregulated, CD44-centered immune-cell crosstalk that may explain the well-documented persistence of NETotic neutrophils and phagocytic macrophages in diabetic wounds. These findings pave the way for the development of CD44-targeted therapies that could potentially normalize the inflammatory microenvironment and restore healing potential in patients with T2DM. Future studies should explore the spatial distribution of these CD44-rich niches, including their impact on the extracellular matrix, to better understand the biomechanical and metabolic forces that maintain chronicity in the diabetic granulation tissue.

## SUPPLEMENTARY FIGURE LEGENDS

**Figure S1. Quality control, clustering, and protein decomplexing of integrated mouse wound cells.**

(A) Quality control scatter plots showing UMI counts (nCount_RNA), detected genes (nFeature_RNA), and percent mitochondrial genes across Day 3, 6, and 10 post-wounding for non-diabetic (ND) and diabetic (db/db) mice prior to filtration. (B) Elbow plot showing the standard deviation of the first 200 principal components (PCs) utilized to select optimal dimensionality for downstream analyses. (C) UMAP visualization of the final 12 Seurat-determined cell clusters. (D) UMAP visualizations of integrated cells split by genotype (ND vs db/db) and harvest time point, demonstrating no severe batch effects across technical replicates. (E) UMAP visualizations colored by the top-scoring antibody-derived surface proteins (Protein_maxID, top) and the second top-scoring surface proteins (Protein_secondID, bottom) derived from HTODemux decomplexing.

**Figure S2. Quality control and expression profiling of major cell populations.**

(A) Dot plots correlating the targeted cell markers’ protein expressions and their corresponding RNA expressions across cells grouped by their Protein_maxID (left) and Protein_secondID (right). (B) Dot plot visualizing the protein and RNA expressions of top cell markers across the 12 unsupervised Seurat-identified clusters. (C) Heatmap of the top differentially expressed proteins (DEP) identifying each initial Seurat cluster. (D) Methodological workflow for identifying and annotating major cell clusters based on the sequential presence or absence of key immune and non-immune protein markers. (E) Top DEPs confirming the proteins used to annotate the major cell types. (F) Dot plot showing the concordance between average surface protein expression and corresponding RNA marker expression across all annotated major cell types. (G) Violin plots showing the distributions of UMI counts (nCount_RNA) and detected features (nFeature_RNA) grouped according to the major cell annotations to ensure consistent data quality across cell types.

**Figure S3. Differentially-expressed gene analysis of Seurat clusters.**

Tables detailing the top differentially expressed genes (DEGs) calculated for each initial Seurat cluster (0-11). The top 20 genes expressed in at least 75% of the cells within the given cluster are presented. The table lists the Log_2_ fold change (Log_2_FC), percent of positive cells in the target cluster (% C pos), percent of positive cells outside the target cluster (% NC pos), and False Discovery Rate (FDR).

**Figure S4. Differentially expressed gene analysis of annotated major cells.**

Tables detailing the top differentially expressed genes (DEGs) calculated for each annotated major cell population (Fibroblasts, Monocytes/Macrophages, Neutrophils, Endothelial cells, Smooth muscle cells, T cells/NK cells, Muscle progenitor cells, and Skeletal muscle cells). The top 30 genes expressed in at least 75% of the cells within the major cell type are presented. The table lists the Log_2_ fold change (Log_2_FC), percent of positive cells in the target cluster (% C pos), percent of positive cells outside the target cluster (% NC pos), and False Discovery Rate (FDR).

**Figure S5. Protein-RNA expression concordance in the integrated single-cell dataset.**

(A) Color key for the major cell type annotations utilized in the scatter plots. (B-R) Paired UMAP feature plots and scatter plots demonstrating the expression patterns of top cell markers at both the protein (left UMAP, x-axis on scatter plot) and RNA (right UMAP, y-axis on scatter plot) levels. Markers shown include: CD45/*Ptprc* (B), CD11b/*Itgam* (C), F4/80/*Adgre1* (D), Ly6C/*Ly6c2* (E), CD11c/*Itgax* (F), Ly6G/*Ly6g* (G), CD31/*Pecam1* (H), CD26/*Dpp4* (I), CD140a/*Pdgfra* (J), CD3e/*Cd3e* (K), CD4/*Cd4* (L), NK1.1/*Klrb1c* (M), CD335/*Ncr1* (N), CD202b/*Tek* (O), CD117/*Kit* (P), CD115/*Csf1r* (Q), CD326/*Epcam* (R).

**Figure S6. Cell-cell communication signaling changes between major cell types in diabetic wounds.** (A) CellChat scatter plots showing the overall outgoing and incoming interaction strength of major cell types in non-diabetic (ND, left) and diabetic (db/db, right) wounds at 3 days post-wounding. (B) Circle plots visualizing the absolute number of interactions (top) and the interaction strength network (bottom) between major cell types in ND (left) and db/db (right) wounds. (C) Bar graphs comparing total inferred interactions and overall interaction strengths in ND vs db/db wounds. (D) Differential circle plots showing relative changes in the number of interactions (top) and interaction strength (bottom) between major cell types (db/db vs ND), where red indicates increased signaling in db/db wounds and blue indicates decreased signaling. (E) Heatmap of the differential number of interactions among specific major cell sender and receiver groups. (F) Scatter plots mapping the differential incoming and outgoing interaction strengths of individual signaling pathways in neutrophils, macrophages, fibroblasts, smooth muscle cells and endothelial cells (db/db vs ND).

Figure S7. **Identification and functional enrichment of proliferative fibroblast and macrophage subtypes.**

(A) Elbow plots utilized to determine the appropriate number of principal components (PCs) for the isolated sub-clustering of fibroblasts, monocytes/macrophages (Mo/MΦ) and neutrophils. (B) Table showing the top differentially expressed genes (DEGs) alongside Reactome pathway functional enrichment results confirming the identity of the fδ: Proliferative fibroblast (FB) subtype. (C)*Table showing the top DEGs alongside Reactome pathway functional enrichment results confirming the identity of the mδ: Proliferative Mo/MΦ subtype.

**Figure S8. Cell-cell communication signaling changes between cell sub-types in diabetic wounds.** (A) CellChat scatter plots mapping the overall outgoing and incoming interaction strengths among all isolated cell subtypes in non-diabetic (ND, left) and diabetic (db/db, right) wounds. (B) Circle plots visualizing the network of interaction counts (top) and total interaction strength (bottom) between all cell subtypes in ND (left) and db/db (right) wounds. (C) Bar graphs comparing total inferred interactions and total interaction strength across all subtypes in ND vs db/db wounds. (D) Differential circle plots showing relative changes in the number of interactions (top) and interaction strength (bottom) between individual cell subtypes (db/db vs ND), where red indicates increased signaling in db/db wounds. (E-F) Scatter plots mapping specific differential incoming and outgoing interaction pathway strengths across individual immune cell subtypes (E) and fibroblast subtypes (F) in db/db vs ND wounds.

**Figure S9. Flow cytometry and *in vitro* analyses of macrophage polarization markers.**

(A) Flow cytometry sequential gating strategy utilizing Zombie viability dye, CD140a, CD11b, Ly6G, F4/80, Ly6C, and CD11c to isolate fibroblasts, specific monocytes/macrophages (Mo/Ma) and neutrophils from single-cell skin wound suspensions. (B) Representative flow cytometry histogram showing baseline CD44 surface expression on dermal cells from non-diabetic (ND) and diabetic (DB) wounds compared to unstained controls. (C) Additional *in vitro* RT-qPCR gene expression analyses for *Ncf4* (C) and *Spp1* (D) of primary bone marrow-derived macrophages from db/db and WT mice differentially polarized into M0, M1, M2a, and M2c functional states and treated with or without a blocking CD44 antibody (CD44-ab). N=3-4 biological replicates;= *P<0.05 by two-way ANOVA and Tukey’s post-hoc tests; results of two-way ANOVA shown under each graph. (E) Representative flow cytometry histograms demonstrating specific surface expression shifts of CD44 and ICAM-1 in db/db wounds treated via intradermal injection with either CD44-blocking antibody (CD44-Ab) or IgG negative control. (F) Percent wound closure as measured by external wound morphometry at 2 and 3 days post-wounding (DPW). N=5-6 wounds per treatment. All graphs show values as mean ± SEM.

**Figure S10. Re-analysis and cell-typing of independent human chronic wound and skin datasets.**

(A-F) Individual re-analyses and normalization of previously published human single-cell transcriptomic datasets: GSE223964 (A), GSE231643 (B), GSE241132 (C), GSE248247 (D), GSE265972 (E), and GSE268834 (F). Each panel displays UMAP visualizations of the specific dataset colored by wound condition/tissue type (left) and annotated cell types (middle). Dot plots (right) validate the cell annotations by confirming the expression of key cell type markers across annotated cell clusters.

**Figure S11. Integration of human single-cell datasets and cell sub-type distributions.**

(A-B) UMAP visualizations of the un-integrated, merged human datasets colored by the original dataset source/tissue type (A) and by individual sample ID (B), revealing considerable batch effects prior to harmonization. (C-D) UMAP visualizations of the final RPCA-integrated human dataset colored by tissue type (C) and individual sample ID (D), demonstrating successful cell integration and algorithmic batch correction. (E) Distribution table showing the absolute cell counts for major cell types mapped across all sample tissue types and dataset sources within the final integrated dataset. (F) Distribution table showing the absolute cell counts for the individual human neutrophil, macrophage and fibroblast subtypes calculated across all samples in the final human dataset.

**Figure S12. Sub-clustering of human dermal fibroblasts, macrophages, and neutrophils.**

(A) Elbow plots utilized to determine the appropriate number of principal components (PCs) for the isolated sub-clustering of human fibroblasts, macrophages and neutrophils. (B-D) UMAP visualizations of the subsetted and sub-clustered human fibroblasts (B), macrophages (C) and neutrophils (D). For each major cell subset, UMAPs are shown colored by source wound type (left), Seurat-determined sub-clusters (middle), and the final annotated cell subtypes (right).

**Figure S13. Cell-cell communication signaling changes between cell sub-types in human diabetic vs non-diabetic wounds.**

(A) CellChat scatter plots showing overall outgoing and incoming interaction strengths of human cell subtypes in non-diabetic (ND, left) and diabetic (D, right) foot ulcers. (B) Circle plots visualizing the interaction count networks (top) and interaction strength networks (bottom) between human subtypes in ND (left) and D (right) wounds. (C) Bar graphs detailing the total inferred interactions and interaction strengths across subtypes in ND vs D human foot ulcers. (D) Differential circle plots mapping the relative changes in the number of interactions (top) and interaction strength (bottom) between cell sub-types (D vs ND), where red nodes/edges indicate significantly increased signaling in diabetic conditions. (E) Additional scatter plots mapping the differential incoming and outgoing interaction strengths of specific individual signaling pathways across the Nα, Nβ, Mγ, Fα, Fβ, and Fγ human subtypes in D vs ND wounds.

## ACKNOWLEDGMENTS

The authors wish to thank Karla Herrera, Kartik Dixit, Siri Pothula, and Dr. Nathaniel Henning for valuable feedback that contributed to this project.

## REFERENCES

1. Armstrong, D. G., Tan, T.-W., Boulton, A. J. M. & Bus, S. A. Diabetic Foot Ulcers: A Review. JAMA 330, 62 (2023).

2. Hsia, H. C. et al. Management of Diabetic Wounds: Expert Panel Consensus Statement. Adv Wound Care (New Rochelle) 21621918251366586 (2025) doi:10.1177/21621918251366586.

3. Wietecha, M. S. et al. Phase-specific signatures of wound fibroblasts and matrix patterns define cancer-associated fibroblast subtypes. Matrix Biol 119, 19–56 (2023).

4. Rodrigues, M., Kosaric, N., Bonham, C. A. & Gurtner, G. C. Wound Healing: A Cellular Perspective. Physiol Rev 99, 665–706 (2019).

5. Clayton, S. M., Shafikhani, S. H. & Soulika, A. M. Macrophage and Neutrophil Dysfunction in Diabetic Wounds. Adv Wound Care (New Rochelle) 13, 463–484 (2024).

6. Januszyk, M. et al. Characterization of Diabetic and Non-Diabetic Foot Ulcers Using Single-Cell RNA-Sequencing. Micromachines (Basel) 11, 815 (2020).

7. Choi, D. et al. Single-Cell Analysis of Debrided Diabetic Foot Ulcers Reveals Dysregulated Wound Healing Environment in Non-Hispanic Black Patients. J Invest Dermatol 145, 678–690 (2025).

8. Sandoval-Schaefer, T. et al. Transcriptional heterogeneity in human diabetic foot wounds. bioRxiv 2023.02.16.528839 (2023) doi:10.1101/2023.02.16.528839.

9. Theocharidis, G. et al. Single cell transcriptomic landscape of diabetic foot ulcers. Nat Commun 13, 181 (2022).

10. Liu, Z. et al. Spatiotemporal single-cell roadmap of human skin wound healing. Cell Stem Cell 32, 479–498.e8 (2025).

11. Verma, S. S. et al. Tissue nanotransfection-based endothelial PLCγ2-targeted epigenetic gene editing rescues perfusion and diabetic ischemic wound healing. Mol Ther 33, 950–969 (2025).

12. Saeed, S. & Martins-Green, M. Assessing Animal Models to Study Impaired and Chronic Wounds. Int J Mol Sci 25, 3837 (2024).

13. Suriano, F. et al. Novel insights into the genetically obese (ob/ob) and diabetic (db/db) mice: two sides of the same coin. Microbiome 9, 147 (2021).

14. Ojeh, N., et al. The Wound Reporting in Animal and Human Preclinical Studies (WRAHPS) Guidelines. Wound Repair Regen 33, e13232 (2025).

15. Pang, J., Maienschein-Cline, M. & Koh, T. J. Reduced apoptosis of monocytes and macrophages is associated with their persistence in wounds of diabetic mice. Cytokine 142, 155516 (2021).

16. Pang, J., Urao, N. & Koh, T. J. Diet-Induced Obesity Increases Monocyte/Macrophage Proliferation during Skin Wound Healing in Mice. Cells 13, 401 (2024).

17. Pang, J., Maienschein-Cline, M. & Koh, T. J. Enhanced Proliferation of Ly6C+ Monocytes/Macrophages Contributes to Chronic Inflammation in Skin Wounds of Diabetic Mice. J Immunol 206, 621–630 (2021).

18. Joshi, N. et al. Comprehensive characterization of myeloid cells during wound healing in healthy and healing-impaired diabetic mice. Eur J Immunol 50, 1335–1349 (2020).

19. Keiser, S., Botello, N., Cruz, E. & Wietecha, M. S. Using R, Seurat, and CellChat to Analyze a Single-Cell Transcriptomics Dataset of Mouse Skin Wound Healing. J Vis Exp https://doi.org/10.3791/67266 (2025) doi:10.3791/67266.

20. Hao, Y. et al. Dictionary learning for integrative, multimodal and scalable single-cell analysis. Nat Biotechnol 42, 293–304 (2024).

21. Germain, P.-L., Lun, A., Garcia Meixide, C., Macnair, W. & Robinson, M. D. Doublet identification in single-cell sequencing data using scDblFinder. F1000Res 10, 979 (2021).

22. Jin, S., Plikus, M. V. & Nie, Q. CellChat for systematic analysis of cell–cell communication from single-cell transcriptomics. Nat Protoc https://doi.org/10.1038/s41596-024-01045-4 (2024) doi:10.1038/s41596-024-01045-4.

23. Jin, S. et al. Inference and analysis of cell-cell communication using CellChat. Nat Commun 12, 1088 (2021).

24. Doncheva, N. T., Morris, J. H., Gorodkin, J. & Jensen, L. J. Cytoscape StringApp: Network Analysis and Visualization of Proteomics Data. J Proteome Res 18, 623–632 (2019).

25. Szklarczyk, D. et al. The STRING database in 2023: protein–protein association networks and functional enrichment analyses for any sequenced genome of interest. Nucleic Acids Research 51, D638–D646 (2023).

26. Stoeckius, M. et al. Cell Hashing with barcoded antibodies enables multiplexing and doublet detection for single cell genomics. Genome Biol 19, 224 (2018).

27. Dong, Y. et al. CD44 Loss Disrupts Lung Lipid Surfactant Homeostasis and Exacerbates Oxidized Lipid-Induced Lung Inflammation. Front Immunol 11, 29 (2020).

28. Almet, A. A., Liu, Y., Nie, Q. & Plikus, M. V. Integrated Single-Cell Analysis Reveals Spatially and Temporally Dynamic Heterogeneity in Fibroblast States during Wound Healing. J Invest Dermatol 145, 645–659.e25 (2025).

29. Zandigohar, M. et al. Transcription Factor Activity Regulating Macrophage Heterogeneity during Skin Wound Healing. J Immunol 213, 506–518 (2024).

30. Strizova, Z. et al. M1/M2 macrophages and their overlaps - myth or reality? Clin Sci (Lond) 137, 1067–1093 (2023).

31. Sanin, D. E., et al. A common framework of monocyte-derived macrophage activation. Sci Immunol 7, eabl7482 (2022).

32. Mihaila, A. C. et al. Transcriptional Profiling and Functional Analysis of N1/N2 Neutrophils Reveal an Immunomodulatory Effect of S100A9-Blockade on the Pro-Inflammatory N1 Subpopulation. Front Immunol 12, 708770 (2021).

33. Grieshaber-Bouyer, R. et al. The neutrotime transcriptional signature defines a single continuum of neutrophils across biological compartments. Nat Commun 12, 2856 (2021).

34. Kodama, K., Toda, K., Morinaga, S., Yamada, S. & Butte, A. J. Anti-CD44 antibody treatment lowers hyperglycemia and improves insulin resistance, adipose inflammation, and hepatic steatosis in diet-induced obese mice. Diabetes 64, 867–875 (2015).

35. Fadini, G. P. et al. NETosis Delays Diabetic Wound Healing in Mice and Humans. Diabetes 65, 1061– 1071 (2016).

36. Shrestha, S. et al. Diabetes Primes Neutrophils for Neutrophil Extracellular Trap Formation through Trained Immunity. Research 7, 0365 (2024).

37. Wong, S. L. et al. Diabetes primes neutrophils to undergo NETosis, which impairs wound healing. Nat Med 21, 815–819 (2015).

38. Sharifiaghdam, M. et al. Macrophages as a therapeutic target to promote diabetic wound healing. Mol Ther 30, 2891–2908 (2022).

39. Barman, P. K. & Koh, T. J. Macrophage Dysregulation and Impaired Skin Wound Healing in Diabetes. Front Cell Dev Biol 8, 528 (2020).

40. Wietecha, M. S. et al. Activin-mediated alterations of the fibroblast transcriptome and matrisome control the biomechanical properties of skin wounds. Nat Commun 11, 2604 (2020).

41. Kodama, K. et al. Expression-based genome-wide association study links the receptor CD44 in adipose tissue with type 2 diabetes. Proc Natl Acad Sci U S A 109, 7049–7054 (2012).

42. Hasib, A. et al. CD44 contributes to hyaluronan-mediated insulin resistance in skeletal muscle of high-fat-fed C57BL/6 mice. Am J Physiol Endocrinol Metab 317, E973–E983 (2019).

43. Liu, L. F. et al. The receptor CD44 is associated with systemic insulin resistance and proinflammatory macrophages in human adipose tissue. Diabetologia 58, 1579–1586 (2015).

44. Weiss, L. et al. Induction of resistance to diabetes in non-obese diabetic mice by targeting CD44 with a specific monoclonal antibody. Proc Natl Acad Sci U S A 97, 285–290 (2000).

45. Kim, J. H. et al. High Levels of Oxidative Stress Create a Microenvironment That Significantly Decreases the Diversity of the Microbiota in Diabetic Chronic Wounds and Promotes Biofilm Formation. Front Cell Infect Microbiol 10, 259 (2020).

46. Jabbari, P. et al. Chronic Wound Initiation: Single-Cell RNAseq of Cutaneous Wound Tissue and Contributions of Oxidative Stress to Initiation of Chronicity. Antioxidants (Basel) 14, 214 (2025).

47. Zhao, G. et al. Time course study of delayed wound healing in a biofilm-challenged diabetic mouse model. Wound Repair Regen 20, 342–352 (2012).

48. Berryman, K. S. et al. Mechanotransduction unifies healthy non-diabetic wound healing over time by promoting a Cd14+/C1qa+ fibroblast subpopulation. J Invest Dermatol S0022–202X(26)01035–3 (2026) doi:10.1016/j.jid.2026.02.027.

49. Ma, J. et al. Single-cell RNA-Seq analysis of diabetic wound macrophages in STZ-induced mice. J Cell Commun Signal 17, 103–120 (2023).

50. Xie, Y., Yang, J., Zhu, H., Yang, R. & Fan, Y. The efferocytosis dilemma: how neutrophil extracellular traps and PI3K/Rac1 complicate diabetic wound healing. Cell Commun Signal 23, 103 (2025).

