## Supplemental Figures S1-S13 for "Multi-omics analysis of type II diabetic wound healing reveals CD44-mediated immune cell crosstalk dysfunction in mice and humans"

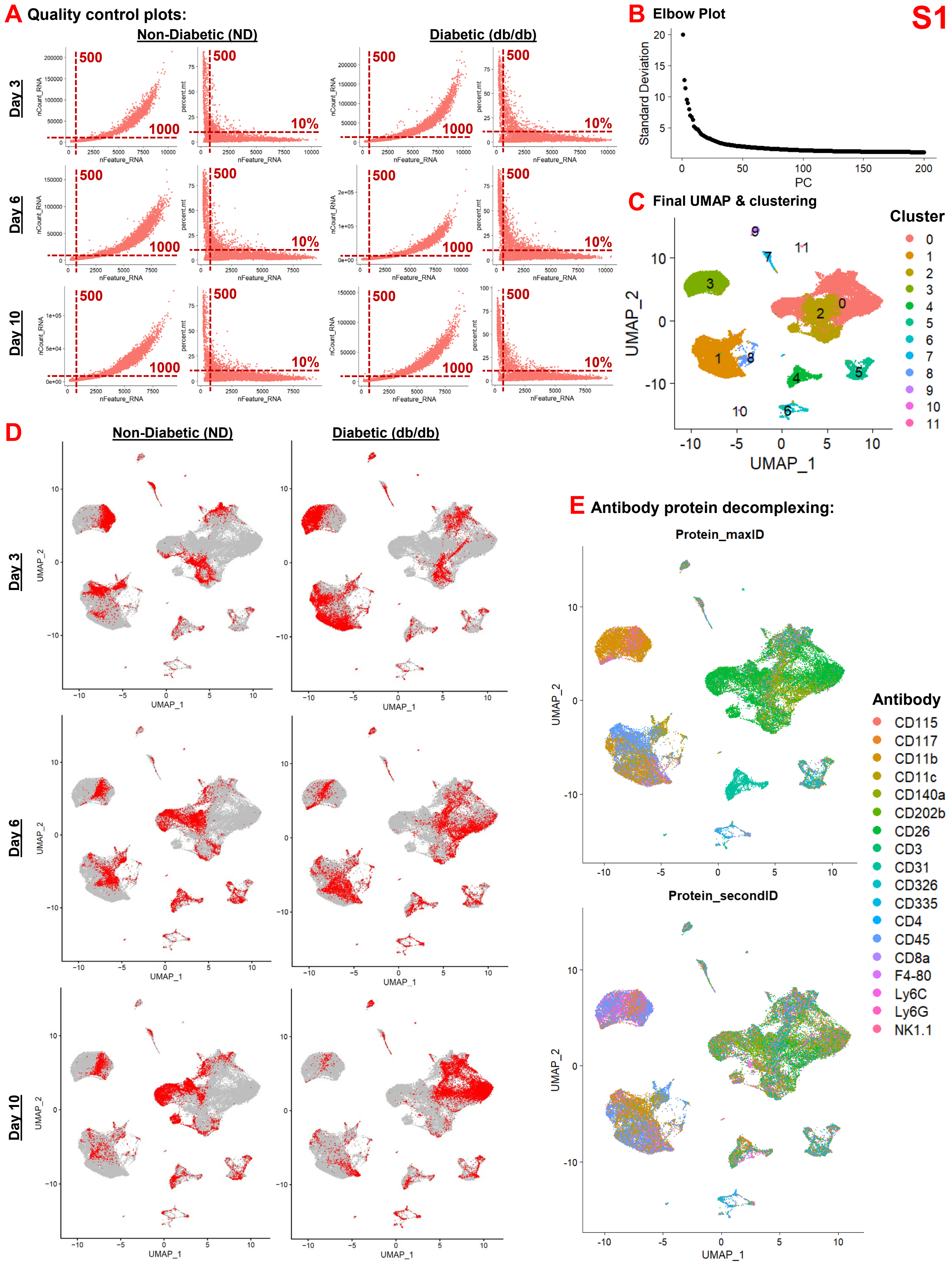
S1
A Quality control plots:
Non-Diabetic (ND)
Diabetic (db/db)
Day 3
Day 6
Day 10
500
1000
10%
nCount_RNA
percent.mt
nFeature_RNA
B Elbow Plot
Standard Deviation
PC
C Final UMAP & clustering
Cluster
UMAP_1
UMAP_2
D
E Antibody protein decomplexing:
Protein_maxID
Protein_secondID
Antibody
CD115
CD117
CD11b
CD11c
CD140a
CD202b
CD26
CD3
CD31
CD326
CD335
CD4
CD45
CD8a
F4-80
Ly6C
Ly6G
NK1.1

S2

#### A Protein maxID correlation to protein/RNA:

#### Protein_secondID correlation to protein/RNA:

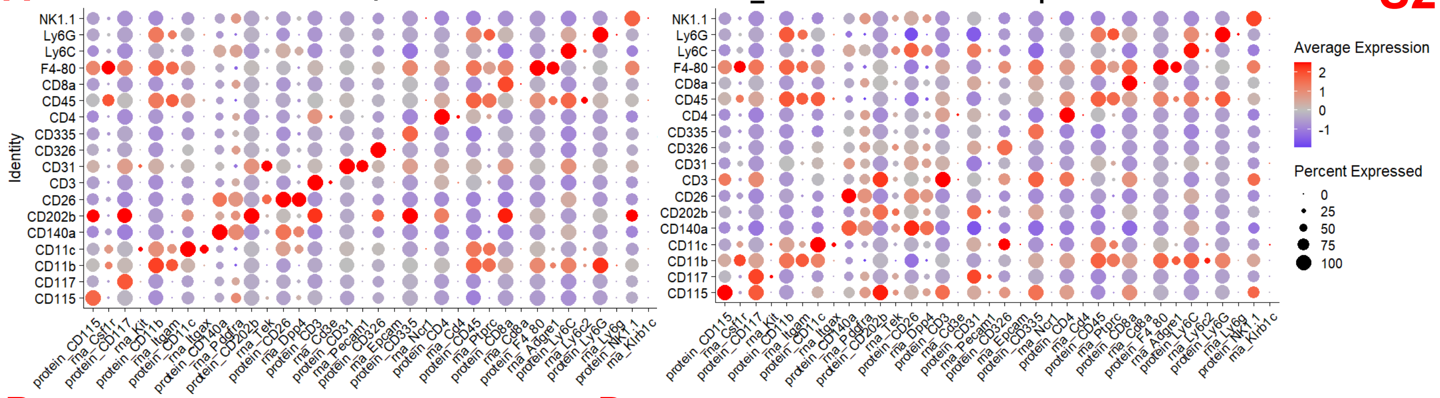


#### B Seurat clusters' protein/RNA expression:

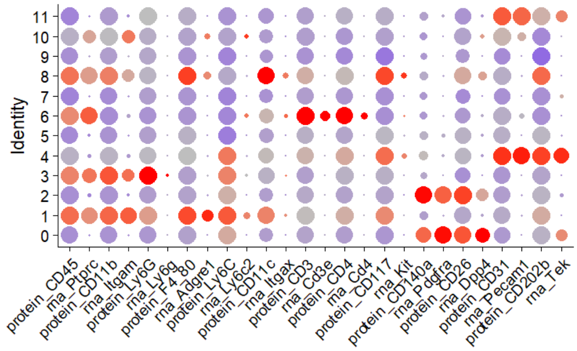


#### D Cluster identification workflow:

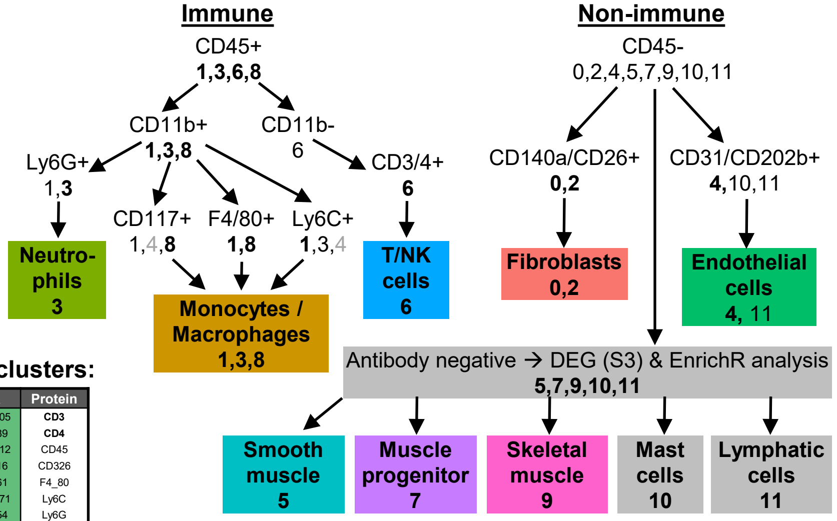


#### C Differentially-expressed protein (DEP) for Seurat clusters:

| Seurat clusters | | | |
|---|---|---|---|
| 0 | 1 | 2 | 3 |
| 4 | 5 | 6 | 7 |
| 8 | 9 | 10 | 11 |

| C# | $Log_2FC$ | % C pos | FDR | Protein |
|---|---|---|---|---|
| 0 | 1.24 | 0.98 | 0.00E+00 | **CD26** |
| | 0.98 | 0.88 | 0.00E+00 | **CD140a** |
| | -0.85 | 1.00 | 0.00E+00 | F4_80 |
| | -0.89 | 0.77 | 0.00E+00 | CD11c |
| | -1.01 | 0.92 | 0.00E+00 | CD31 |
| | -1.31 | 0.94 | 0.00E+00 | Ly6G |
| | -2.40 | 0.98 | 0.00E+00 | CD45 |
| | -2.48 | 0.97 | 0.00E+00 | CD11b |
| 1 | 1.71 | 1.00 | 0.00E+00 | **CD45** |
| | 1.57 | 1.00 | 0.00E+00 | **CD11b** |
| | 1.48 | 1.00 | 0.00E+00 | **F4_80** |
| | 1.18 | 0.95 | 0.00E+00 | **CD11c** |
| | -1.57 | 0.94 | 0.00E+00 | CD26 |
| | -1.97 | 0.44 | 0.00E+00 | CD140a |
| 2 | 1.47 | 0.94 | 0.00E+00 | **CD140a** |
| | 1.10 | 0.99 | 0.00E+00 | **CD26** |
| | -0.66 | 1.00 | 0.00E+00 | F4_80 |
| | -0.75 | 0.78 | 0.00E+00 | CD11c |
| | -0.81 | 0.94 | 3.56E-216 | CD31 |
| | -1.02 | 0.96 | 0.00E+00 | Ly6G |
| | -2.12 | 0.99 | 0.00E+00 | CD45 |
| | -2.13 | 0.99 | 0.00E+00 | CD11b |
| 3 | 2.09 | 1.00 | 0.00E+00 | **Ly6G** |
| | 1.63 | 1.00 | 0.00E+00 | **CD11b** |
| | 1.20 | 1.00 | 0.00E+00 | **CD45** |
| | -0.55 | 0.96 | 1.03E-06 | CD31 |
| | -2.26 | 0.36 | 0.00E+00 | CD140a |
| | -2.48 | 0.86 | 0.00E+00 | CD26 |
| 4 | 3.07 | 1.00 | 0.00E+00 | **CD31** |
| | 0.52 | 1.00 | 1.13E-201 | CD202b |
| | -1.05 | 0.50 | 2.34E-100 | CD140a |
| | -1.07 | 0.99 | 1.66E-27 | CD45 |
| | -1.19 | 0.99 | 8.62E-30 | CD11b |
| | -1.26 | 0.92 | 3.44E-141 | CD26 |
| 5 | -0.82 | 0.97 | 7.21E-47 | Ly6G |
| | -1.06 | 1.00 | 0.00E+00 | Ly6C |
| | -1.31 | 0.46 | 3.78E-127 | CD140a |
| | -1.51 | 0.89 | 3.09E-229 | CD26 |
| | -2.17 | 0.98 | 3.67E-225 | CD11b |
| | -2.17 | 0.98 | 2.08E-243 | CD45 |
| 6 | 1.34 | 0.99 | 9.18E-205 | **CD3** |
| | 1.09 | 0.98 | 7.52E-89 | **CD4** |
| | 0.89 | 1.00 | 2.64E-112 | CD45 |
| | 0.50 | 0.81 | 4.71E-16 | CD326 |
| | -0.75 | 1.00 | 1.04E-61 | F4_80 |
| | -0.89 | 1.00 | 2.34E-171 | Ly6C |
| | -1.03 | 0.95 | 9.09E-54 | Ly6G |
| | -1.92 | 0.87 | 5.06E-133 | CD26 |
| | -1.96 | 0.99 | 7.83E-50 | CD11b |
| | -2.00 | 0.38 | 6.71E-105 | CD140a |
| 7 | -0.69 | 1.00 | 7.78E-43 | F4_80 |
| | -0.78 | 0.78 | 1.08E-34 | CD11c |
| | -0.80 | 0.97 | 1.29E-19 | Ly6G |
| | -0.88 | 1.00 | 9.11E-143 | Ly6C |
| | -1.18 | 0.89 | 9.87E-74 | CD31 |
| | -1.97 | 0.39 | 4.90E-92 | CD140a |
| | -2.11 | 0.97 | 5.28E-54 | CD11b |
| | -2.22 | 0.99 | 1.18E-83 | CD45 |
| | -2.60 | 0.80 | 2.62E-210 | CD26 |
| 8 | 1.71 | 0.97 | 3.77E-151 | **CD11c** |
| | 1.08 | 1.00 | 1.30E-105 | **CD45** |
| | 0.82 | 1.00 | 2.09E-39 | F4_80 |
| | 0.64 | 1.00 | 1.44E-44 | CD11b |
| | -0.57 | 1.00 | 4.43E-40 | Ly6C |
| | -1.58 | 0.47 | 6.26E-38 | CD140a |
| 9 | -0.50 | 0.95 | 4.87E-30 | CD4 |
| | -0.77 | 1.00 | 6.51E-36 | F4_80 |
| | -0.82 | 0.94 | 1.80E-18 | Ly6G |
| | -0.90 | 0.74 | 2.81E-27 | CD11c |
| | -0.95 | 1.00 | 4.80E-88 | Ly6C |
| | -1.13 | 0.89 | 1.13E-31 | CD31 |
| | -1.98 | 0.96 | 5.02E-41 | CD11b |
| | -2.06 | 0.35 | 2.71E-53 | CD140a |
| | -2.10 | 0.97 | 4.87E-51 | CD45 |
| | -2.41 | 0.77 | 4.95E-106 | CD26 |
| 10 | 1.32 | 1.00 | 3.96E-42 | **CD31** |
| | -0.57 | 1.00 | 1.92E-08 | Ly6C |
| | -1.39 | 0.94 | 3.28E-06 | CD26 |
| | -1.90 | 0.41 | 9.83E-10 | CD140a |
| 11 | 2.33 | 1.00 | 4.72E-41 | **CD31** |
| | -0.72 | 1.00 | 8.54E-11 | Ly6C |
| | -1.44 | 0.47 | 1.47E-05 | CD140a |
| | -1.74 | 0.99 | 5.26E-03 | CD11b |
| | -1.99 | 0.99 | 7.23E-05 | CD45 |
| | -2.02 | 0.92 | 2.30E-13 | CD26 |

#### E DEP analysis for major cells:

| Major cell | $Log_2FC$ | FDR | Protein |
|---|---|---|---|
| Fibroblast | 2.52 | 0.00E+00 | **CD140a** |
| | 2.37 | 0.00E+00 | **CD26** |
| | -1.02 | 0.00E+00 | F4_80 |
| | -1.10 | 0.00E+00 | CD11c |
| | -1.21 | 0.00E+00 | CD31 |
| | -1.51 | 0.00E+00 | Ly6G |
| | -2.73 | 0.00E+00 | CD45 |
| | -2.79 | 0.00E+00 | CD11b |
| Macrophage | 1.73 | 0.00E+00 | **CD45** |
| | 1.56 | 0.00E+00 | **CD11b** |
| | 1.50 | 0.00E+00 | **F4_80** |
| | 1.28 | 0.00E+00 | **CD11c** |
| | -1.51 | 0.00E+00 | CD26 |
| | -1.97 | 0.00E+00 | CD140a |
| Neutrophil | 2.10 | 0.00E+00 | **Ly6G** |
| | 1.64 | 0.00E+00 | **CD11b** |
| | 1.21 | 0.00E+00 | **CD45** |
| | -0.54 | 7.40E-06 | CD31 |
| | -2.26 | 0.00E+00 | CD140a |
| | -2.48 | 0.00E+00 | CD26 |
| Endothelial cell | 3.07 | 0.00E+00 | **CD31** |
| | 0.50 | 3.21E-186 | CD202b |
| | -1.09 | 4.48E-108 | CD140a |
| | -1.10 | 6.53E-32 | CD45 |
| | -1.21 | 5.13E-33 | CD11b |
| | -1.30 | 2.04E-156 | CD26 |
| T cell / NK cell | 1.34 | 9.10E-205 | **CD3** |
| | 1.09 | 7.54E-89 | **CD4** |
| | 0.90 | 1.88E-112 | CD45 |
| | -0.74 | 1.61E-61 | F4_80 |
| | -0.88 | 1.19E-169 | Ly6C |
| | -1.02 | 2.75E-53 | Ly6G |
| | -1.93 | 1.84E-133 | CD26 |
| | -1.94 | 1.95E-49 | CD11b |
| | -2.00 | 3.28E-105 | CD140a |

#### G Count/feature violin plots for major cells:

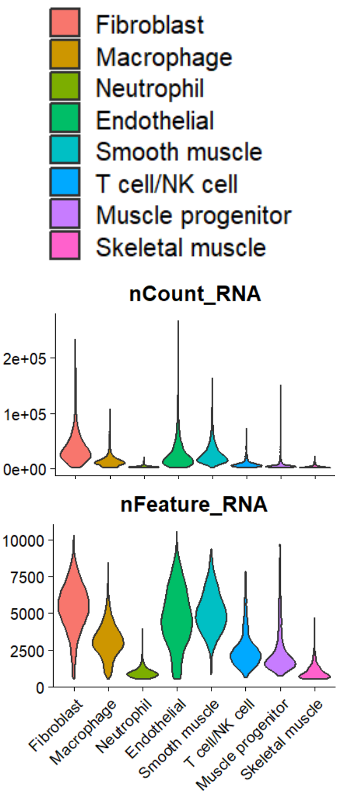


#### F Major cells' protein/RNA expression:

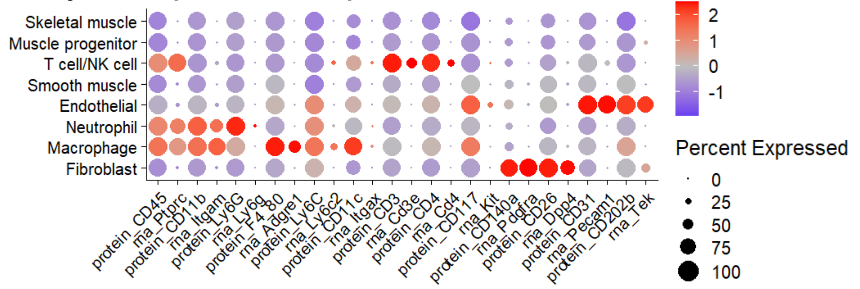

### Differentially-expressed gene analysis of Seurat clusters:

% cluster (C) pos ≥ 0.75
FDR < 0.01

**S3**

#### Cluster 0

| $Log_2FC$ | % C pos | % NC pos | FDR | Gene |
|---|---|---|---|---|
| 5.58 | 0.89 | 0.08 | 0.00E+00 | Scara5 |
| 5.31 | 0.79 | 0.13 | 0.00E+00 | Pcolce2 |
| 4.90 | 0.96 | 0.23 | 0.00E+00 | Clec3b |
| 4.65 | 0.85 | 0.14 | 0.00E+00 | Entpd2 |
| 4.51 | 0.75 | 0.12 | 0.00E+00 | Pcsk6 |
| 4.51 | 0.75 | 0.09 | 0.00E+00 | Pamr1 |
| 4.47 | 0.86 | 0.21 | 0.00E+00 | Sema3c |
| 4.41 | 0.99 | 0.79 | 0.00E+00 | Gsn |
| 4.31 | 0.98 | 0.34 | 0.00E+00 | Tnxb |
| 4.31 | 0.91 | 0.24 | 0.00E+00 | Gpc3 |
| 4.04 | 0.92 | 0.23 | 0.00E+00 | Islr |
| 4.02 | 0.92 | 0.21 | 0.00E+00 | Lama2 |
| 3.94 | 0.76 | 0.13 | 0.00E+00 | Tmeff2 |
| 3.90 | 0.88 | 0.26 | 0.00E+00 | Apod |
| 3.89 | 0.87 | 0.19 | 0.00E+00 | Lrrn4cl |
| 3.85 | 0.95 | 0.33 | 0.00E+00 | Col14a1 |
| 3.85 | 0.90 | 0.43 | 0.00E+00 | Pi16 |
| 3.84 | 0.79 | 0.17 | 0.00E+00 | Nova1 |
| 3.82 | 0.93 | 0.25 | 0.00E+00 | Ogn |
| 3.78 | 0.88 | 0.24 | 0.00E+00 | Dpep1 |

#### Cluster 1

| $Log_2FC$ | % C pos | % NC pos | FDR | Gene |
|---|---|---|---|---|
| 5.62 | 0.88 | 0.09 | 0.00E+00 | Ms4a6c |
| 5.45 | 0.81 | 0.08 | 0.00E+00 | Ms4a6d |
| 5.42 | 0.80 | 0.11 | 0.00E+00 | F13a1 |
| 5.40 | 0.99 | 0.24 | 0.00E+00 | Ctss |
| 5.37 | 0.77 | 0.06 | 0.00E+00 | Clec4a1 |
| 5.24 | 0.85 | 0.07 | 0.00E+00 | Cybb |
| 5.00 | 0.97 | 0.52 | 0.00E+00 | Lyz2 |
| 4.78 | 0.85 | 0.13 | 0.00E+00 | Fcgr2b |
| 4.74 | 0.91 | 0.19 | 0.00E+00 | Ccl9 |
| 4.60 | 0.77 | 0.06 | 0.00E+00 | Ccr5 |
| 4.30 | 0.90 | 0.67 | 0.00E+00 | Apoe |
| 4.12 | 0.90 | 0.26 | 0.00E+00 | Ctsc |
| 3.94 | 0.87 | 0.26 | 0.00E+00 | Csf1r |
| 3.72 | 0.83 | 0.43 | 0.00E+00 | Mafb |
| 3.61 | 0.91 | 0.16 | 0.00E+00 | Mpeg1 |
| 3.57 | 0.87 | 0.33 | 0.00E+00 | Pid1 |
| 3.57 | 0.85 | 0.15 | 0.00E+00 | Cd68 |
| 3.48 | 0.84 | 0.16 | 0.00E+00 | Cfp |
| 3.41 | 0.90 | 0.71 | 0.00E+00 | Lgmn |
| 3.29 | 0.87 | 0.52 | 0.00E+00 | Ifi30 |

#### Cluster 2

| $Log_2FC$ | % C pos | % NC pos | FDR | Gene |
|---|---|---|---|---|
| 4.27 | 0.89 | 0.21 | 0.00E+00 | Tnc |
| 4.06 | 0.86 | 0.22 | 0.00E+00 | Serpine1 |
| 4.02 | 0.88 | 0.15 | 0.00E+00 | Lrrc15 |
| 3.96 | 1.00 | 0.57 | 0.00E+00 | Timp1 |
| 3.90 | 0.83 | 0.10 | 0.00E+00 | Ccn4 |
| 3.85 | 0.81 | 0.18 | 0.00E+00 | Cxcl5 |
| 3.83 | 0.98 | 0.31 | 0.00E+00 | Col12a1 |
| 3.82 | 0.89 | 0.15 | 0.00E+00 | Ddah1 |
| 3.30 | 0.97 | 0.29 | 0.00E+00 | Plod2 |
| 3.12 | 0.82 | 0.23 | 0.00E+00 | Il33 |
| 3.10 | 0.88 | 0.15 | 0.00E+00 | Bcat1 |
| 3.03 | 0.94 | 0.71 | 0.00E+00 | Saa3 |
| 3.00 | 0.80 | 0.21 | 0.00E+00 | Ndufa4l2 |
| 2.99 | 0.76 | 0.17 | 0.00E+00 | Chl1 |
| 2.93 | 0.94 | 0.20 | 0.00E+00 | Nxn |
| 2.80 | 0.82 | 0.15 | 0.00E+00 | Col27a1 |
| 2.70 | 0.95 | 0.35 | 0.00E+00 | Pcsk5 |
| 2.70 | 0.99 | 0.67 | 0.00E+00 | Mt2 |
| 2.66 | 0.97 | 0.33 | 0.00E+00 | Fkbp11 |
| 2.65 | 0.95 | 0.27 | 0.00E+00 | Tmem45a |

#### Cluster 3

| $Log_2FC$ | % C pos | % NC pos | FDR | Gene |
|---|---|---|---|---|
| 6.49 | 0.80 | 0.08 | 0.00E+00 | Hcar2 |
| 6.48 | 0.99 | 0.40 | 0.00E+00 | S100a9 |
| 6.40 | 0.77 | 0.05 | 0.00E+00 | Cxcr2 |
| 6.22 | 0.98 | 0.34 | 0.00E+00 | S100a8 |
| 6.06 | 0.87 | 0.12 | 0.00E+00 | Acod1 |
| 5.91 | 0.92 | 0.13 | 0.00E+00 | Hdc |
| 5.59 | 0.81 | 0.11 | 0.00E+00 | Trem1 |
| 5.42 | 0.86 | 0.18 | 0.00E+00 | Csf3r |
| 5.27 | 0.89 | 0.16 | 0.00E+00 | Slc7a11 |
| 5.15 | 0.90 | 0.21 | 0.00E+00 | Samsn1 |
| 5.08 | 0.76 | 0.29 | 0.00E+00 | Ccl3 |
| 5.04 | 0.99 | 0.67 | 0.00E+00 | Cxcl2 |
| 4.87 | 0.99 | 0.38 | 0.00E+00 | Il1b |
| 4.87 | 0.81 | 0.20 | 0.00E+00 | Nlrp3 |
| 4.81 | 0.88 | 0.40 | 0.00E+00 | Il1r2 |
| 4.59 | 0.89 | 0.29 | 0.00E+00 | Slpi |
| 4.53 | 0.88 | 0.41 | 0.00E+00 | Trib1 |
| 4.53 | 0.79 | 0.24 | 0.00E+00 | Il1rn |
| 4.52 | 0.78 | 0.17 | 0.00E+00 | Sorl1 |
| 4.51 | 0.91 | 0.28 | 0.00E+00 | Clec4e |

#### Cluster 4

| $Log_2FC$ | % C pos | % NC pos | FDR | Gene |
|---|---|---|---|---|
| 8.93 | 0.97 | 0.02 | 0.00E+00 | Ptprb |
| 8.49 | 0.84 | 0.01 | 0.00E+00 | Tmem252 |
| 8.40 | 0.84 | 0.01 | 0.00E+00 | Grrp1 |
| 8.37 | 0.96 | 0.02 | 0.00E+00 | Mmrn2 |
| 8.36 | 0.81 | 0.01 | 0.00E+00 | Robo4 |
| 8.33 | 0.90 | 0.01 | 0.00E+00 | Adgrl4 |
| 8.27 | 0.87 | 0.01 | 0.00E+00 | Podxl |
| 8.26 | 0.90 | 0.01 | 0.00E+00 | Myct1 |
| 8.25 | 0.90 | 0.01 | 0.00E+00 | Cyyr1 |
| 8.19 | 0.77 | 0.03 | 0.00E+00 | Vwf |
| 8.02 | 0.97 | 0.03 | 0.00E+00 | Cdh5 |
| 7.93 | 0.88 | 0.01 | 0.00E+00 | Sox18 |
| 7.86 | 0.91 | 0.02 | 0.00E+00 | Emcn |
| 7.65 | 0.90 | 0.09 | 0.00E+00 | Plvap |
| 7.54 | 0.85 | 0.02 | 0.00E+00 | C130074G19Rik |
| 7.43 | 0.92 | 0.02 | 0.00E+00 | Tie1 |
| 7.40 | 0.78 | 0.01 | 0.00E+00 | Erg |
| 7.40 | 0.76 | 0.01 | 0.00E+00 | She |
| 7.39 | 0.79 | 0.02 | 0.00E+00 | Arhgef15 |
| 7.34 | 0.98 | 0.06 | 0.00E+00 | Pecam1 |

#### Cluster 5

| $Log_2FC$ | % C pos | % NC pos | FDR | Gene |
|---|---|---|---|---|
| 9.09 | 0.93 | 0.07 | 0.00E+00 | Rgs5 |
| 8.80 | 0.92 | 0.02 | 0.00E+00 | Myh11 |
| 8.69 | 0.87 | 0.01 | 0.00E+00 | Higd1b |
| 8.47 | 0.88 | 0.01 | 0.00E+00 | Ano1 |
| 8.14 | 0.77 | 0.01 | 0.00E+00 | Lmod1 |
| 7.81 | 0.82 | 0.01 | 0.00E+00 | Trarg1 |
| 7.69 | 0.83 | 0.01 | 0.00E+00 | Mrvi1 |
| 7.34 | 0.99 | 0.03 | 0.00E+00 | Gucy1a1 |
| 7.17 | 0.97 | 0.04 | 0.00E+00 | Gm13889 |
| 6.95 | 0.89 | 0.02 | 0.00E+00 | Atp1b2 |
| 6.91 | 0.84 | 0.02 | 0.00E+00 | Cox4i2 |
| 6.90 | 0.95 | 0.03 | 0.00E+00 | Gucy1b1 |
| 6.79 | 0.75 | 0.03 | 0.00E+00 | Tbx2 |
| 6.73 | 0.78 | 0.03 | 0.00E+00 | Parm1 |
| 6.56 | 0.87 | 0.04 | 0.00E+00 | Rgs4 |
| 6.48 | 1.00 | 0.06 | 0.00E+00 | Notch3 |
| 6.47 | 0.87 | 0.03 | 0.00E+00 | Daam2 |
| 6.42 | 1.00 | 0.16 | 0.00E+00 | Mylk |
| 6.35 | 0.91 | 0.05 | 0.00E+00 | Ppp1r14a |
| 6.30 | 0.98 | 0.16 | 0.00E+00 | Myl9 |

#### Cluster 6

| $Log_2FC$ | % C pos | % NC pos | FDR | Gene |
|---|---|---|---|---|
| 9.81 | 0.75 | 0.01 | 0.00E+00 | Gimap3 |
| 8.50 | 0.87 | 0.01 | 0.00E+00 | Ptprcap |
| 5.68 | 0.82 | 0.13 | 0.00E+00 | Sept1 |
| 5.23 | 0.83 | 0.19 | 0.00E+00 | AW112010 |
| 4.46 | 0.83 | 0.09 | 0.00E+00 | Ltb |
| 4.35 | 0.95 | 0.28 | 0.00E+00 | Ets1 |
| 3.88 | 0.76 | 0.23 | 0.00E+00 | H2-Q7 |
| 2.93 | 0.79 | 0.21 | 0.00E+00 | Hcst |
| 2.79 | 0.85 | 0.26 | 0.00E+00 | Dock2 |
| 2.52 | 0.91 | 0.31 | 0.00E+00 | Rac2 |
| 2.37 | 0.83 | 0.45 | 3.74E-265 | Id2 |
| 2.33 | 0.83 | 0.47 | 1.92E-255 | Ccnd2 |
| 2.32 | 0.97 | 0.43 | 0.00E+00 | Ptprc |
| 2.28 | 0.90 | 0.33 | 0.00E+00 | Ptpn18 |
| 2.24 | 0.78 | 0.28 | 0.00E+00 | Arhgap45 |
| 2.09 | 0.83 | 0.67 | 1.10E-142 | Hmgb2 |
| 2.04 | 0.97 | 0.43 | 0.00E+00 | Cd52 |
| 2.04 | 0.98 | 0.80 | 0.00E+00 | Mbnl1 |
| 2.03 | 0.85 | 0.70 | 5.09E-187 | Ifngr1 |
| 2.03 | 0.78 | 0.57 | 1.60E-182 | Arhgef1 |

#### Cluster 7

| $Log_2FC$ | % C pos | % NC pos | FDR | Gene |
|---|---|---|---|---|
| 11.19 | 0.84 | 0.00 | 0.00E+00 | Pax7 |
| 10.95 | 0.77 | 0.01 | 0.00E+00 | Myf5 |
| 10.19 | 0.87 | 0.01 | 0.00E+00 | Chodl |
| 6.41 | 0.76 | 0.07 | 0.00E+00 | Crlf1 |
| 5.27 | 0.84 | 0.18 | 0.00E+00 | Spry1 |
| 4.68 | 0.88 | 0.26 | 0.00E+00 | Peg3 |
| 4.36 | 0.79 | 0.17 | 0.00E+00 | Cd200 |
| 4.17 | 0.84 | 0.18 | 0.00E+00 | Ncam1 |
| 3.41 | 0.91 | 0.60 | 0.00E+00 | Dag1 |
| 3.30 | 0.77 | 0.42 | 8.93E-271 | Pdlim4 |
| 2.91 | 0.81 | 0.22 | 0.00E+00 | Ltbp4 |
| 2.79 | 0.85 | 0.53 | 0.00E+00 | Cnn3 |
| 2.78 | 0.98 | 0.66 | 0.00E+00 | Zbtb20 |
| 2.72 | 0.93 | 0.60 | 0.00E+00 | Gpx3 |
| 2.67 | 0.81 | 0.40 | 4.44E-288 | Crip2 |
| 2.66 | 0.92 | 0.50 | 0.00E+00 | Gas1 |
| 2.43 | 0.80 | 0.53 | 6.75E-192 | Gstm1 |
| 2.42 | 0.93 | 0.55 | 0.00E+00 | Nfib |
| 2.41 | 0.81 | 0.57 | 9.79E-221 | Ttc3 |
| 2.26 | 0.84 | 0.52 | 5.96E-205 | Slc43a3 |

#### Cluster 8

| $Log_2FC$ | % C pos | % NC pos | FDR | Gene |
|---|---|---|---|---|
| 3.25 | 0.83 | 0.49 | 6.61E-94 | Naaa |
| 3.10 | 0.86 | 0.14 | 0.00E+00 | Shtn1 |
| 2.92 | 0.81 | 0.31 | 2.06E-174 | Cd74 |
| 2.91 | 1.00 | 0.91 | 1.76E-159 | Cst3 |
| 2.65 | 0.75 | 0.23 | 4.38E-208 | Pak1 |
| 2.63 | 0.86 | 0.34 | 1.52E-174 | Psmb9 |
| 2.62 | 0.77 | 0.26 | 2.41E-195 | Ezh2 |
| 2.55 | 0.80 | 0.24 | 1.26E-221 | Sms |
| 2.54 | 0.81 | 0.46 | 5.60E-100 | Ckb |
| 2.54 | 0.81 | 0.33 | 1.79E-170 | Dctpp1 |
| 2.48 | 0.95 | 0.58 | 2.53E-153 | Psmb8 |
| 2.23 | 0.77 | 0.24 | 8.36E-188 | Vrk1 |
| 2.21 | 0.81 | 0.36 | 5.24E-142 | Olfm1 |
| 2.13 | 0.91 | 0.39 | 3.84E-191 | Rgs10 |
| 2.11 | 0.88 | 0.46 | 8.68E-135 | Fnbp1 |
| 2.07 | 0.98 | 0.60 | 1.55E-195 | Ifi30 |
| 2.05 | 0.90 | 0.30 | 1.07E-198 | Dock10 |
| 2.05 | 0.84 | 0.22 | 5.56E-240 | Arhgef6 |
| 1.98 | 0.87 | 0.20 | 4.18E-277 | AI662270 |
| 1.96 | 0.91 | 0.50 | 1.12E-144 | Anp32e |

#### Cluster 9

| $Log_2FC$ | % C pos | % NC pos | FDR | Gene |
|---|---|---|---|---|
| 12.49 | 0.82 | 0.00 | 0.00E+00 | Myot |
| 12.47 | 1.00 | 0.03 | 0.00E+00 | Tnnc2 |
| 12.34 | 0.99 | 0.02 | 0.00E+00 | Myl1 |
| 12.30 | 0.83 | 0.00 | 0.00E+00 | Mybpc2 |
| 12.30 | 1.00 | 0.07 | 0.00E+00 | Acta1 |
| 12.21 | 1.00 | 0.02 | 0.00E+00 | Ckm |
| 12.18 | 0.80 | 0.00 | 0.00E+00 | Smpx |
| 12.16 | 0.79 | 0.00 | 0.00E+00 | Trdn |
| 12.15 | 0.99 | 0.07 | 0.00E+00 | Mylpf |
| 12.15 | 0.90 | 0.01 | 0.00E+00 | Pvalb |
| 12.08 | 0.90 | 0.00 | 0.00E+00 | Apobec2 |
| 11.98 | 0.99 | 0.01 | 0.00E+00 | Atp2a1 |
| 11.97 | 0.89 | 0.01 | 0.00E+00 | Mb |
| 11.96 | 0.90 | 0.01 | 0.00E+00 | Myh1 |
| 11.90 | 0.99 | 0.06 | 0.00E+00 | Tnnt3 |
| 11.79 | 0.91 | 0.01 | 0.00E+00 | Xirp2 |
| 11.78 | 0.95 | 0.01 | 0.00E+00 | Pgam2 |
| 11.54 | 0.93 | 0.01 | 0.00E+00 | Myoz1 |
| 11.53 | 0.98 | 0.01 | 0.00E+00 | Tcap |
| 11.40 | 0.87 | 0.00 | 0.00E+00 | Cox8b |

#### Cluster 10

| $Log_2FC$ | % C pos | % NC pos | FDR | Gene |
|---|---|---|---|---|
| 12.63 | 0.87 | 0.00 | 0.00E+00 | Ms4a2 |
| 12.25 | 0.93 | 0.00 | 0.00E+00 | Cd200r3 |
| 11.31 | 0.95 | 0.01 | 0.00E+00 | Cyp11a1 |
| 9.07 | 0.85 | 0.01 | 0.00E+00 | Sytl3 |
| 8.85 | 1.00 | 0.03 | 0.00E+00 | Gata2 |
| 8.03 | 0.89 | 0.01 | 0.00E+00 | Cdh1 |
| 6.10 | 0.92 | 0.17 | 1.00E-106 | Hgf |
| 5.77 | 0.77 | 0.16 | 4.35E-72 | Itga2b |
| 5.07 | 0.83 | 0.18 | 1.17E-75 | Tbc1d4 |
| 5.05 | 0.78 | 0.12 | 2.66E-96 | Tmem71 |
| 4.69 | 0.85 | 0.14 | 1.80E-97 | Tec |
| 4.42 | 0.81 | 0.15 | 2.83E-79 | Lat2 |
| 4.38 | 0.82 | 0.16 | 2.14E-75 | Csf2rb2 |
| 4.34 | 0.91 | 0.32 | 2.07E-58 | Syne1 |
| 3.92 | 0.78 | 0.09 | 8.91E-121 | Rab44 |
| 3.71 | 0.86 | 0.12 | 7.36E-113 | Ripor2 |
| 3.64 | 0.76 | 0.17 | 5.67E-58 | Arhgap15 |
| 3.55 | 0.87 | 0.14 | 1.09E-93 | Il18rap |
| 3.54 | 0.98 | 0.46 | 3.14E-50 | Ccl6 |
| 3.52 | 0.86 | 0.29 | 3.42E-49 | Ets1 |

#### Cluster 11

| $Log_2FC$ | % C pos | % NC pos | FDR | Gene |
|---|---|---|---|---|
| 12.80 | 1.00 | 0.00 | 0.00E+00 | Mmrn1 |
| 11.60 | 1.00 | 0.00 | 0.00E+00 | Reln |
| 11.41 | 0.92 | 0.01 | 0.00E+00 | Ccl21a |
| 10.06 | 0.86 | 0.00 | 0.00E+00 | Tbx1 |
| 10.04 | 0.77 | 0.00 | 0.00E+00 | Dtx1 |
| 9.53 | 0.99 | 0.02 | 0.00E+00 | Flt4 |
| 9.28 | 0.90 | 0.01 | 0.00E+00 | Gpm6a |
| 8.75 | 0.90 | 0.01 | 0.00E+00 | Tc2n |
| 8.69 | 0.93 | 0.02 | 0.00E+00 | Slc45a3 |
| 8.48 | 0.88 | 0.03 | 0.00E+00 | Lyve1 |
| 8.46 | 0.77 | 0.01 | 0.00E+00 | Sh3gl3 |
| 7.85 | 0.93 | 0.02 | 0.00E+00 | Klhl4 |
| 7.80 | 0.84 | 0.01 | 0.00E+00 | Wipf3 |
| 7.55 | 0.96 | 0.03 | 0.00E+00 | Prox1 |
| 7.06 | 1.00 | 0.14 | 7.67E-124 | Pard6g |
| 6.97 | 1.00 | 0.03 | 0.00E+00 | Cldn5 |
| 6.22 | 0.78 | 0.02 | 0.00E+00 | Gpr182 |
| 6.20 | 0.84 | 0.05 | 3.19E-231 | Clca3a1 |
| 5.95 | 0.82 | 0.02 | 0.00E+00 | Rassf9 |
| 5.88 | 0.90 | 0.05 | 3.89E-244 | Myzap |

### Differentially-expressed gene analysis of annotated major cells:

% cluster (C) pos ≥ 0.75 (* ≥ 0.5)
FDR < 0.01

**S4**

#### Fibroblasts

| $Log_2FC$ | % C pos | % NC pos | FDR | Gene |
|---|---|---|---|---|
| 5.67 | 0.82 | 0.11 | 0.00E+00 | Clec3b |
| 5.62 | 1.00 | 0.43 | 0.00E+00 | Dcn |
| 5.55 | 0.94 | 0.12 | 0.00E+00 | Col14a1 |
| 5.43 | 0.78 | 0.05 | 0.00E+00 | Sema3c |
| 5.32 | 0.87 | 0.06 | 0.00E+00 | Ogn |
| 5.31 | 0.84 | 0.08 | 0.00E+00 | Efemp1 |
| 5.27 | 0.97 | 0.06 | 0.00E+00 | Pdgfra |
| 5.22 | 0.85 | 0.04 | 0.00E+00 | Dpep1 |
| 5.21 | 0.75 | 0.03 | 0.00E+00 | Osr1 |
| 5.19 | 0.84 | 0.06 | 0.00E+00 | Gpc3 |
| 5.08 | 0.89 | 0.04 | 0.00E+00 | Dclk1 |
| 5.06 | 0.77 | 0.04 | 0.00E+00 | Lrrn4cl |
| 5.01 | 0.98 | 0.22 | 0.00E+00 | Mfap5 |
| 5.00 | 0.98 | 0.18 | 0.00E+00 | Dpt |
| 5.00 | 0.87 | 0.16 | 0.00E+00 | Sfrp2 |
| 4.92 | 0.96 | 0.19 | 0.00E+00 | Lum |
| 4.89 | 0.99 | 0.23 | 0.00E+00 | Aebp1 |
| 4.87 | 0.91 | 0.17 | 0.00E+00 | Tnxb |
| 4.85 | 0.80 | 0.04 | 0.00E+00 | Ptgis |
| 4.82 | 0.83 | 0.14 | 0.00E+00 | Prg4 |
| 4.81 | 0.96 | 0.09 | 0.00E+00 | Abi3bp |
| 4.81 | 0.82 | 0.09 | 0.00E+00 | Apod |
| 4.81 | 0.94 | 0.12 | 0.00E+00 | Fndc1 |
| 4.78 | 0.87 | 0.06 | 0.00E+00 | Fbln1 |
| 4.77 | 1.00 | 0.82 | 0.00E+00 | Col1a2 |
| 4.76 | 0.86 | 0.05 | 0.00E+00 | Lrrc17 |
| 4.76 | 0.99 | 0.19 | 0.00E+00 | Serpinf1 |
| 4.74 | 1.00 | 0.90 | 0.00E+00 | Col3a1 |
| 4.70 | 1.00 | 0.85 | 0.00E+00 | Col1a1 |
| 4.70 | 0.93 | 0.13 | 0.00E+00 | Cpxm1 |

#### Monocytes/Macrophages

| $Log_2FC$ | % C pos | % NC pos | FDR | Gene |
|---|---|---|---|---|
| 5.88 | 0.87 | 0.08 | 0.00E+00 | Ms4a6c |
| 5.65 | 0.78 | 0.10 | 0.00E+00 | F13a1 |
| 5.65 | 0.80 | 0.08 | 0.00E+00 | Ms4a6d |
| 5.63 | 0.99 | 0.23 | 0.00E+00 | Ctss |
| 5.47 | 0.75 | 0.06 | 0.00E+00 | Clec4a1 |
| 5.29 | 0.83 | 0.07 | 0.00E+00 | Cybb |
| 5.10 | 0.97 | 0.52 | 0.00E+00 | Lyz2 |
| 5.04 | 0.90 | 0.18 | 0.00E+00 | Ccl9 |
| 4.92 | 0.85 | 0.12 | 0.00E+00 | Fcgr2b |
| 4.63 | 0.75 | 0.06 | 0.00E+00 | Ccr5 |
| 4.33 | 0.90 | 0.67 | 0.00E+00 | Apoe |
| 4.21 | 0.89 | 0.25 | 0.00E+00 | Ctsc |
| 4.05 | 0.85 | 0.25 | 0.00E+00 | Csf1r |
| 3.72 | 0.85 | 0.14 | 0.00E+00 | Cd68 |
| 3.71 | 0.90 | 0.15 | 0.00E+00 | Mpeg1 |
| 3.69 | 0.82 | 0.43 | 0.00E+00 | Mafb |
| 3.62 | 0.87 | 0.33 | 0.00E+00 | Pid1 |
| 3.54 | 0.83 | 0.15 | 0.00E+00 | Cfp |
| 3.52 | 0.87 | 0.51 | 0.00E+00 | Ifi30 |
| 3.46 | 0.89 | 0.71 | 0.00E+00 | Lgmn |
| 3.36 | 0.80 | 0.54 | 0.00E+00 | Dab2 |
| 3.35 | 0.80 | 0.11 | 0.00E+00 | Fermt3 |
| 3.32 | 0.95 | 0.27 | 0.00E+00 | Ucp2 |
| 3.18 | 0.85 | 0.18 | 0.00E+00 | Ptpn18 |
| 3.14 | 0.88 | 0.46 | 0.00E+00 | Unc93b1 |
| 3.10 | 0.98 | 0.24 | 0.00E+00 | Laptm5 |
| 3.04 | 0.77 | 0.13 | 0.00E+00 | Nrros |
| 2.98 | 0.97 | 0.71 | 0.00E+00 | Lgals3 |
| 2.97 | 0.86 | 0.33 | 0.00E+00 | Ccl6 |
| 2.69 | 0.76 | 0.38 | 0.00E+00 | Pycard |

#### Neutrophils

| $Log_2FC$ | % C pos | % NC pos | FDR | Gene |
|---|---|---|---|---|
| 6.64 | 0.80 | 0.08 | 0.00E+00 | Hcar2 |
| 6.59 | 0.99 | 0.40 | 0.00E+00 | S100a9 |
| 6.53 | 0.77 | 0.05 | 0.00E+00 | Cxcr2 |
| 6.33 | 0.98 | 0.33 | 0.00E+00 | S100a8 |
| 6.16 | 0.87 | 0.12 | 0.00E+00 | Acod1 |
| 6.03 | 0.92 | 0.12 | 0.00E+00 | Hdc |
| 5.66 | 0.81 | 0.11 | 0.00E+00 | Trem1 |
| 5.49 | 0.86 | 0.17 | 0.00E+00 | Csf3r |
| 5.33 | 0.89 | 0.16 | 0.00E+00 | Slc7a11 |
| 5.19 | 0.90 | 0.21 | 0.00E+00 | Samsn1 |
| 5.11 | 0.76 | 0.29 | 0.00E+00 | Ccl3 |
| 5.09 | 0.99 | 0.67 | 0.00E+00 | Cxcl2 |
| 4.91 | 0.81 | 0.20 | 0.00E+00 | Nlrp3 |
| 4.91 | 0.99 | 0.38 | 0.00E+00 | Il1b |
| 4.85 | 0.88 | 0.40 | 0.00E+00 | Il1r2 |
| 4.61 | 0.89 | 0.29 | 0.00E+00 | Slpi |
| 4.57 | 0.88 | 0.41 | 0.00E+00 | Trib1 |
| 4.57 | 0.79 | 0.23 | 0.00E+00 | Il1rn |
| 4.56 | 0.79 | 0.17 | 0.00E+00 | Sorl1 |
| 4.53 | 0.91 | 0.28 | 0.00E+00 | Clec4e |
| 4.45 | 0.87 | 0.43 | 0.00E+00 | Lmnb1 |
| 4.42 | 1.00 | 0.63 | 0.00E+00 | Srgn |
| 4.41 | 0.91 | 0.34 | 0.00E+00 | Mxd1 |
| 4.38 | 0.97 | 0.30 | 0.00E+00 | Clec4d |
| 4.36 | 0.91 | 0.26 | 0.00E+00 | Ccr1 |
| 4.34 | 0.96 | 0.39 | 0.00E+00 | Plek |
| 4.30 | 0.77 | 0.19 | 0.00E+00 | Cd300lf |
| 4.23 | 0.92 | 0.43 | 0.00E+00 | Cd14 |
| 4.18 | 0.96 | 0.39 | 0.00E+00 | Lilr4b |
| 4.12 | 0.84 | 0.39 | 0.00E+00 | Csrnp1 |

#### Endothelial cells

| $Log_2FC$ | % C pos | % NC pos | FDR | Gene |
|---|---|---|---|---|
| 9.03 | 0.96 | 0.02 | 0.00E+00 | Ptprb |
| 8.94 | 0.87 | 0.01 | 0.00E+00 | Podxl |
| 8.81 | 0.90 | 0.01 | 0.00E+00 | Myct1 |
| 8.67 | 0.95 | 0.02 | 0.00E+00 | Mmrn2 |
| 8.63 | 0.88 | 0.01 | 0.00E+00 | Sox18 |
| 8.61 | 0.80 | 0.01 | 0.00E+00 | Robo4 |
| 8.60 | 0.89 | 0.01 | 0.00E+00 | Cyyr1 |
| 8.57 | 0.83 | 0.01 | 0.00E+00 | Grrp1 |
| 8.51 | 0.81 | 0.01 | 0.00E+00 | Tmem252 |
| 8.42 | 0.89 | 0.01 | 0.00E+00 | Adgrl4 |
| 8.29 | 0.97 | 0.03 | 0.00E+00 | Cdh5 |
| 8.24 | 0.91 | 0.02 | 0.00E+00 | Emcn |
| 8.14 | 0.92 | 0.01 | 0.00E+00 | Tie1 |
| 7.82 | 0.76 | 0.01 | 0.00E+00 | She |
| 7.76 | 0.89 | 0.09 | 0.00E+00 | Plvap |
| 7.68 | 0.77 | 0.01 | 0.00E+00 | Erg |
| 7.54 | 0.84 | 0.02 | 0.00E+00 | C130074G19Rik |
| 7.51 | 0.78 | 0.02 | 0.00E+00 | Arhgef15 |
| 7.49 | 0.98 | 0.07 | 0.00E+00 | Egfl7 |
| 7.48 | 0.98 | 0.06 | 0.00E+00 | Pecam1 |
| 7.21 | 0.81 | 0.02 | 0.00E+00 | Ushbp1 |
| 6.78 | 0.83 | 0.05 | 0.00E+00 | Mecom |
| 6.67 | 0.82 | 0.02 | 0.00E+00 | Rapgef5 |
| 6.61 | 0.80 | 0.14 | 0.00E+00 | Aqp1 |
| 6.58 | 0.75 | 0.02 | 0.00E+00 | Spns2 |
| 6.44 | 0.81 | 0.02 | 0.00E+00 | Icam2 |
| 6.33 | 0.88 | 0.04 | 0.00E+00 | Tspan7 |
| 6.24 | 0.83 | 0.02 | 0.00E+00 | Adgrg1 |
| 6.19 | 0.85 | 0.07 | 0.00E+00 | Kdr |
| 6.07 | 0.92 | 0.07 | 0.00E+00 | Flt1 |

#### Smooth muscle cells

| $Log_2FC$ | % C pos | % NC pos | FDR | Gene |
|---|---|---|---|---|
| 9.06 | 0.93 | 0.07 | 0.00E+00 | Rgs5 |
| 8.80 | 0.92 | 0.02 | 0.00E+00 | Myh11 |
| 8.72 | 0.87 | 0.01 | 0.00E+00 | Higd1b |
| 8.46 | 0.89 | 0.01 | 0.00E+00 | Ano1 |
| 8.09 | 0.76 | 0.01 | 0.00E+00 | Lmod1 |
| 7.80 | 0.82 | 0.01 | 0.00E+00 | Trarg1 |
| 7.67 | 0.83 | 0.01 | 0.00E+00 | Mrvi1 |
| 7.34 | 0.99 | 0.03 | 0.00E+00 | Gucy1a1 |
| 7.17 | 0.97 | 0.04 | 0.00E+00 | Gm13889 |
| 6.94 | 0.89 | 0.02 | 0.00E+00 | Atp1b2 |
| 6.91 | 0.95 | 0.03 | 0.00E+00 | Gucy1b1 |
| 6.90 | 0.84 | 0.03 | 0.00E+00 | Cox4i2 |
| 6.79 | 0.75 | 0.03 | 0.00E+00 | Tbx2 |
| 6.72 | 0.78 | 0.03 | 0.00E+00 | Parm1 |
| 6.56 | 0.87 | 0.04 | 0.00E+00 | Rgs4 |
| 6.47 | 1.00 | 0.06 | 0.00E+00 | Notch3 |
| 6.47 | 0.87 | 0.03 | 0.00E+00 | Daam2 |
| 6.42 | 1.00 | 0.16 | 0.00E+00 | Mylk |
| 6.36 | 0.91 | 0.05 | 0.00E+00 | Ppp1r14a |
| 6.30 | 0.98 | 0.16 | 0.00E+00 | Myl9 |
| 5.96 | 0.78 | 0.05 | 0.00E+00 | Carmn |
| 5.95 | 0.92 | 0.15 | 0.00E+00 | Tagln |
| 5.72 | 0.96 | 0.29 | 0.00E+00 | Acta2 |
| 5.61 | 0.87 | 0.03 | 0.00E+00 | Synpo2 |
| 5.55 | 0.95 | 0.09 | 0.00E+00 | Ednra |
| 5.53 | 0.85 | 0.02 | 0.00E+00 | Gja4 |
| 5.50 | 0.79 | 0.13 | 0.00E+00 | Abcc9 |
| 5.31 | 0.96 | 0.06 | 0.00E+00 | Tinagl1 |
| 5.27 | 0.82 | 0.04 | 0.00E+00 | Cobll1 |
| 5.20 | 0.86 | 0.08 | 0.00E+00 | Rasl12 |

#### T cells/NK cells*

| $Log_2FC$ | % C pos | % NC pos | FDR | Gene |
|---|---|---|---|---|
| 10.40 | 0.55 | 0.00 | 0.00E+00 | Cd3e |
| 10.03 | 0.59 | 0.00 | 0.00E+00 | Ikzf3 |
| 9.93 | 0.61 | 0.00 | 0.00E+00 | Cd3g |
| 9.81 | 0.75 | 0.01 | 0.00E+00 | Gimap3 |
| 9.71 | 0.75 | 0.01 | 0.00E+00 | Trbc2 |
| 9.66 | 0.75 | 0.01 | 0.00E+00 | Il2rb |
| 9.63 | 0.61 | 0.01 | 0.00E+00 | Trbc1 |
| 9.52 | 0.53 | 0.00 | 0.00E+00 | Sh2d2a |
| 9.13 | 0.58 | 0.01 | 0.00E+00 | Gata3 |
| 9.12 | 0.53 | 0.01 | 0.00E+00 | Icos |
| 8.92 | 0.58 | 0.01 | 0.00E+00 | Cd3d |
| 8.49 | 0.87 | 0.01 | 0.00E+00 | Ptprcap |
| 8.44 | 0.72 | 0.01 | 0.00E+00 | Skap1 |
| 8.24 | 0.53 | 0.02 | 0.00E+00 | Trac |
| 8.10 | 0.61 | 0.01 | 0.00E+00 | Lck |
| 7.88 | 0.55 | 0.02 | 0.00E+00 | Bcl11b |
| 7.71 | 0.68 | 0.02 | 0.00E+00 | Cd2 |
| 7.31 | 0.64 | 0.02 | 0.00E+00 | Tnfrsf18 |
| 7.18 | 0.58 | 0.03 | 0.00E+00 | Ms4a4b |
| 7.03 | 0.56 | 0.08 | 0.00E+00 | Ikzf2 |
| 6.70 | 0.51 | 0.04 | 0.00E+00 | Il2ra |
| 6.05 | 0.60 | 0.03 | 0.00E+00 | Il18r1 |
| 5.84 | 0.63 | 0.04 | 0.00E+00 | Gimap4 |
| 5.73 | 0.54 | 0.03 | 0.00E+00 | Gimap8 |
| 5.67 | 0.82 | 0.13 | 0.00E+00 | Sept1 |
| 5.44 | 0.72 | 0.04 | 0.00E+00 | Spn |
| 5.43 | 0.71 | 0.06 | 0.00E+00 | Il7r |
| 5.42 | 0.73 | 0.04 | 0.00E+00 | Gimap1 |
| 5.23 | 0.58 | 0.04 | 0.00E+00 | Gimap5 |
| 5.23 | 0.83 | 0.19 | 0.00E+00 | AW112010 |

#### Muscle progenitor cells*

| $Log_2FC$ | % C pos | % NC pos | FDR | Gene |
|---|---|---|---|---|
| 11.41 | 0.88 | 0.00 | 0.00E+00 | Pax7 |
| 10.95 | 0.81 | 0.00 | 0.00E+00 | Myf5 |
| 10.90 | 0.55 | 0.00 | 0.00E+00 | Chrdl2 |
| 10.47 | 0.79 | 0.01 | 0.00E+00 | Fgfr4 |
| 10.21 | 0.89 | 0.01 | 0.00E+00 | Chodl |
| 9.51 | 0.68 | 0.01 | 0.00E+00 | Cdh15 |
| 8.88 | 0.51 | 0.01 | 0.00E+00 | Pde1c |
| 6.77 | 0.54 | 0.02 | 0.00E+00 | Map2 |
| 6.69 | 0.56 | 0.02 | 0.00E+00 | Prox1 |
| 6.64 | 0.67 | 0.08 | 0.00E+00 | Clmn |
| 6.42 | 0.78 | 0.07 | 0.00E+00 | Crlf1 |
| 6.08 | 0.65 | 0.04 | 0.00E+00 | Heyl |
| 5.30 | 0.87 | 0.18 | 0.00E+00 | Spry1 |
| 4.74 | 0.92 | 0.26 | 0.00E+00 | Peg3 |
| 4.59 | 0.54 | 0.12 | 0.00E+00 | Ncald |
| 4.35 | 0.81 | 0.17 | 0.00E+00 | Cd200 |
| 4.31 | 0.69 | 0.18 | 0.00E+00 | Dmd |
| 4.29 | 0.64 | 0.14 | 0.00E+00 | Sema6a |
| 4.28 | 0.59 | 0.10 | 0.00E+00 | Bmp4 |
| 4.27 | 0.64 | 0.18 | 0.00E+00 | Ank3 |
| 4.16 | 0.62 | 0.24 | 2.05E-222 | Pde10a |
| 4.15 | 0.72 | 0.21 | 0.00E+00 | F2r |
| 4.10 | 0.85 | 0.18 | 0.00E+00 | Ncam1 |
| 4.05 | 0.72 | 0.25 | 0.00E+00 | Dtx4 |
| 3.98 | 0.57 | 0.13 | 0.00E+00 | Abca8b |
| 3.93 | 0.52 | 0.19 | 2.42E-176 | Nedd4l |
| 3.88 | 0.51 | 0.05 | 0.00E+00 | Atp1a2 |
| 3.81 | 0.75 | 0.13 | 0.00E+00 | Fxyd6 |
| 3.76 | 0.63 | 0.24 | 1.26E-250 | Sytl2 |
| 3.75 | 0.70 | 0.31 | 3.93E-249 | Tanc2 |

#### Skeletal muscle cells

| $Log_2FC$ | % C pos | % NC pos | FDR | Gene |
|---|---|---|---|---|
| 12.24 | 1.00 | 0.03 | 0.00E+00 | Tnnc2 |
| 12.17 | 1.00 | 0.02 | 0.00E+00 | Myl1 |
| 12.17 | 0.82 | 0.00 | 0.00E+00 | Myot |
| 12.12 | 1.00 | 0.07 | 0.00E+00 | Acta1 |
| 12.08 | 0.83 | 0.00 | 0.00E+00 | Mybpc2 |
| 12.02 | 1.00 | 0.02 | 0.00E+00 | Ckm |
| 12.01 | 0.80 | 0.00 | 0.00E+00 | Smpx |
| 11.98 | 0.90 | 0.01 | 0.00E+00 | Pvalb |
| 11.97 | 0.99 | 0.07 | 0.00E+00 | Mylpf |
| 11.94 | 0.79 | 0.00 | 0.00E+00 | Trdn |
| 11.83 | 0.90 | 0.00 | 0.00E+00 | Apobec2 |
| 11.82 | 0.89 | 0.01 | 0.00E+00 | Mb |
| 11.81 | 0.90 | 0.01 | 0.00E+00 | Myh1 |
| 11.77 | 0.99 | 0.06 | 0.00E+00 | Tnnt3 |
| 11.75 | 0.99 | 0.01 | 0.00E+00 | Atp2a1 |
| 11.65 | 0.95 | 0.01 | 0.00E+00 | Pgam2 |
| 11.59 | 0.91 | 0.01 | 0.00E+00 | Xirp2 |
| 11.46 | 0.99 | 0.01 | 0.00E+00 | Tcap |
| 11.37 | 0.93 | 0.01 | 0.00E+00 | Myoz1 |
| 11.25 | 0.87 | 0.00 | 0.00E+00 | Cox8b |
| 11.01 | 0.97 | 0.01 | 0.00E+00 | Cox6a2 |
| 10.93 | 0.94 | 0.02 | 0.00E+00 | Actn3 |
| 10.93 | 0.99 | 0.08 | 0.00E+00 | Tnni2 |
| 10.08 | 0.90 | 0.01 | 0.00E+00 | Ttn |
| 9.64 | 0.88 | 0.05 | 0.00E+00 | Ankrd23 |
| 9.60 | 0.96 | 0.01 | 0.00E+00 | Car3 |
| 9.57 | 0.81 | 0.01 | 0.00E+00 | Mybpc1 |
| 9.42 | 0.97 | 0.07 | 0.00E+00 | Eno3 |
| 9.25 | 0.85 | 0.02 | 0.00E+00 | Neb |
| 8.86 | 0.85 | 0.05 | 0.00E+00 | Hspb7 |

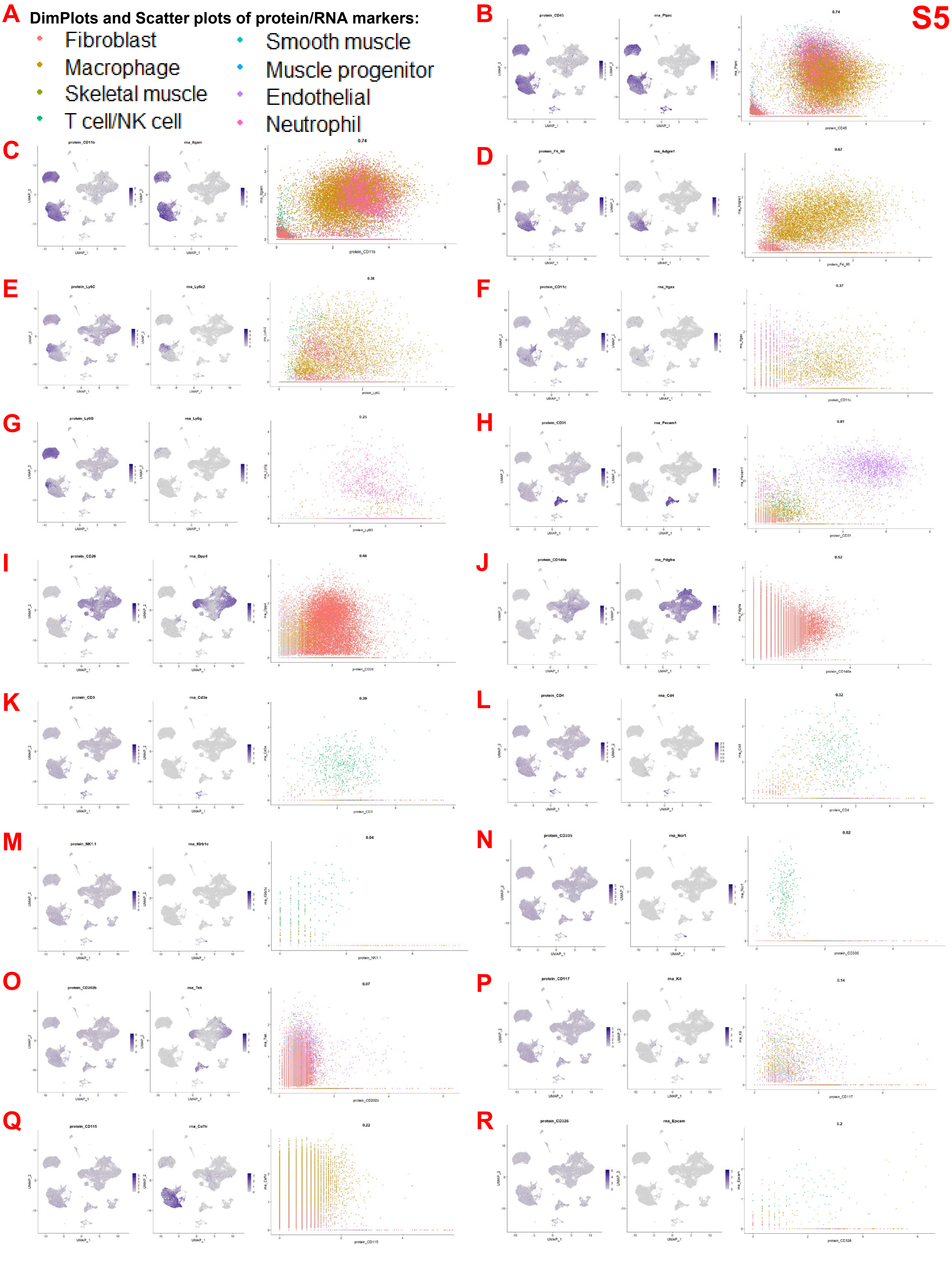
S5
A
DimPlots and Scatter plots of protein/RNA markers:
Fibroblast
Macrophage
Skeletal muscle
T cell/NK cell
Smooth muscle
Muscle progenitor
Endothelial
Neutrophil
B
protein_CD45
rna_Ptprc
0.74
C
protein_CD11b
rna_Itgam
0.74
D
protein_F4_80
rna_Adgre1
0.67
E
protein_Ly6C
rna_Ly6c2
0.38
F
protein_CD11c
rna_Itgax
0.37
G
protein_Ly6G
rna_Ly6g
0.25
H
protein_CD31
rna_Pecam1
0.81
I
protein_CD26
rna_Dpp4
0.66
J
protein_CD140a
rna_Pdgfra
0.52
K
protein_CD3
rna_Cd3e
0.39
L
protein_CD4
rna_Cd4
0.32
M
protein_NK1.1
rna_Klrb1c
0.04
N
protein_CD335
rna_Ncr1
0.02
O
protein_CD202b
rna_Tek
0.07
P
protein_CD117
rna_Kit
0.14
Q
protein_CD115
rna_Csf1r
0.22
R
protein_CD326
rna_Epcam
0.2

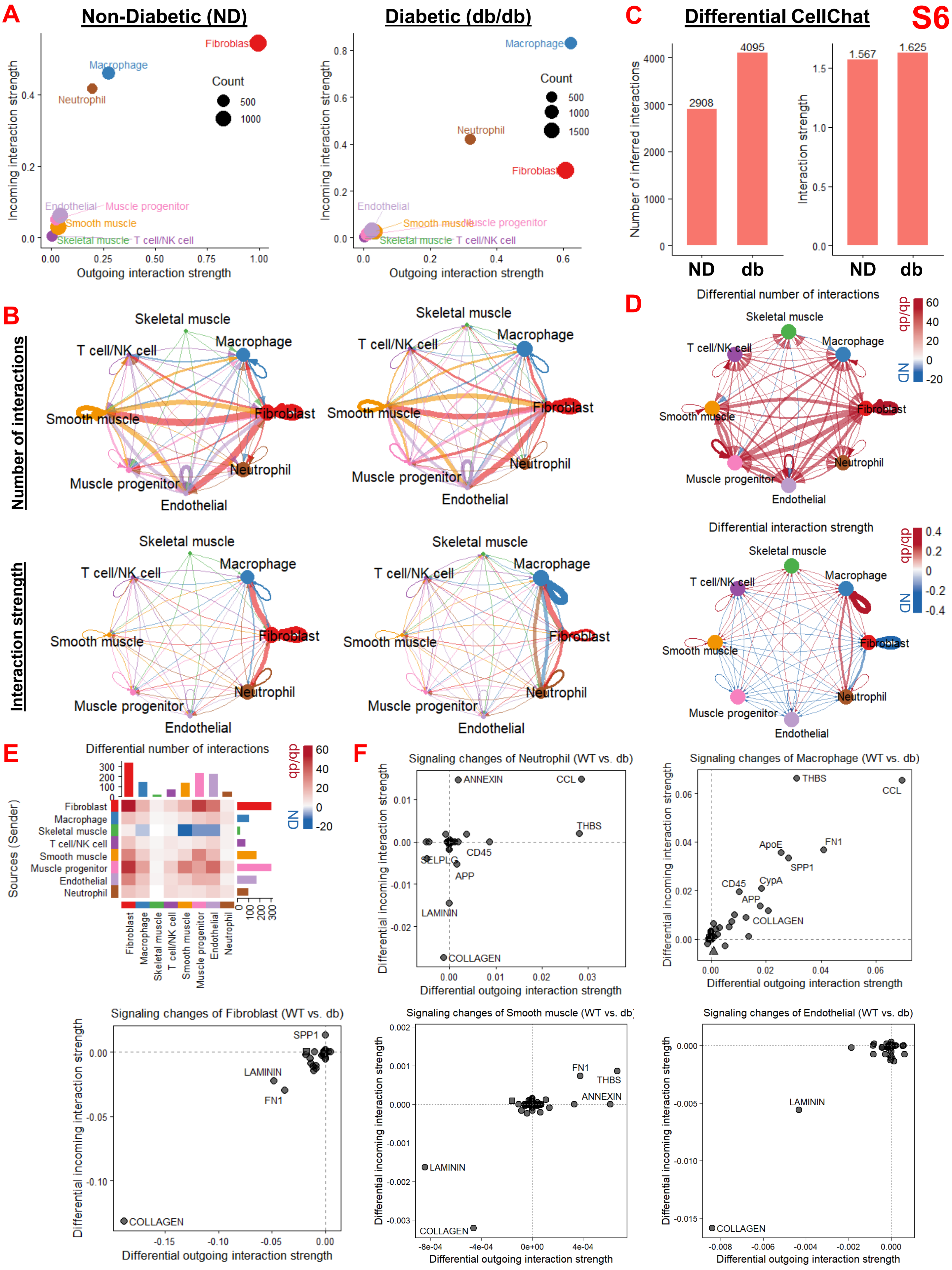

S6
A
Non-Diabetic (ND)
Diabetic (db/db)
Incoming interaction strength
Outgoing interaction strength
Count
Fibroblast
Macrophage
Neutrophil
Endothelial
Muscle progenitor
Smooth muscle
Skeletal muscle
T cell/NK cell
C
Differential CellChat
Number of inferred interactions
Interaction strength
2908
4095
1.567
1.625
ND
db
B
Number of interactions
Interaction strength
D
Differential number of interactions
Differential interaction strength
db/db
ND
E
Sources (Sender)
F
Signaling changes of Neutrophil (WT vs. db)
Signaling changes of Macrophage (WT vs. db)
Signaling changes of Fibroblast (WT vs. db)
Signaling changes of Smooth muscle (WT vs. db)
Signaling changes of Endothelial (WT vs. db)
Differential incoming interaction strength
Differential outgoing interaction strength
ANNEXIN
CCL
THBS
CD45
SELPLG
APP
LAMININ
COLLAGEN
ApoE
FN1
SPP1
CypA

S7

**A** Elbow plots:

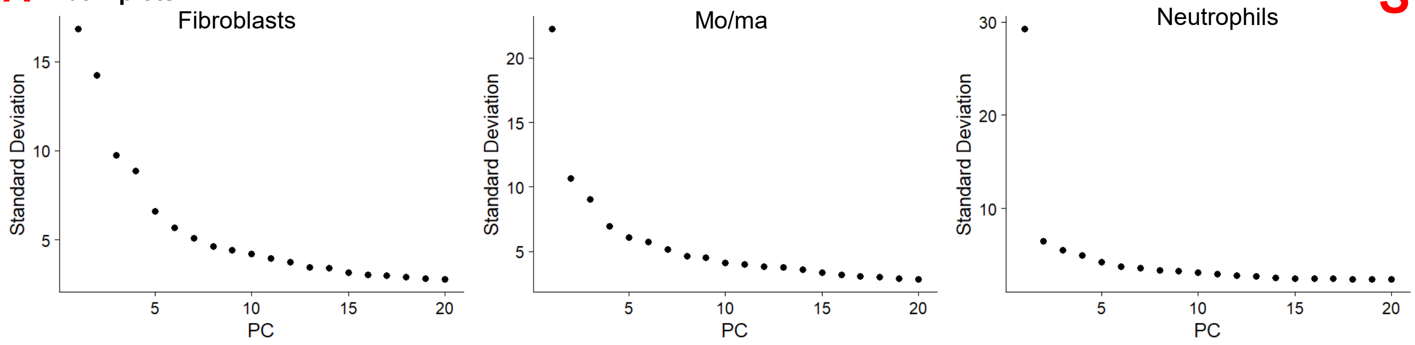


**B** Fibroblast sub-types DEG analysis:

| **fδ :** *Proliferative FB* | | |
|---|---|---|
| **$Log_2FC$** | **% pos** | **Gene** |
| 3.73 | 0.98 | *Cks2* |
| 3.47 | 0.99 | *Stmn1* |
| 3.36 | 0.99 | *Hmgb2* |
| 3.13 | 0.96 | *Smc2* |
| 2.46 | 0.94 | *Tyms* |
| 2.36 | 0.98 | *Smc4* |
| 2.36 | 0.99 | *Tubb4b* |
| 2.34 | 0.91 | *H2afx* |
| 2.20 | 1.00 | *Tuba1b* |
| 2.20 | 1.00 | *H2afz* |
| 2.13 | 0.88 | *Plk4* |
| 2.11 | 0.96 | *Tuba1c* |
| 2.10 | 0.80 | *Lig1* |
| 2.10 | 1.00 | *Ube2s* |
| 2.05 | 0.99 | *Selenoh* |
| 2.04 | 0.96 | *Rangap1* |
| 2.01 | 1.00 | *Ran* |
| 1.98 | 0.99 | *Cks1b* |
| 1.93 | 0.97 | *Rrm1* |
| 1.92 | 0.96 | *Lmnb1* |

**Pathway enrichment:**

| **Reactome** FDR < 0.05 |
|---|
| Cell Cycle, Mitotic |
| Cell Cycle |
| M Phase |
| Nuclear Envelope (NE) Reassembly |

**C** Macrophage sub-types DEG analysis:

| **mδ :** *Proliferative Mo/Ma* | | |
|---|---|---|
| **$Log_2FC$** | **% pos** | **Gene** |
| 4.24 | 0.60 | *Ctsk* |
| 4.16 | 0.75 | *Acp5* |
| 2.79 | 0.77 | *Nfatc1* |
| 2.30 | 0.87 | *Smc4* |
| 2.29 | 0.89 | *Tubb4b* |
| 2.16 | 0.99 | *Ranbp1* |
| 2.15 | 0.92 | *Sms* |
| 2.14 | 0.90 | *Dtymk* |
| 2.14 | 0.96 | *Hmgb2* |
| 2.13 | 0.92 | *Hmgn1* |
| 2.11 | 0.86 | *Ccdc34* |
| 2.10 | 0.87 | *Lpl* |
| 2.08 | 0.98 | *Pa2g4* |
| 2.06 | 1.00 | *Tubb5* |
| 2.04 | 0.94 | *Nrp2* |
| 2.04 | 0.94 | *C1qbp* |
| 2.01 | 0.93 | *Nsd2* |
| 1.99 | 0.83 | *Dctpp1* |
| 1.98 | 1.00 | *Ran* |
| 1.95 | 0.99 | *Nucks1* |

**Pathway enrichment:**

| **Reactome** FDR < 0.05 |
|---|
| Nuclear Envelope (NE) Reassembly |
| Cell Cycle |
| Metabolism of Nucleotides |
| M Phase |

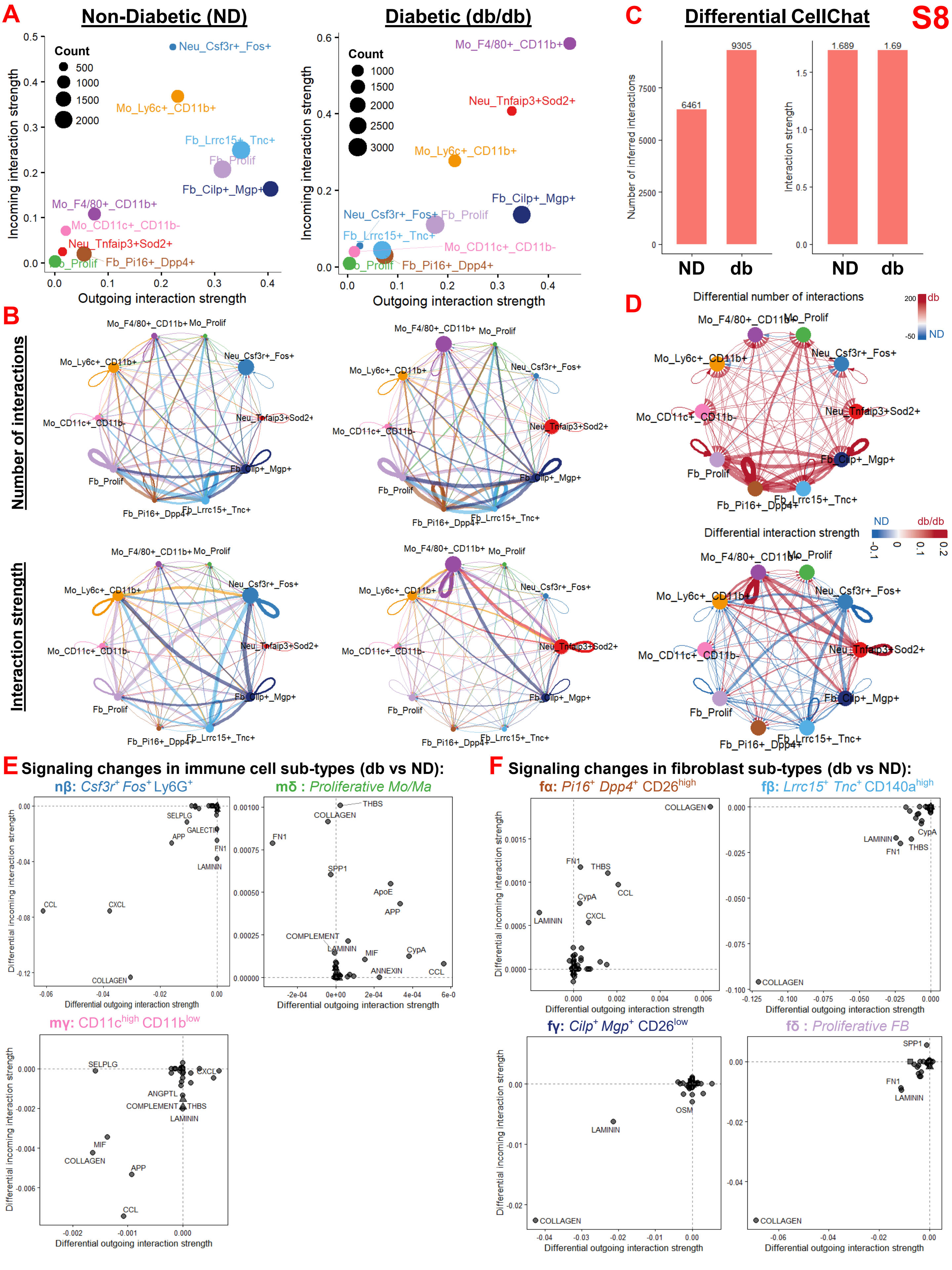
S8
A
Non-Diabetic (ND)
Diabetic (db/db)
Count
Incoming interaction strength
Outgoing interaction strength
Neu_Csf3r+_Fos+
Mo_Ly6c+_CD11b+
Fb_Lrrc15+_Tnc+
Fb_Prolif
Fb_Cilp+_Mgp+
Mo_F4/80+_CD11b+
Mo_CD11c+_CD11b-
Neu_Tnfaip3+Sod2+
Fb_Pi16+_Dpp4+
Mo_Prolif
C
Differential CellChat
Number of inferred interactions
Interaction strength
6461
9305
1.689
1.69
ND
db
B
Number of interactions
Interaction strength
D
Differential number of interactions
Differential interaction strength
ND
db/db
E
Signaling changes in immune cell sub-types (db vs ND):
nβ: Csf3r+ Fos+ Ly6G+
mδ : Proliferative Mo/Ma
mγ: CD11c^high CD11b^low
F
Signaling changes in fibroblast sub-types (db vs ND):
fα: Pi16+ Dpp4+ CD26^high
fβ: Lrrc15+ Tnc+ CD140a^high
fγ: Cilp+ Mgp+ CD26^low
fδ : Proliferative FB
Differential incoming interaction strength
Differential outgoing interaction strength
SELPLG
GALECTIN
APP
FN1
LAMININ
CCL
CXCL
COLLAGEN
THBS
SPP1
ApoE
COMPLEMENT
MIF
CypA
ANNEXIN
ANGPTL
OSM

**A** Gating strategy for flow cytometry:

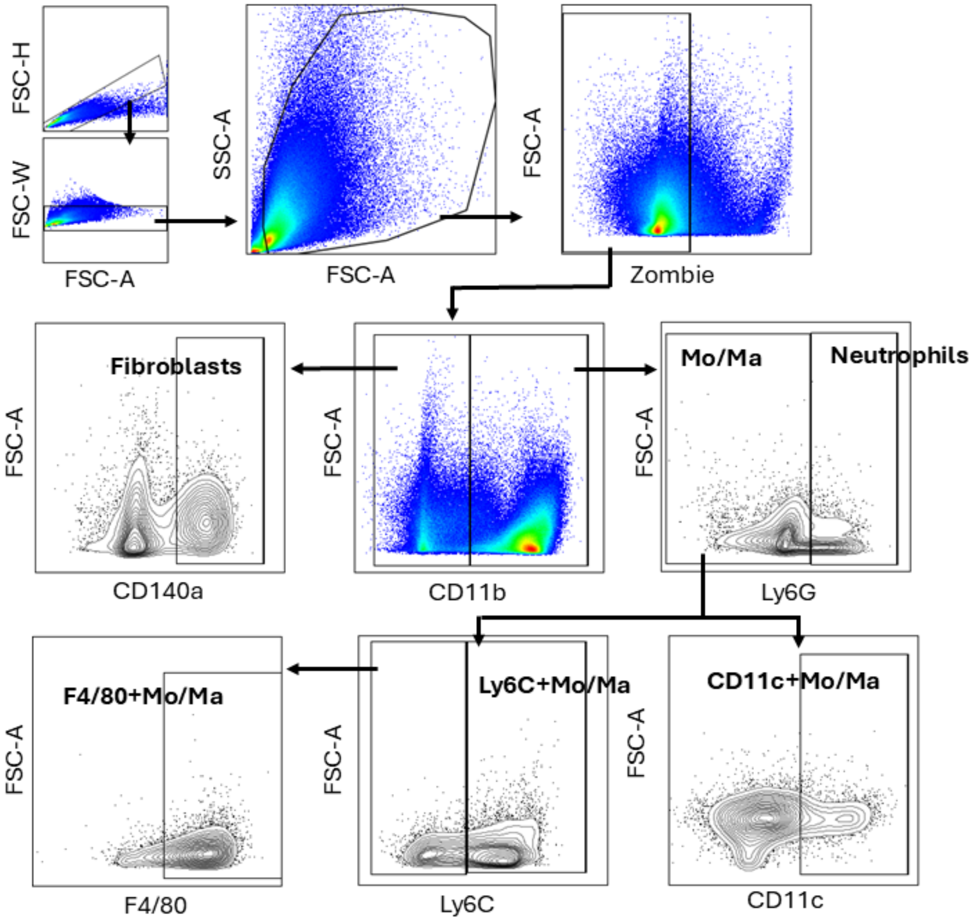


**B** Gating for CD44 expression in ND and db/db wounds:

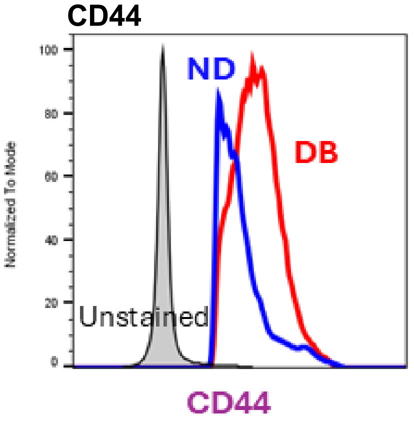


**E** Gating for CD44 and ICAM-1 expression in db/db wounds treated with CD44-ab or IgG:

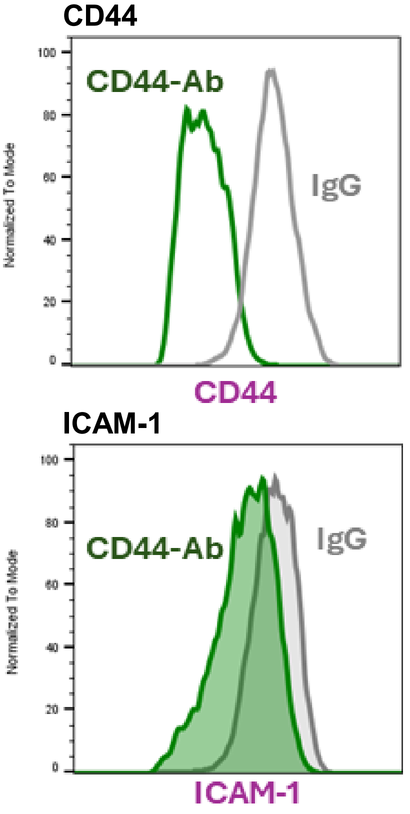


Additional gene expression analyses:

Treatment groups: ND, ND + CD44-ab, db/db, db/db + CD44-ab

**C**

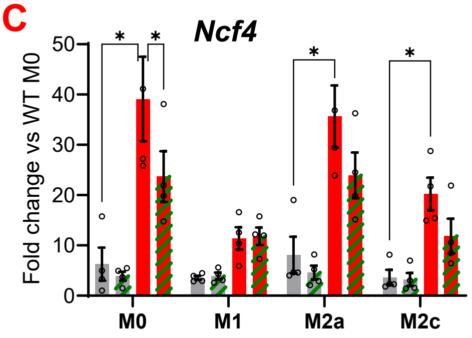


| Source | % variation | P value |
|---|---|---|
| Interaction | 11.20 | 0.020258 |
| Polarization | 13.50 | 0.000078 |
| Treatment | 51.29 | <0.000001 |

**D**

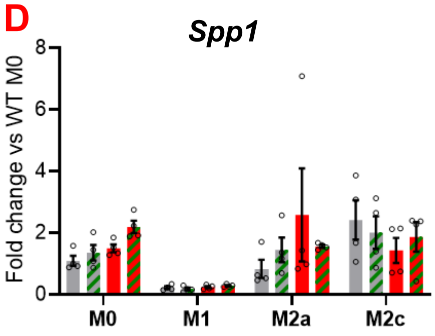


| Source | % variation | P value |
|---|---|---|
| Interaction | 11.81 | 0.339938 |
| Polarization | 32.52 | 0.000045 |
| Treatment | 1.462 | 0.731433 |

**F** Wound closure in db/db wounds treated with CD44-ab or IgG:

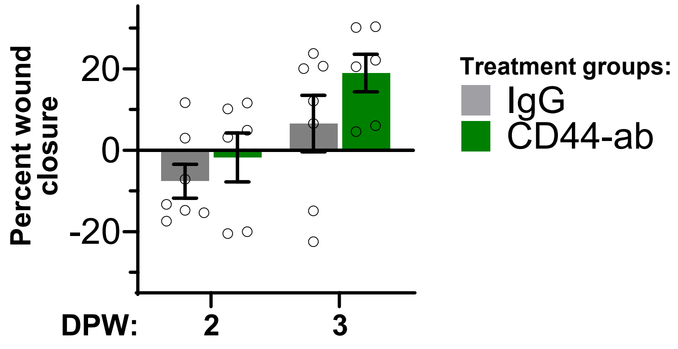

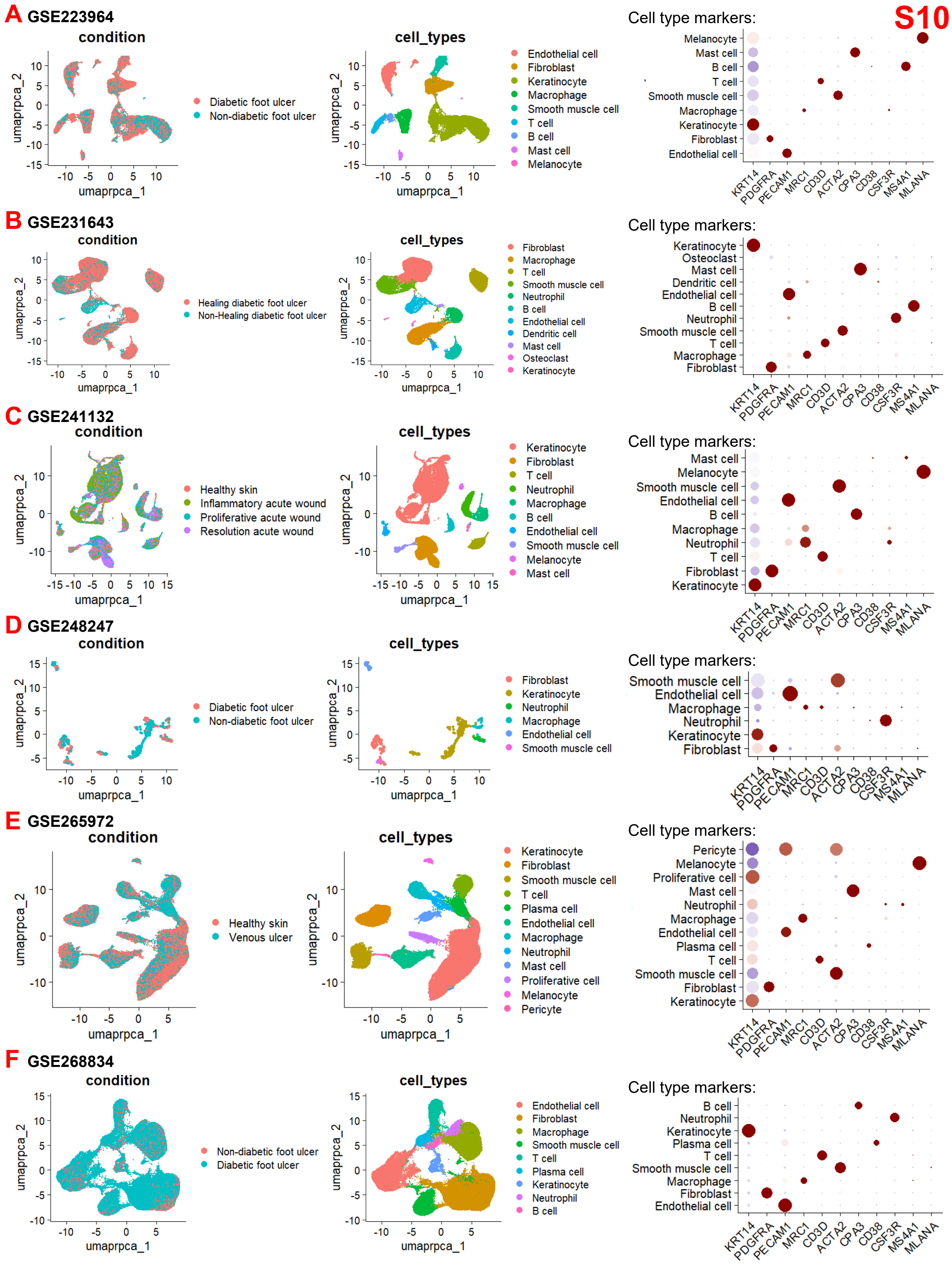
S10
A GSE223964
condition
umaprpca_2
umaprpca_1
Diabetic foot ulcer
Non-diabetic foot ulcer
cell_types
Endothelial cell
Fibroblast
Keratinocyte
Macrophage
Smooth muscle cell
T cell
B cell
Mast cell
Melanocyte
Cell type markers:
Melanocyte
Mast cell
B cell
T cell
Smooth muscle cell
Macrophage
Keratinocyte
Fibroblast
Endothelial cell
KRT14
PDGFRA
PECAM1
MRC1
CD3D
ACTA2
CPA3
CD38
CSF3R
MS4A1
MLANA
B GSE231643
condition
Healing diabetic foot ulcer
Non-Healing diabetic foot ulcer
cell_types
Fibroblast
Macrophage
T cell
Smooth muscle cell
Neutrophil
B cell
Endothelial cell
Dendritic cell
Mast cell
Osteoclast
Keratinocyte
Cell type markers:
C GSE241132
condition
Healthy skin
Inflammatory acute wound
Proliferative acute wound
Resolution acute wound
cell_types
Keratinocyte
Fibroblast
T cell
Neutrophil
Macrophage
B cell
Endothelial cell
Smooth muscle cell
Melanocyte
Mast cell
Cell type markers:
D GSE248247
condition
Diabetic foot ulcer
Non-diabetic foot ulcer
cell_types
Fibroblast
Keratinocyte
Neutrophil
Macrophage
Endothelial cell
Smooth muscle cell
Cell type markers:
E GSE265972
condition
Healthy skin
Venous ulcer
cell_types
Keratinocyte
Fibroblast
Smooth muscle cell
T cell
Plasma cell
Endothelial cell
Macrophage
Neutrophil
Mast cell
Proliferative cell
Melanocyte
Pericyte
Cell type markers:
F GSE268834
condition
Non-diabetic foot ulcer
Diabetic foot ulcer
cell_types
Endothelial cell
Fibroblast
Macrophage
Smooth muscle cell
T cell
Plasma cell
Keratinocyte
Neutrophil
B cell
Cell type markers:

S11

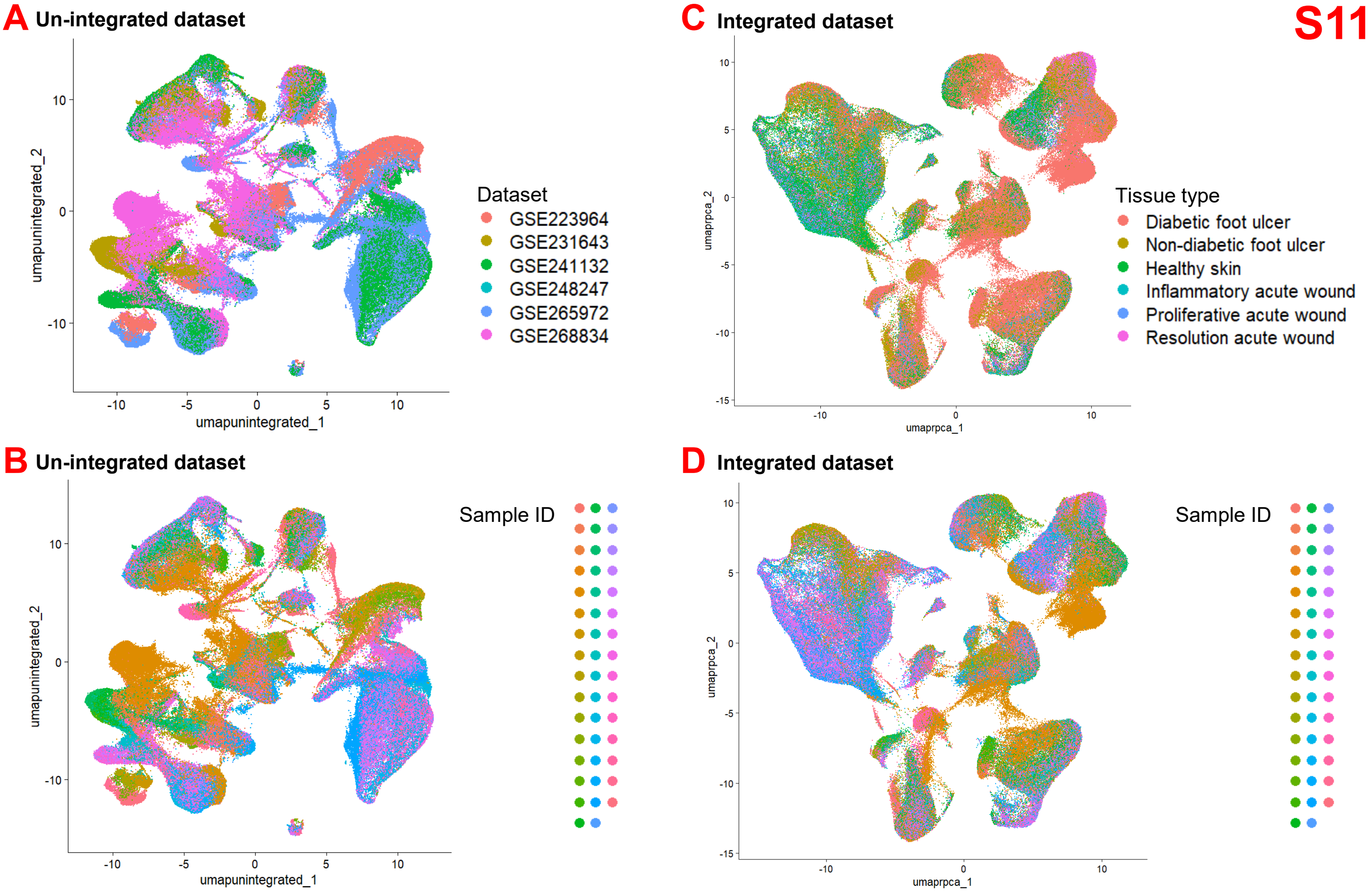


#### E Cell types per sample in final integrated dataset

| Dataset | GSE 223964 | | | | | | | | GSE 231643 | | | | | | | | GSE 248247 | | GSE 241132 | | | GSE 265972 | | | | | | | | | GSE 268834 | | | | | | | |
|---|---|---|---|---|---|---|---|---|---|---|---|---|---|---|---|---|---|---|---|---|---|---|---|---|---|---|---|---|---|---|---|---|---|---|---|---|---|---|
| Tissue type | NDFU | NDFU | NDFU | DFU | DFU | DFU | DFU | DFU | DFU | DFU | DFU | DFU | DFU | DFU | DFU | DFU | NDFU | DFU | Skin | Skin | Skin | Skin | Skin | Skin | Skin | Skin | NDFU | NDFU | NDFU | NDFU | NDFU | NDFU | NDFU | NDFU | DFU | DFU | DFU | DFU |
| Keratinocyte | 2062 | 485 | 1713 | 4973 | 186 | 548 | 113 | 169 | 10 | 4 | 1 | 27 | 3 | 37 | 58 | 27 | 152 | 28 | 1720 | 1571 | 1789 | 2815 | 2739 | 3717 | 2660 | 6891 | 509 | 3750 | 1903 | 2003 | 709 | 128 | 18 | 238 | 130 | 134 | 375 | 1050 |
| Fibroblast | 45 | 499 | 318 | 1673 | 136 | 57 | 1024 | 320 | 821 | 1653 | 1346 | 1736 | 412 | 462 | 195 | 451 | 38 | 28 | 703 | 512 | 882 | 936 | 660 | 738 | 1061 | 433 | 1016 | 321 | 674 | 2006 | 601 | 2307 | 28 | 439 | 2537 | 603 | 15708 | 3177 |
| Endothelial | 679 | 303 | 269 | 1124 | 684 | 35 | 879 | 304 | 229 | 165 | 108 | 224 | 44 | 224 | 65 | 959 | 30 | 13 | 177 | 136 | 60 | 180 | 489 | 329 | 406 | 1061 | 946 | 1613 | 238 | 316 | 782 | 667 | 871 | 2731 | 2273 | 279 | 5558 | 1396 |
| Macrophage | 37 | 98 | 386 | 297 | 128 | 41 | 290 | 624 | 369 | 1495 | 510 | 874 | 236 | 101 | 1036 | 322 | 9 | 9 | 567 | 378 | 457 | 425 | 626 | 266 | 270 | 248 | 1029 | 208 | 413 | 450 | 436 | 1713 | 38 | 546 | 1446 | 296 | 4955 | 568 |
| SMC | 17 | 170 | 105 | 229 | 136 | 23 | 777 | 388 | 202 | 327 | 193 | 853 | 506 | 325 | 391 | 686 | 17 | 19 | 204 | 113 | 114 | 276 | 549 | 839 | 496 | 549 | 768 | 651 | 243 | 207 | 231 | 243 | 171 | 2880 | 624 | 189 | 1184 | 583 |
| Neutrophil | 5 | 1 | 183 | 2 | 1 | 3 | 11 | 95 | 2177 | 22 | 16 | 10 | 5 | 19 | 106 | 2 | 8 | 34 | 0 | 0 | 0 | 35 | 5 | 1 | 0 | 13 | 185 | 3 | 287 | 18 | 52 | 91 | 0 | 129 | 420 | 889 | 12 | 26 |
| Mast cell | 3 | 2 | 10 | 0 | 3 | 0 | 4 | 497 | 945 | 29 | 16 | 880 | 66 | 0 | 104 | 85 | 1 | 1 | 1 | 3 | 1 | 10 | 36 | 3 | 13 | 15 | 221 | 119 | 9 | 27 | 19 | 29 | 10 | 105 | 6 | 12 | 133 | 7 |
| T cell | 59 | 46 | 375 | 149 | 326 | 155 | 49 | 407 | 783 | 918 | 88 | 1083 | 147 | 34 | 414 | 301 | 9 | 3 | 364 | 206 | 340 | 446 | 301 | 78 | 216 | 75 | 1789 | 694 | 328 | 1166 | 203 | 658 | 64 | 973 | 277 | 429 | 1543 | 1323 |
| B cell | 6 | 81 | 61 | 14 | 63 | 11 | 326 | 3 | 6 | 6 | 10 | 43 | 10 | 76 | 3 | 42 | 1 | 1 | 321 | 213 | 321 | 148 | 116 | 210 | 26 | 156 | 160 | 696 | 57 | 356 | 146 | 146 | 28 | 109 | 10 | 31 | 427 | 143 |
| Plasma cell | 0 | 0 | 0 | 0 | 2 | 0 | 0 | 0 | 12 | 3 | 1 | 102 | 3 | 0 | 0 | 5 | 0 | 2 | 0 | 1 | 1 | 0 | 8 | 0 | 1 | 0 | 1988 | 3 | 1 | 1 | 41 | 0 | 1 | 832 | 0 | 1 | 2258 | 88 |
| Melanocyte | 35 | 0 | 28 | 2 | 0 | 2 | 0 | 0 | 0 | 2 | 0 | 0 | 0 | 0 | 0 | 0 | 0 | 0 | 27 | 32 | 56 | 53 | 11 | 82 | 74 | 154 | 8 | 241 | 97 | 0 | 0 | 2 | 7 | 3 | 0 | 30 | 3 | 72 |

#### F Cell sub-types per sample in final dataset

| Dataset | GSE 223964 | | | | | | | | GSE 231643 | | | | | | | | GSE 248247 | | GSE 241132 | | | GSE 265972 | | | | | | | | | GSE 268834 | | | | | | | |
|---|---|---|---|---|---|---|---|---|---|---|---|---|---|---|---|---|---|---|---|---|---|---|---|---|---|---|---|---|---|---|---|---|---|---|---|---|---|---|
| Tissue type | NDFU | NDFU | NDFU | DFU | DFU | DFU | DFU | DFU | DFU | DFU | DFU | DFU | DFU | DFU | DFU | DFU | NDFU | DFU | Skin | Skin | Skin | Skin | Skin | Skin | Skin | Skin | NDFU | NDFU | NDFU | NDFU | NDFU | NDFU | NDFU | NDFU | DFU | DFU | DFU | DFU |
| *TNFAIP3*$^{+}$ *SOD2*$^{+}$ | 3 | 1 | 144 | 1 | 1 | 3 | 9 | 80 | 839 | 5 | 11 | 1 | 2 | 6 | 54 | 0 | 4 | 19 | 0 | 0 | 0 | 9 | 3 | 0 | 0 | 1 | 176 | 0 | 281 | 17 | 45 | 69 | 0 | 101 | 379 | 767 | 6 | 18 |
| *CSF3R*$^{+}$ *FOS*$^{+}$ | 2 | 0 | 39 | 1 | 0 | 0 | 2 | 15 | 1338 | 17 | 5 | 9 | 3 | 13 | 52 | 2 | 4 | 15 | 0 | 0 | 0 | 26 | 2 | 1 | 0 | 12 | 9 | 3 | 6 | 1 | 7 | 22 | 0 | 28 | 41 | 122 | 6 | 8 |
| *CD163*$^{+}$ *MRC1*$^{+}$ | 17 | 61 | 209 | 220 | 111 | 28 | 154 | 383 | 238 | 879 | 236 | 556 | 106 | 69 | 401 | 162 | 8 | 7 | 250 | 140 | 207 | 128 | 409 | 151 | 118 | 145 | 850 | 137 | 170 | 357 | 274 | 1196 | 19 | 372 | 669 | 84 | 4776 | 380 |
| *VCAN*$^{+}$ *IL1B*$^{+}$ | 19 | 32 | 173 | 67 | 12 | 12 | 89 | 219 | 91 | 467 | 207 | 191 | 98 | 20 | 554 | 61 | 1 | 1 | 36 | 29 | 26 | 158 | 45 | 24 | 50 | 12 | 113 | 3 | 219 | 53 | 108 | 461 | 0 | 144 | 711 | 209 | 109 | 107 |
| *CD86*$^{+}$ *CCR7*$^{+}$ | 1 | 5 | 4 | 10 | 5 | 1 | 47 | 22 | 40 | 149 | 67 | 127 | 32 | 12 | 81 | 99 | 0 | 1 | 281 | 209 | 224 | 139 | 172 | 91 | 102 | 91 | 66 | 68 | 24 | 40 | 54 | 56 | 19 | 30 | 66 | 3 | 70 | 81 |
| *PRG4*$^{+}$ *DPP4*$^{+}$ | 2 | 7 | 1 | 26 | 2 | 0 | 1 | 0 | 4 | 2 | 22 | 30 | 1 | 7 | 0 | 40 | 15 | 11 | 12 | 17 | 43 | 66 | 31 | 94 | 221 | 173 | 29 | 84 | 11 | 10 | 32 | 20 | 4 | 77 | 112 | 53 | 10519 | 471 |
| *LRRC15*$^{+}$ *TNC*$^{+}$ | 43 | 489 | 317 | 1628 | 125 | 57 | 1023 | 318 | 814 | 1642 | 1274 | 1685 | 409 | 454 | 195 | 269 | 10 | 17 | 17 | 38 | 59 | 134 | 197 | 83 | 55 | 24 | 670 | 17 | 645 | 1955 | 305 | 1868 | 19 | 230 | 2109 | 142 | 4693 | 11 |
| *CILP*$^{+}$ *MGP*$^{+}$ | 0 | 3 | 0 | 19 | 9 | 0 | 0 | 2 | 3 | 9 | 50 | 21 | 2 | 1 | 0 | 142 | 13 | 0 | 674 | 457 | 780 | 736 | 432 | 561 | 785 | 236 | 317 | 220 | 18 | 41 | 264 | 419 | 5 | 132 | 316 | 408 | 496 | 2695 |

S12

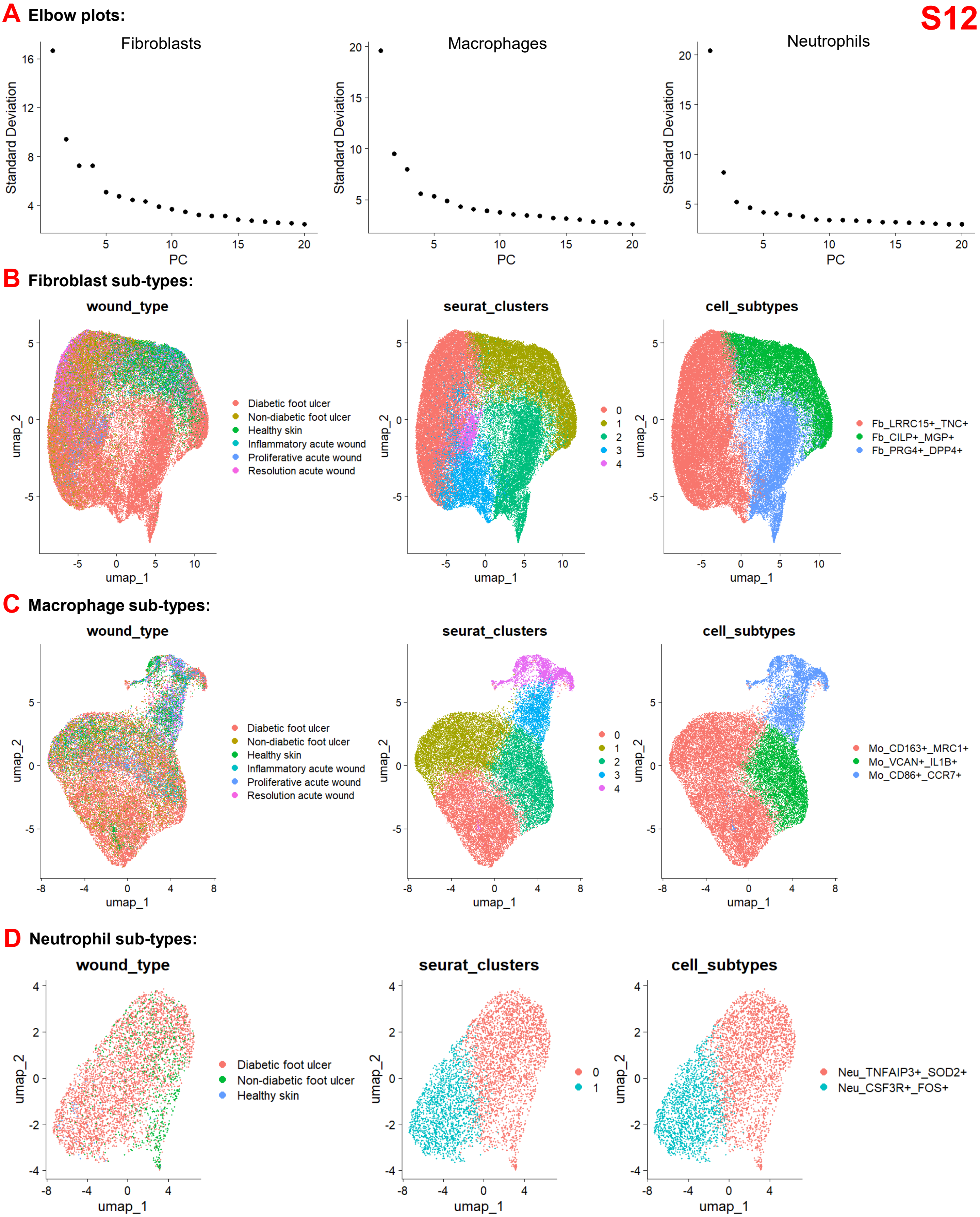
A Elbow plots:
Fibroblasts
Macrophages
Neutrophils
Standard Deviation
PC
B Fibroblast sub-types:
wound_type
seurat_clusters
cell_subtypes
umap_1
umap_2
Diabetic foot ulcer
Non-diabetic foot ulcer
Healthy skin
Inflammatory acute wound
Proliferative acute wound
Resolution acute wound
Fb_LRRC15+_TNC+
Fb_CILP+_MGP+
Fb_PRG4+_DPP4+
C Macrophage sub-types:
Mo_CD163+_MRC1+
Mo_VCAN+_IL1B+
Mo_CD86+_CCR7+
D Neutrophil sub-types:
Neu_TNFAIP3+_SOD2+
Neu_CSF3R+_FOS+

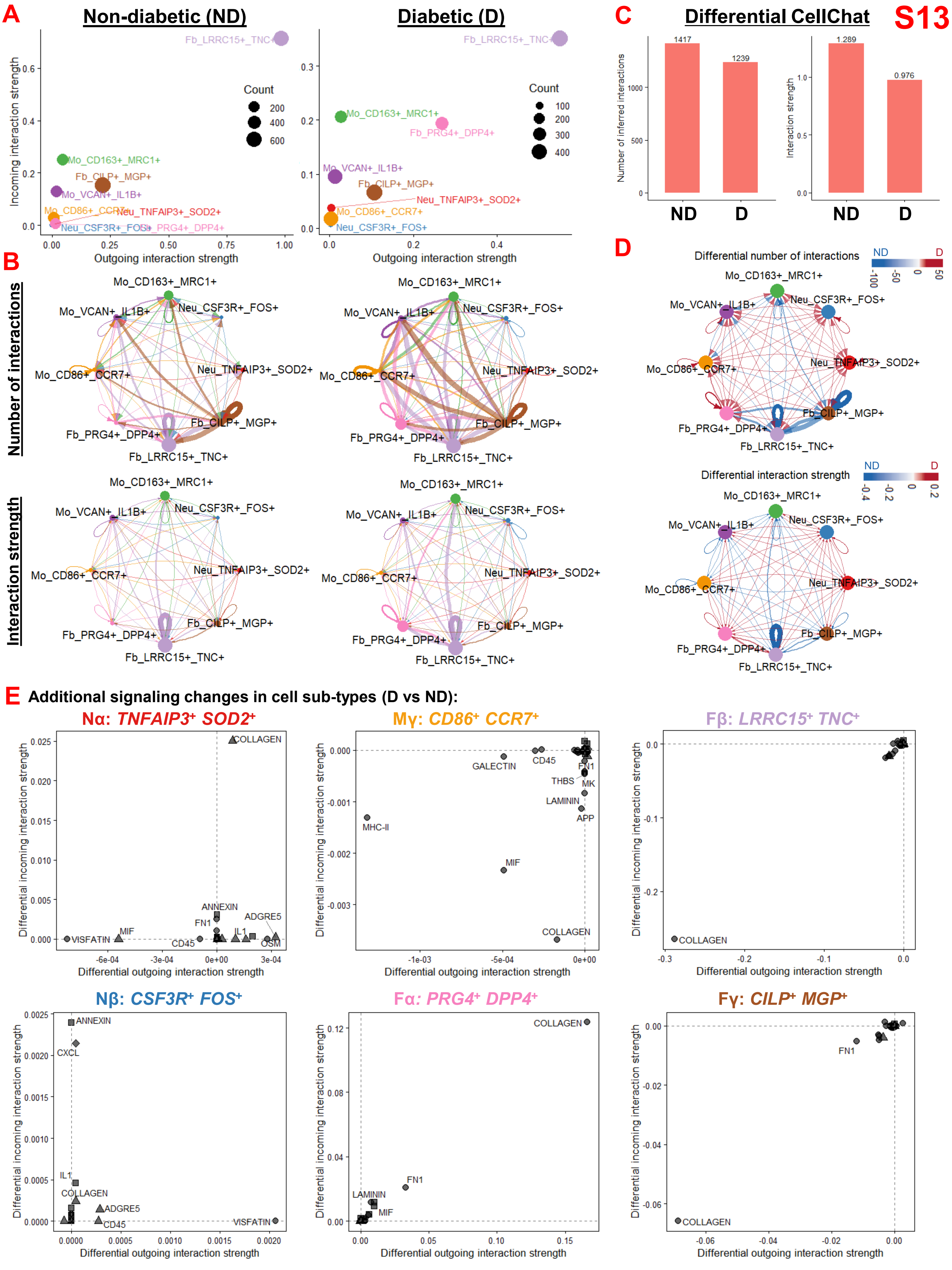
S13
A
Non-diabetic (ND)
Diabetic (D)
Incoming interaction strength
Outgoing interaction strength
Count
200
400
600
100
300
Fb_LRRC15+_TNC+
Mo_CD163+_MRC1+
Fb_CILP+_MGP+
Mo_VCAN+_IL1B+
Mo_CD86+_CCR7+
Neu_TNFAIP3+_SOD2+
Neu_CSF3R+_FOS+
Fb_PRG4+_DPP4+
C
Differential CellChat
Number of inferred interactions
Interaction strength
1417
1239
1.289
0.976
ND
D
B
Number of interactions
Interaction strength
D
Differential number of interactions
Differential interaction strength
E
Additional signaling changes in cell sub-types (D vs ND):
Nα: TNFAIP3+ SOD2+
Mγ: CD86+ CCR7+
Fβ: LRRC15+ TNC+
Nβ: CSF3R+ FOS+
Fα: PRG4+ DPP4+
Fγ: CILP+ MGP+
Differential incoming interaction strength
Differential outgoing interaction strength
COLLAGEN
ANNEXIN
ADGRE5
FN1
IL1
VISFATIN
MIF
CD45
OSM
GALECTIN
THBS
MK
LAMININ
APP
MHC-II
CXCL
